# Single-molecule detection of transient dimerization of opioid receptors 1: Homodimers’ effect on signaling and internalization

**DOI:** 10.1101/2024.07.25.605080

**Authors:** Peng Zhou, Taka A. Tsunoyama, Rinshi S. Kasai, Koichiro M. Hirosawa, Ziya Kalay, Amine Aladag, Takahiro K. Fujiwara, Simone Pigolotti, Akihiro Kusumi

## Abstract

Opioid receptors (ORs) are critical for endogenous and synthetic analgesics. Their homodimerization is considered important for their pharmacological diversities, but whether they form homodimers remains controversial. Here, we established that the three classical ORs, μ-, κ-, and δ-ORs (MOR, KOR, and DOR, respectively) undergo repeated transient (120-180 ms) homodimerizations every few seconds. This was done by using single-molecule imaging and developing theories for analyzing single-molecule colocalization data, which provide the key parameters, homodimer-monomer dissociation equilibrium constants and rate constants. Their 9-26 amino-acid C-terminal cytoplasmic domains, without sequence similarities, are involved in specific homodimerization, whereas the transmembrane domains provide less specific affinities. Using the membrane-permeable peptides mimicking the C-terminal homodimerization sequences which block homodimerizations, functions of monomers and homodimers were dissected. KOR and DOR homodimers, but not MOR homodimers, activate downstream G-proteins differently from monomers upon agonist addition, without influencing OR internalization. These findings could guide strategies to enhance OR-based analgesia.

## Introduction

Three classical opioid receptors (ORs), μ-, κ-, and δ-ORs (MOR, KOR, and DOR, respectively), are distributed across the central and peripheral nervous systems and play important roles in regulating pain perception, hedonic homeostasis, mood, and well-being.^1, 2^ ORs are prototypical class A G-protein-coupled receptors (GPCRs), and serve as key receptors for a variety of endogenous and synthetic analgesics. These OR subtypes play critical roles in pain therapeutics, and at the same time, in the development of tolerance to and dependence on analgesics, which are likely associated with activations of the inhibitory G-protein and β-arrestin pathways in complex manners.^2, 3, 4^

The diversity of OR conformations and their interactions with other proteins not only drives various cellular physiological responses but also contributes to a spectrum of brain circuit responses, both beneficial and adverse.^4, 5, 6, 7^ Hence, a profound understanding of the mechanisms by which ORs’ distinct downstream responses are induced is essential for the development of analgesic drugs with minimized side effects. In addition, receptor dimerization and clustering are often key steps for triggering the downstream signaling cascades in other receptor signaling.^8^

The pharmacological diversities of ORs might be enhanced by their propensity to form homodimers^9, 10^ and heterodimers.^11, 12, 13^ Agonist-induced homodimer formation significantly influences the balance of MOR’s downstream G-protein and β-arrestin pathways.^14^ Other studies suggested that dimerization, both homo and hetero, might modulate or be modulated by biased agonism, highlighting its significance in receptor signaling (for example, see Refs^15,16^ for homodimers and Refs^17, 18^ for heterodimers). For the β2-adrenergic receptor, another class A GPCR, homodimers are responsible for generating the basal receptor signal in the absence of agonists, a key common characteristic of GPCRs.^19^ Furthermore, OR heterodimerization is important for their cooperative functions, as supported by functional and pharmacological studies.^17, 20^ Therefore, understanding both the homo- and hetero-dimerizations of ORs, as well as those of other class-A GPCRs, is fundamental to deciphering their functional regulation mechanisms.^16, 21, 22, 23, 24, 25^

Despite such extensive research, the very existence of homo- and hetero-dimers of ORs, particularly at physiological expression levels, remains controversial.^26^ Previous detection of homodimers of DOR,^9, 27, 28^ MOR,^27, 28^ and KOR,^13, 28^ and heterodimers of all OR pairs^28^ contrasts with recent single-molecule imaging studies. These studies, performed at physiological expression levels in the plasma membrane (PM) of living cells, in contrast to the results obtained under the conditions of overexpression in cells and very high concentrations in vitro, which are often necessary for the use of bulk observation methods, showed that MOR,^29, 30^ DOR,^30^ and KOR^31^ are predominantly monomeric. Therefore, they concluded that previously observed dimerization and its functional and pharmacological consequences would be artifacts due to the employment of very high OR concentrations. Meanwhile, other single-molecule investigations detected the nanodomains of KOR and MOR^32^ as well as the transient MOR dimerization^33^ or that only after the agonist DAMGO binding, but not morphine.^14^

To address the overexpression issue and resolve these controversies, determining the dimer-monomer equilibrium constant K_D_ is imperative, as it is unaffected by the expression levels. In addition, since several GPCR homodimers are apparently metastable with lifetimes on the order of 0.1-1 s,^19, 21, 33, 34, 35, 36, 37^ estimating the dimer dissociation and association rate constants, k_off_ and k_on_, respectively, is essential for establishing the comprehensive description of the dynamic dimer-monomer equilibrium of ORs. Such insights would enable us to unequivocally determine whether ORs form homo- and hetero-dimers and to what extent under various expression conditions, including both the lower and higher regions of physiological expression levels in different cells in various brain regions. Furthermore, identifying specific binding sites for homo- and hetero-dimerizations, given the high amino acid similarities among the three classical ORs, would be interesting from the protein-interaction viewpoint and highly beneficial for developing reagents that modify dimerizations.

In the present research, we addressed these issues, using advanced single-molecule imaging, tracking, and analysis. The single-molecule approach was selected for its ability to directly visualize receptor molecules’ dimerization and dissociation in living cells. To extract the dimer-monomer equilibrium constant, K_D_, and rate constants, k_off_ and k_on_, from single-molecule imaging data, we developed two novel theories: one to evaluate k_off_ from the distribution of single-molecule colocalization durations (instead of intuitive methods employed previously), and another to estimates K_D_ (k_on_ can be calculated as k_off_/K_D_).

We present our findings in two back-to-back papers. This paper focuses on homodimers, while the companion paper addresses heterodimers.^38^ In this paper, we report that all three ORs engage in continual, repeated, transient homodimerizations at 37°C, with K_D_s of 5.87 ± 0.56, 15.24 ± 1.54, and 16.42 ± 0.53 copies/µm^2^ for KOR, MOR, and DOR, respectively. The homodimer lifetimes are in the range of 120-180 ms (dimer dissociation rate constants in the range of 6.71 to 8.47 s^-1^), unequivocally demonstrating that even at low physiological expression levels, the three ORs form transient homodimers. Furthermore, we identified that the ≈9-26 amino acid sequences in the cytoplasmic C-terminal domains, without sequence similarities, are involved in the distinct homodimerizations of the three ORs, in line with the previous DOR homodimerization data,^9^ whereas the transmembrane domains might promote both homo- and heterodimerizations in a less specific manner, in partial agreement with previous results.^21, 39, 40, 41, 42^

Building on these amino-acid sequences, we developed peptides that suppress OR homodimerization within live cells. This has allowed us to distinguish the functions of OR dimers from monomers, which could not be done before. OR monomerization by these peptides enhances and diminishes agonist-induced signals downstream from G proteins for KOR and DOR, respectively, without influencing agonist-induced internalization. Meanwhile, monomerization had no discernible effect on MOR signaling or internalization. These findings may inform novel strategies in developing more effective and safer pain treatments.

## Results

### All three ORs continually interconvert between transient homodimers and monomers: analysis by colocalization index based on the pair cross-correlation function (PCCF)

ORs conjugated with the SNAPf tag protein at their N-termini (SNAPf-ORs) were expressed in the PM of CHO-K1 cells, which do not express ORs.^43^ These SNAPf-tagged molecules were functional (Supplementary Fig. 1a, b) and labeled with the fluorescent membrane-impermeable SNAP ligands, SNAP-Surface 549 and SNAP-CF660R, with 60% and 61% efficiencies, respectively (Supplementary Fig. 1c-e). For the experiments examining OR homodimerization, we simultaneously labeled a SNAPf-OR (for example, SNAPf-KOR) with both SNAP-Surface 549 and SNAP-CF660R so that the number densities of the two probes on the PM were about the same, with spot densities of 0.5 ± 0.25 spots/µm^2^ for each color (total spot number densities of 1.0 ± 0.5 spots/µm^2^; for conciseness, we will describe as ≈1 spot/µm^2^ throughout this report), and then performed simultaneous dual-color single-molecule observations at normal video rate (30 Hz) at 37°C, using a home-built total internal reflection fluorescence (TIRF) microscope.^44, 45^

Virtually all of the SNAPf-OR fluorescent spots exhibited diffusion in the PM, but in addition, they often exhibited temporary colocalization and co-diffusion,^37^ suggesting the frequent formation of transient homodimers (Supplementary videos 1 and 2; Fig. 1a). We focused on simultaneous two-color experiments, due to the ease of image analysis. For the single-color imaging data, see Supplementary Fig. 2a, b, which show an example of repeated homodimerizations of a KOR molecule with different partner KOR molecules, consistent with the observations made with other GPCRs.^35, 36, 37^

**Fig 1:**
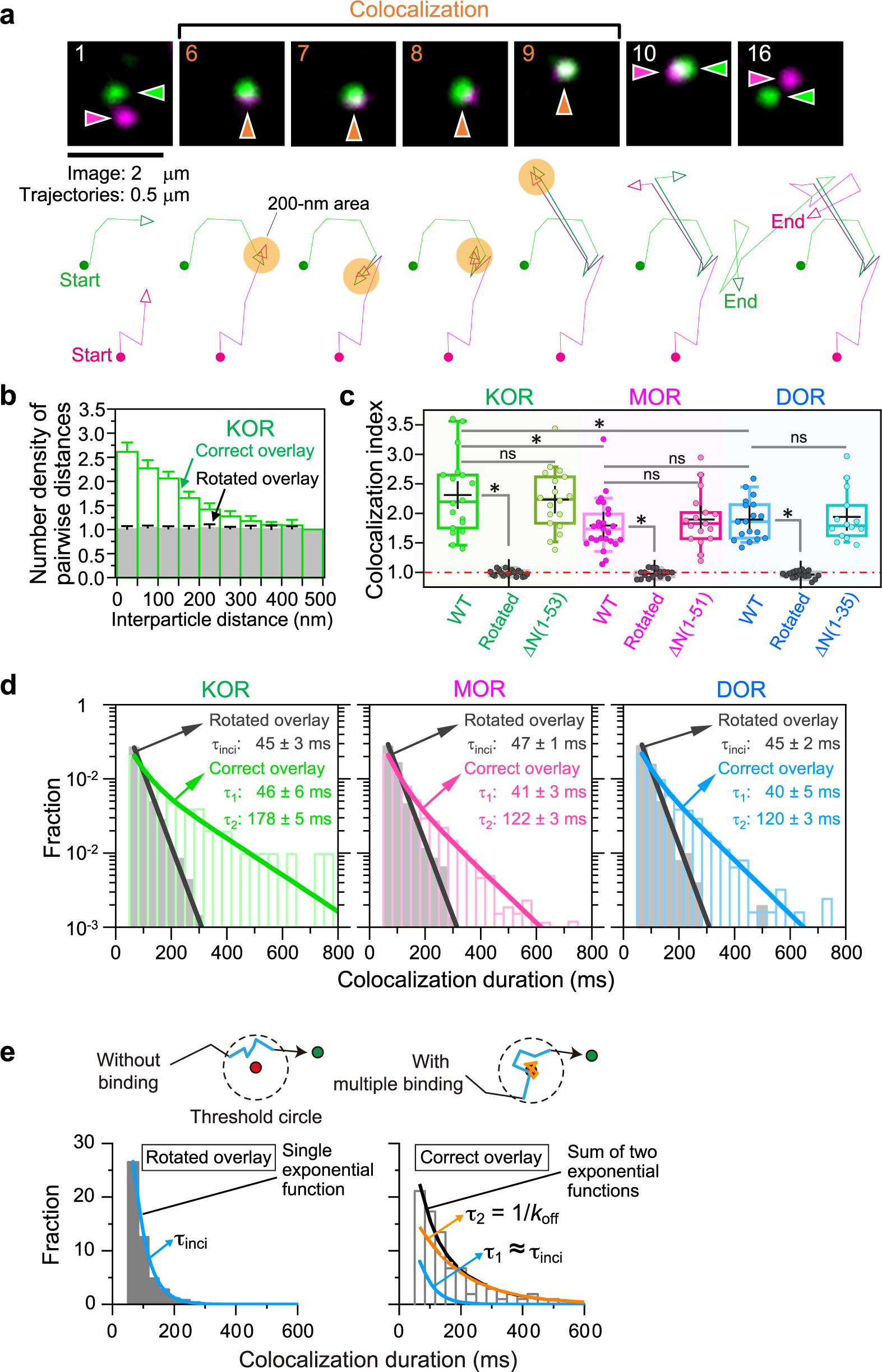
ORs in the PM form metastable homodimers with lifetimes of 120∼150 ms, without involving the N-terminal extracellular domain. **a** Typical image sequence of simultaneous two-color single fluorescent molecule observations, showing transient homo-colocalization and co-diffusion of single molecules of SNAPf-KOR tagged with SNAP-CF 660R (magenta) and SNAP-Surface 549 (green), with their trajectories. **b** PCCFs for the SNAPf-KOR spots (mean ± SEM after area normalization; 18 movies). Green and grey bars indicate the results of the correct and 180°-rotated overlays of the simultaneously observed magenta and green images. The colocalization index is defined as the mean of PCCF values for 0-100 nm divided by the mean of PCCF values for 400-500 nm (see Supplementary Fig. 2c-e). **c** Colocalization indexes showing the homo-colocalizations of three ORs and OR mutants with deletions of the N-terminal extracellular domains. *Throughout this study, we employ the following conventions*: for the box plots, horizontal bars, crosses, boxes, and whiskers indicate median values, mean values, interquartile ranges (25–75%), and 10-90% ranges, respectively; for the bar graphs and tables, mean ± SEM values are provided; and for the statistical analyses, * and ns represent significant (*p* < 0.05) and non-significant (*p* ≥ 0.05) differences, respectively. The *p* values were obtained by Tukey’s multiple comparison test, except the Brunner-Munzel test used for colocalization lifetimes. All of the quantitative results are summarized in Table 1 and their statistical parameters (sample size *n* and *p* values) are provided in Supplementary Table 1. **d** Histograms showing the distributions of colocalization *durations* for correct and rotated overlays. The control histograms for rotated overlays (grey) were fitted by single exponential functions (black), providing the lifetime of the incidental overlap events (*τ*_inci_). The histograms for correct overlays (colors) were fitted by the sum of two exponential functions: The faster decay time (*τ*_1_) was close to *τ*_inci_, and the slower decay time provided the true homodimer lifetime (*τ*_2_; see Supplementary Note 1 for the theory). The homodimer lifetimes, after correction for the trackable duration lifetimes of the two fluorescent probes, are shown in the boxes (summarized in Table 1). **e** The method to determine the true dissociation rate constant (*k*_off_), considering the multiple binding events during an observed colocalization duration. See Supplementary Note 1.

**Table 1.**
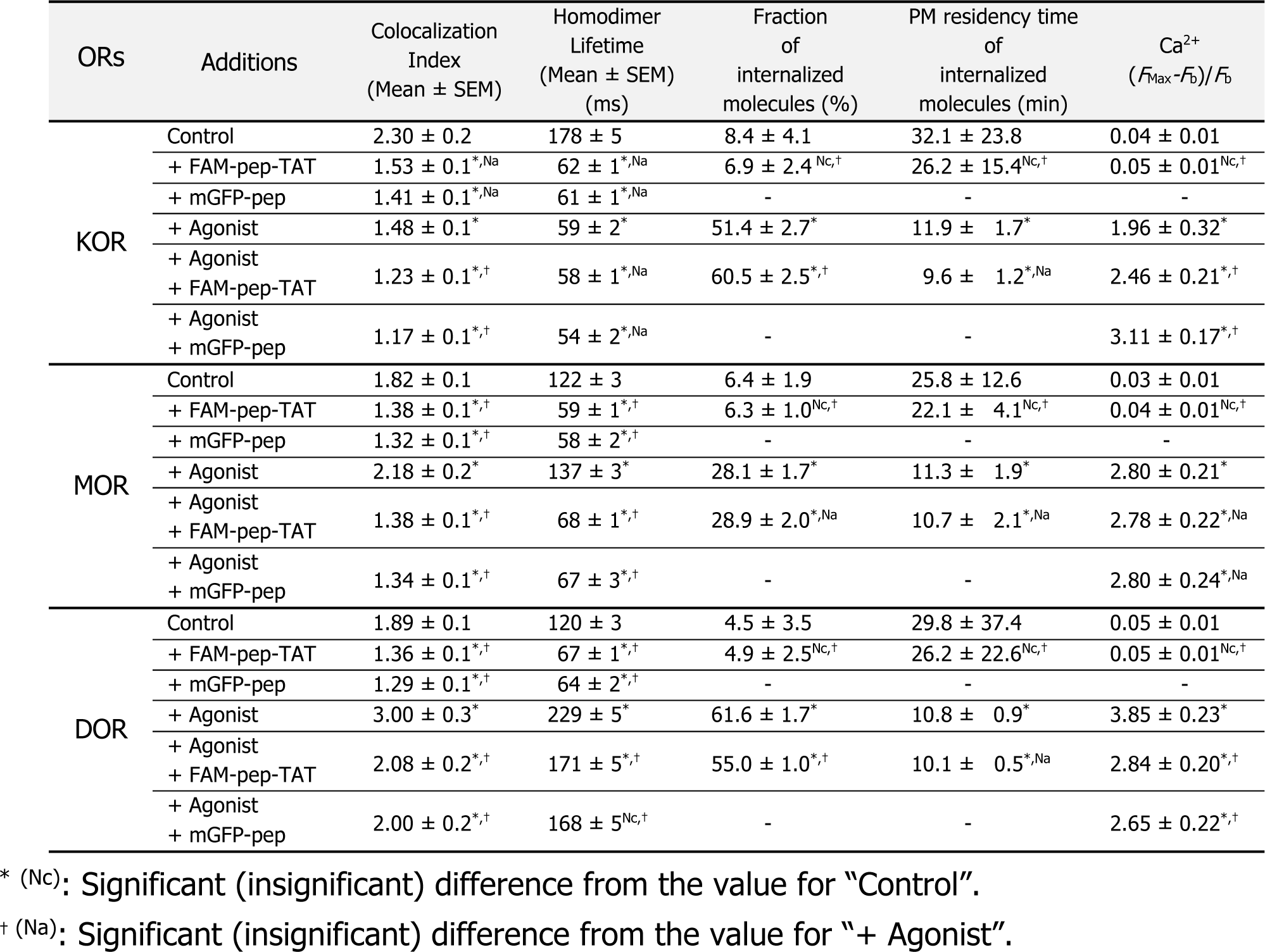
Summary of the colocalization indexes, homodimer lifetimes, fractions and PM residency times of molecules with internalizations detectable by the 35-min observations, and Ca^2+^ mobilization parameter, (*F*_Max_-*F*_b_)/*F*_b_, in the presence and absence of various modulators (0.2 µM agonists, ≈3 µM FAM-Xpep-TATs in the cytoplasm, and ≈6 µM mGFP-Xpeps in the cytoplasm). The agonists employed here were U-50488 for KOR, DAMGO for MOR, and SNC-80 for DOR.

To quantitatively examine the extent of the colocalization of two single molecules, we employed a “colocalization index”, which parameterizes the propensity of a pair of magenta and green molecules to become localized within 100 nm. Namely, the colocalization index is defined from the pair cross-correlation function (PCCF) histogram for the SNAP-Surface 549 (green) and SNAP-CF660R (magenta) spots (Fig. 1b),^46^ as the ratio of PCCF (0-100 nm) vs. PCCF (400-500 nm) (Supplementary Fig. 2c-e). When a molecular interaction does not exist, the colocalization index is equal to one within experimental uncertainty. When it does exist, the index becomes significantly greater than 1. As a negative control (including incidental colocalization), the 180°-rotated magenta image was superimposed on the green image (see Supplementary Fig. 2d, grey histogram in Fig. 1b), providing a colocalization index of ≈1. KOR, MOR, and DOR exhibited significantly higher colocalization indexes than the negative controls (Fig. 1c), suggesting that they form homodimers (and possibly greater oligomers when the expression levels are enhanced).

### Homodimer dissociation rate constant k_off_ (inverse of the dimer lifetime) can be obtained from single-molecule colocalization experiments

To evaluate the homodimer lifetimes, each time we found the colocalization event of spots with different colors within a threshold distance R = 200 nm (see Supplementary Note 1, which shows that the selection of R does not affect the measured k_off_ value, although it could affect its SEM),^36, 37, 44, 45^ its duration was measured. By measuring the durations of >1,500 events (in ≥17 cells; all statistical parameters are summarized in Supplementary Table 1), we obtained the distribution of colocalization durations (Fig. 1d).

To evaluate the homodimer dissociation rate constant k_off_ (inverse of the homodimer lifetime) from the distribution of colocalized durations, we developed a theory based on the diffusion equation that predicts the distribution of colocalization durations (Supplementary Note 1). The crucial result of this theory is that, from the histogram of the colocalization durations, we can rigorously estimate the homodimer dissociation rate constant k_off_ and thus the homodimer lifetime *τ*_2_. This result holds despite the fact that the generally used colocalization threshold distance is ≈200 nm, which is much greater than the molecular scale of 3∼10 nm. The theory shows that, at a time resolution of 33 ms and under the conditions of quite limited signal-to-noise ratios in our experiments, the histogram of the colocalization durations can be fitted with the sum of two exponential functions, where the two decay constants represent the incidental colocalization lifetime (1/α) and the inverse of the dissociation rate constant (1/k_off_).

Previously, we^36, 37, 45^ and others^33, 34, 35^ depended on intuitive methods to obtain the homodimer lifetimes from optical colocalization data. However, due to the theory developed here, we now have a firm basis to obtain the homodimer dissociation rate constant k_off_ from the distribution of single-molecule colocalization durations.

### Brief outline of the theory to evaluate k_off_ from the single-molecule colocalization duration distribution (Supplementary Note 1)

In dual-color single-molecule tracking, a colocalization is defined as an event in which two spots are located closer than a threshold distance R. Starting from the 2D diffusion equation, we found that the distribution of the dwell times in the threshold circle without binding (incidental lifetime *τ*_inci_; i.e., incidental colocalization duration, Fig. 1e) is expressed by

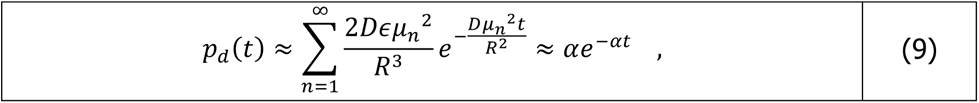

where D is the diffusion coefficient, R is the threshold distance, *ε* is a regularization factor, and μ_n_ is the n-th real positive 0 of the Bessel function (Equation numbers in the main text are matched with those in Supplementary Note 1 and 2 for easy reference). Under our experimental conditions, the first term is sufficient to describe the distribution, providing a single exponential function with a decay time constant of R^2^/Dμ_1_^2^ = 1/α, which is the incidental colocalization lifetime (*τ*_inci_). We indeed found that the colocalization time distribution for incidental colocalizations obtained for the overlay on the 180°-rotated image (Supplementary Fig. 2d) could be fitted with a single exponential function (using both Akaike’s and Bayesian information criteria; AIC and BIC, respectively) (Fig. 1d).

The dwell time in the threshold circle with binding (T; i.e., colocalization duration, Fig. 1e) is given by the sum of *τ*_inci_ added to the sum of multiple binding durations, *τ*_i_.

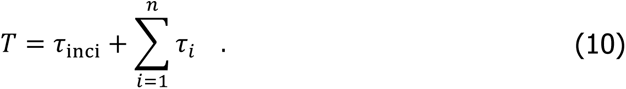

Here, the terms in the sum correspond to possible multiple binding events during a dwell; i.e., to obtain the correct k_off_ value, we will need to consider the cases where the observed colocalization durations are the results of multiple binding events.

Combining these results, we obtain the dwell time distribution in the threshold circle.

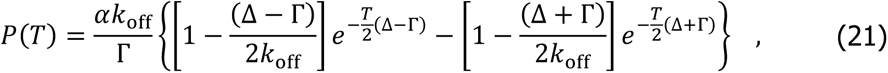

where Δ = (*α* + *k*_off_ + *k*_on_) and 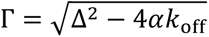.

At a time resolution of 33 ms and under the conditions of the quite limited signal-to-noise ratios in our experiments, we obtain

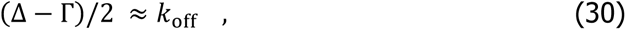

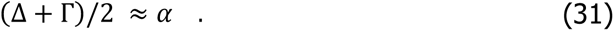

This result explains why the histogram of the colocalization durations can be fitted by the sum of two exponential functions. Most importantly, it permits to identify the two decay constants with the incidental colocalization lifetime (1/α = *τ*_inci_) and the inverse of the dissociation rate constant (1/k_off_), as anticipated.

### OR homodimer dissociation rate constant k_off_ and true homodimer lifetime

As shown in Fig. 1d, we were able to fit the three histograms for KOR, MOR, and DOR, obtained from the correctly overlaid images, with the sum of two exponential functions, as predicted by equations 30 and 31. The fitting provided two time constants, *τ*_1_ and *τ*_2_ (*τ*_1_ ≤ *τ*_2_). The histogram obtained from the rotated overlay images could be fitted with a single exponential function, which provided the time constant *τ*_inci_ (= 1/α) (the numbers of significant exponential terms were always confirmed by both AIC and BIC in this work). We found that *τ*_1_ ≈ *τ*_inci_ for all three ORs. The longer decay time constant obtained from the histogram for the correctly overlaid images (*τ*_2_) provides k_off_ (= 1/*τ*_2_; Supplementary Note 1). As such, using this theory, by simply plotting the histogram of the colocalization durations, we can obtain the correct homodimer dissociation rate constant k_off_ (by taking account of the multiple binding-dissociation cycles).

The homodimer lifetimes of KOR, MOR, and DOR were estimated to be 178 ± 5, 122 ± 3, and 120 ± 3 ms, respectively, from the histograms shown in Fig. 1d (after correction for the trackable duration lifetimes of the two fluorescent probes). These lifetimes are considerably shorter than that reported for KOR (≈1.8 s at room temperature, perhaps ≈20°C).^33^ This difference might be predominantly explained by the lower temperatures employed in the previous study (20°C), whereas our observations were performed at 37°C. In addition, it is possible that our imaging and analysis methods are more sensitive to short-term colocalization events, compared with previous investigations.

### Determination of OR homodimer-monomer equilibrium constant (K_D_) from PCCF

In the literature, the question of whether ORs (or GPCRs or any other membrane proteins) form homodimers or not has been extensively debated. However, the facts of the matter are that many membrane proteins would form homodimers and clusters anyway at higher expression levels, and that the physiological expression levels vary greatly from cell to cell.

Therefore, the critical point should be to quantitatively estimate the homodimer affinity; i.e., to evaluate the homodimer dissociation (homodimer-monomer) equilibrium constant K_D_. Here, we developed a theory (Supplementary Note 2) for evaluating K_D_ from the PCCF of the OR fluorescently-labeled in two colors (PCCF in the form of a histogram, as shown in Fig. 1b and Supplementary Fig. 2e), under conditions where the expression levels are low and so the presence of oligomers greater than dimers could be neglected. The K_D_ values evaluated from the PCCF by this method are quite accurate because we employed the expression conditions of ≈1 fluorescent spot/µm^2^ (Supplementary Fig. 3c, e): Within this range, the colocalization index barely depended on the expression levels when K_D_ >6.5 copies/µm^2^ (see the simulation results shown in Supplementary Fig. 3a-e, particularly Supplementary Fig. 3c). At these copy number densities, we hardly detected oligomers greater than dimers.

The values of K_D_ were 5.87 ± 0.56, 15.24 ± 1.54, and 16.42 ± 0.53 copies/µm^2^ for KOR, MOR, and DOR at 37°C, respectively (Fig. 2a and b; K_D_ was first determined for each movie using the PCCF and the number of fluorescent spots obtained from the movie, fixed fluorescent labeling efficiencies of 0.60 for SNAP-SNAP-Surface549 and 0.61 for SNAP-CF660R, and fitting parameters of K_D_ and the precision of overlaying two-color movies plus single-molecule localizations for the two fluorescent probes, and then the arithmetic mean (± SEM) K_D_ was obtained using K_D_s determined for all the movies; the number of movies are provided in Supplementary Table 1, like other statistical parameters). These K_D_ values mean that KOR is more likely to exist as homodimers than MOR and DOR (the main results are summarized in Table 1 and their statistical test results are in Supplementary Table 1). Our estimated values of K_D_ for the MOR and KOR homodimers found here exhibited much smaller errors compared with previous estimates (for MOR,^33^ 27.43 ± 11.75 copies/µm^2^; and for KOR,^30^ 32 ± 15 copies/µm^2^), and their mean values are smaller by factors of approximately 1.7 and 4.9, respectively. In contrast, these K_D_ values are greater than the K_D_’s of β2AR and FPR homodimers (1.6 and 3.6 copies/μm^2^, respectively).^19, 37^ The K_D_ values for CD28, the constitutive disulfide-linked dimer control molecule,^47^ and the transmembrane domain of the LDL receptor (TM^LDLR^), the monomer control molecule,^37^ are 0.041 ± 0.072 and 132 ±21 copies/µm^2^, respectively, providing the basic dynamic range (4400x) for this method to evaluate K_D_ values (although the value for CD28 is a nominal value, representing the experimental limitation). Our results indicate that the KOR homodimer affinities are higher than those of DOR and MOR, and thus partially explain previous data showing that KOR, but not DOR or MOR, forms homodimers at low membrane densities.^30^ In the following part of this report and in the companion paper, we use the colocalization index when showing the direct experimental data is preferrable, whereas we show the K_D_ values when providing the fundamental constants is desirable. Under our standard experimental conditions, the colocalization index is simply related to K_D_ as shown in Fig. 2c, and K_D_ can be roughly evaluated using the graphs shown there, although we performed actual fitting for the PCCF for each cell to obtain the K_D_ value for each cell (the final K_D_ value and its SEM were obtained as the arithmetic mean of the K_D_s for all the observed cells).

**Fig 2:**
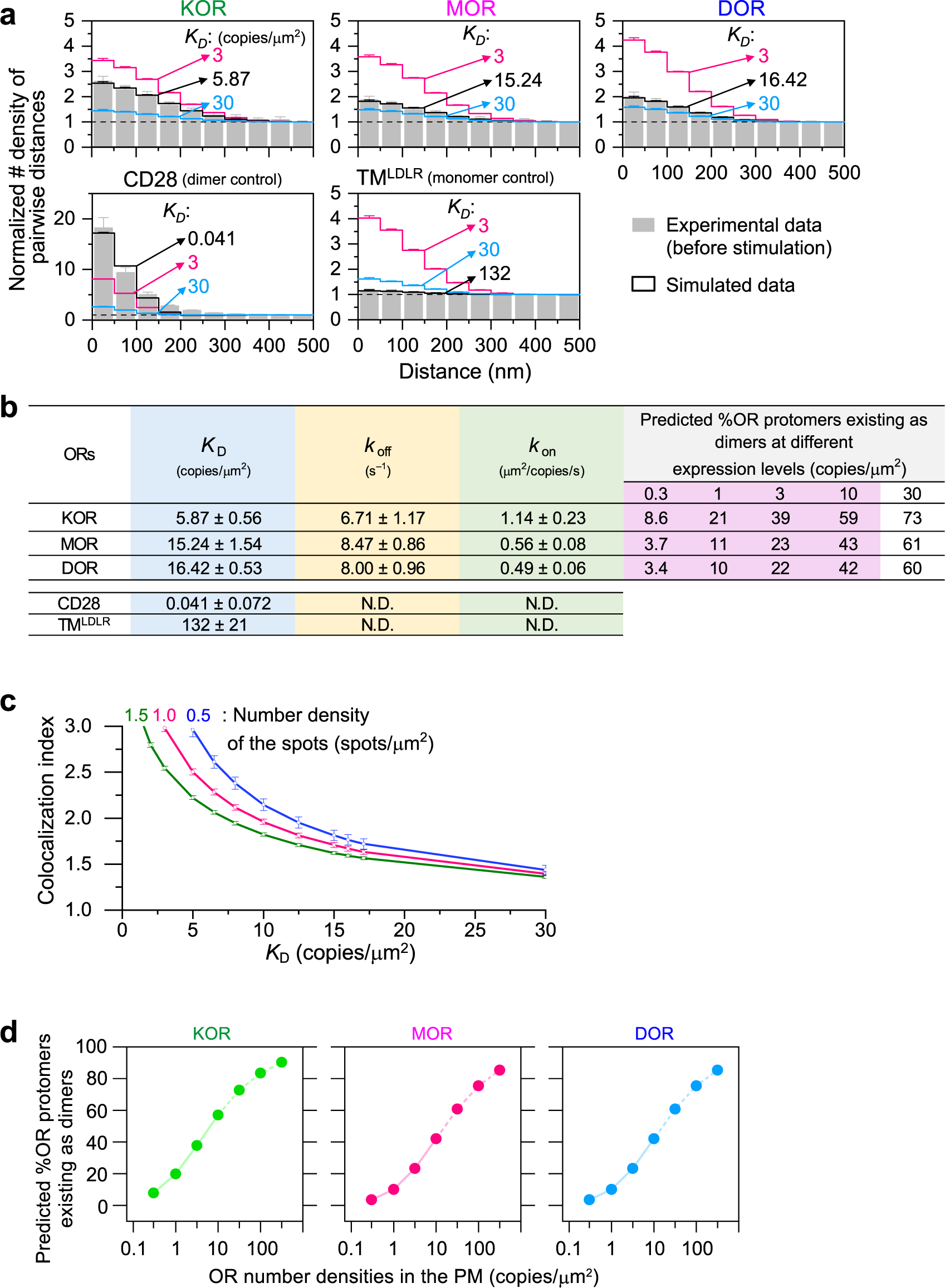
Dynamic OR homodimer-monomer equilibrium constants (*K*_D_s) and homodimer formation and dissociation rate constants (*K*_on_ and *K*_off_, respectively), providing the %protomers existing as dimers as a function of the expression level. **a** Experimental PCCFs for KOR, MOR, DOR, CD28, and TM^LDLR^ (the PCCF for KOR is the same as that shown in Fig. 1c) fitted with theoretical PCCFs (Eq. 24) in Supplementary Note 2; the total numbers of fluorescent spots, *sN_TA_* and *sN_TB_*, were determined from experiments; labeling efficiency = 0.7 [Supplementary Fig. 1e]) using *K*_D_ and σ as fitting parameters. CD28 and TM^LDLR^ were dimer and monomer references, respectively. The black lines indicate the best-fit functions, and their *K*_D_ values are shown. The PCCFs with *K*_D_ values of 3 and 30 are also shown for comparison. **b** Summary of *K*_D_, *K*_off_ and *K*_on_ values for the three ORs without agonist, and also for CD28 and TM^LDLR^. N.D.: not done. Predicted percentages of OR protomers existing as dimers at various expression levels calculated from the *K*_D_s are also shown. Physiological expression levels found in the literature (1-10 copies/µm^2^) are highlighted by a pink background (see the main text). **c** Colocalization index is simply related to *K*_D_, if the labeling efficiency, the precision of single-molecule localization plus image overlaying, and the number density of fluorescent molecules in each movie were the same (0.7, 140 nm, and 0.5, 1, or 1.5 fluorescent spots/µm^2^, respectively). *K*_D_ can be roughly estimated from the colocalization index using these curves. **d** Predicted percentages of OR protomers existing as dimers, plotted as a function of expression levels (without agonist). The curves indicate the mathematical functions calculated from the *K*_D_ values, and solid curves are used for physiological expression levels found in the literature (0.32-10 copies/µm^2^). Tick marks between the exact digits represent ≈3.16x (10^0.5^x) of the smaller digit.

Using *K*_off_ and *K*_D_, *K*_on_ can be calculated as *K*_off_/*K*_D_ (equation 1). Direct determination of *K*_on_ from single-molecule movies is only possible when the labeling efficiency is 100%. This is because the fluorescent spots that appear to be monomers (protomers) of A and B molecules might be dimers of labeled and non-labeled molecules. For the correction to evaluate the actual numbers of protomers, one would need to know *K*_D_, and thus *K*_on_ cannot be determined independently from *K*_D_. The percentages (in copy numbers) of molecules (protomers) that exist as dimers at various expression levels (number density of molecules) were calculated from the obtained *K*_D_ values (Fig. 2d). Their overall variations are in the range of 3.4-59% of protomers in homodimers over the ORs’ predicted physiological expression ranges of 0.3 - 10 copies/µm^2^.^26, 29, 33^

Therefore, substantial amounts of homodimers would exist in various nerve tissues at any time. The expected average protomer copy numbers existing as monomers and homodimers per cell at various expression levels are shown in Fig. 2b and 2d. However, all of these homodimers are forming and dispersing all the time, with lifetimes shorter than 0.2 s (Fig. 1d). Once dissociated into monomers, they will again form homodimers, often with different partner molecules (Supplementary Fig. 2a, b), which will occur more readily at higher expression levels. Therefore, although at any moment, 10-21% of ORs might exist as homodimers at the expression levels we employed (≈1 copy/µm^2^), the actual molecules existing as dimers are turning over all the time, and within a few seconds (even at an expression level of 1 copy/µm^2^; calculated as 1/*K*_off_ + 1/[*K*_on_ x 1]), virtually all of the molecules experience the periods of homodimers and monomers.

A previous biochemical study detected DOR homodimers^9^ and a single-molecule imaging study using a fluorescently-conjugated MOR antagonist detected MOR homodimers,^33^ in agreement with our results. Meanwhile, at apparent variance with our results, previous single-molecule examinations demonstrated that MOR and KOR tend to exist as monomers even at 20°C, at expression levels between 0.1 and 0.3 copies/µm^2^.^14, 31^ Since we employed somewhat higher number densities of ≈1 OR spot/µm^2^, the discrepancy might simply be due to the number densities of homodimers found in an image. This underscores the critical importance of evaluating the homodimer-monomer dissociation equilibrium constant *K*_D_, rather than simply detecting homodimers.^37^

### The 9-26 amino-acid sequences in the C-terminal cytoplasmic domains play critical roles in OR homodimerization

To advance our understanding of OR homodimerization mechanism, we determined the OR domains and amino acid sequences responsible for homodimerization. First, we examined the involvement of the extracellular N-terminal domains in homodimerization, using the N-terminal deletion mutants (KOR(Δ1-53), MOR(Δ1-51), and DOR(Δ1-35)). Given that the amino-acid homologies in the N-terminal domains among the three ORs are quite low (Supplementary Fig. 4a), we considered a possibility that they might be used to distinguish particular homo- and/or hetero-dimers. These amino-acid sequence ranges were selected partly because the removal of the entire N-terminal extracellular domains blocked the OR expression in the PM, probably due to the deletion of the signal sequences. In addition, these deletion mutants were previously used in X-ray crystallography studies to determine the structures of the remaining parts of ORs.^48, 49, 50, 51^

The N-terminal deletion mutants of the three ORs all exhibited diffusion behaviors similar to those of the wild types (without immobilization or clustering) and virtually the same colocalization indexes as those of the wild types (Fig. 1c). These results clearly indicate that the N-terminal extracellular domains are not responsible for homodimerization (see the companion paper about the N-terminal domain involvement in heterodimer formation).

We next examined the involvement of the C-terminal cytoplasmic domains. Their amino-acid homologies are also low, and in addition, a previous biochemical analysis indicated that the 15-amino acid stretch in the C-terminal domain is responsible for DOR homodimerization.^9^ We examined systematically varied C-terminal deletion mutants (Fig. 3a, b). They hardly exhibited immobility or clustering, and their colocalization indexes showed that KOR’s aa 365-380, MOR’s aa 358-382, and DOR’s aa 357-372 are critical for homodimerization (Fig. 3b and Supplementary Table 1). For DOR, this amino-acid sequence agrees with the previous biochemical data.^9^ Meanwhile, the mean colocalization index values never decreased to the index values comparable to those for rotated overlays (≈1) (Fig. 3b) or a monomer control molecule TM^LDLR^ (≈1.1) (Fig. 2a), suggesting the possibility that there might be other weaker homodimerization site(s), which might be the TM domains, as previously suggested for ORs^26, 49, 50, 52, 53^ and other class A GPCRs.^21, 39, 40, 41^ However, as shown in Supplementary Fig. 4a; the aa identities of TM domains among the three ORs are quite high, and thus the homo-interaction between TM domains might be less specific. We will revisit the TM interactions for homodimerization later.

**Fig. 3:**
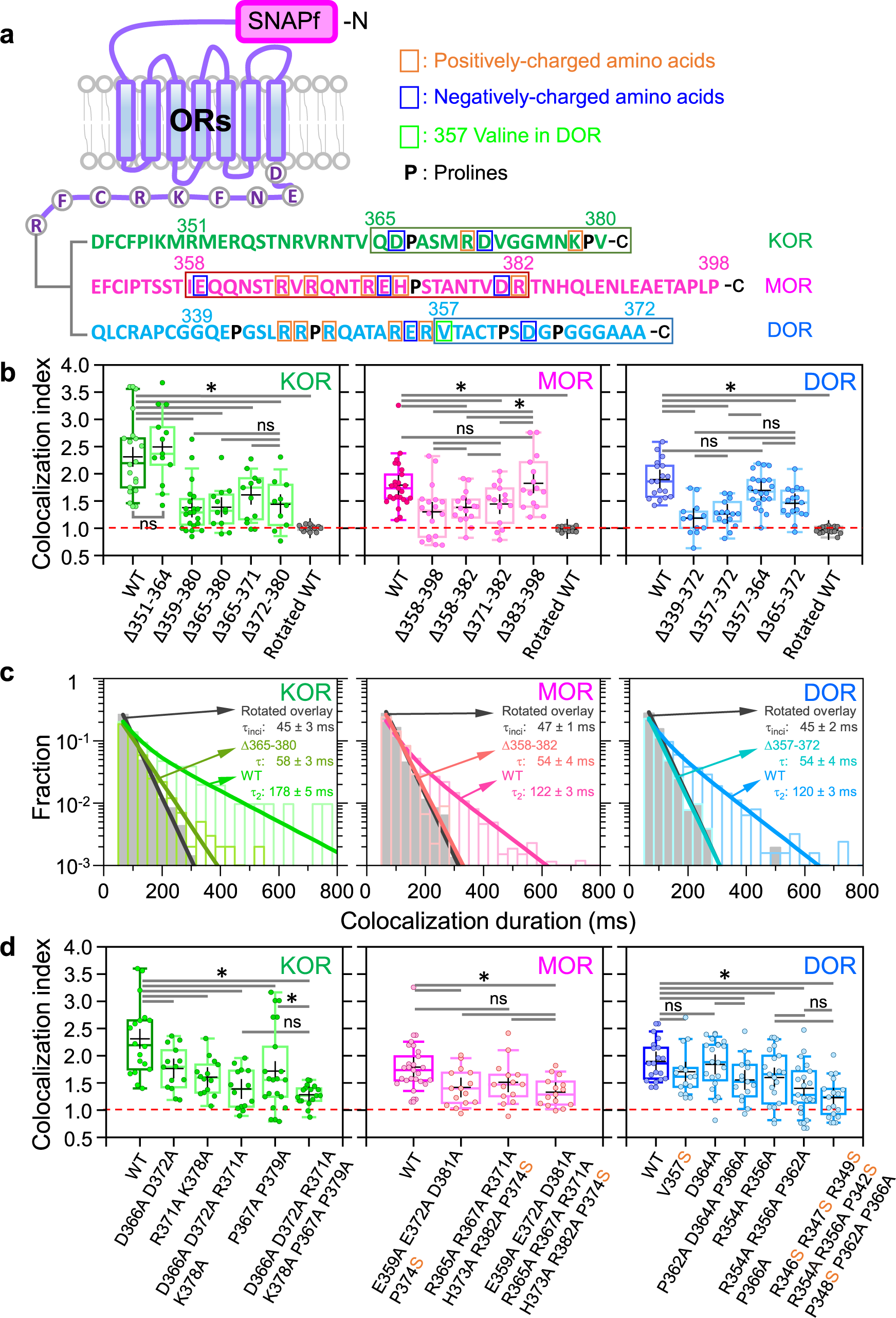
C-terminal cytoplasmic domains are predominantly responsible for OR homodimerization. **a** Schematic figure showing the amino-acid sequences of the C-terminal cytoplasmic domains of the three ORs. The sequences surrounded by rectangles were extensively examined for their involvement in OR homodimerization. **b** Colocalization indexes for WT and various C-terminal cytoplasmic deletion mutants, showing that KOR aa 365-380, MOR aa 358-382, and DOR aa 357-372, with very low amino-acid homologies, are critical for homodimerization. **c** Histograms showing the distributions of colocalization *durations* for the C-terminal cytoplasmic deletion mutants, KOR(Δ365-380), MOR(Δ358-382), and DOR(Δ357-372) (only the results of correct overlays). WT-OR data are reproduced for comparison (for both correct and rotated overlays; Fig. 1d). Deletion mutants exhibited distributions that could be fitted by single exponential functions, and their decay time constants *τ*s (after correction for the trackable duration lifetimes of the two fluorescent probes) are indicated in the boxes. **d** The colocalization indexes of various C-terminal point mutants, indicating that the charged groups and prolines in the C-terminal cytoplasmic domains play important roles in OR homodimerization.

The distribution (histogram) of the colocalization durations for each representative deletion mutant, KOR(Δ365-380), MOR(Δ358-382), or DOR(Δ357-372), could be fitted by a single exponential function, rather than the sum of two exponential functions (based on both AIC and BIC). The decay time constants were only slightly longer than the incidental colocalization lifetimes (Fig. 3c); i.e., the *τ*_2_ component, representing the homodimer lifetime fraction, is small and /or the *τ*_2_ is short, and thus the *τ*_2_ component could not be resolved from the incidental colocalization lifetime decay component.

By introducing point mutations in these critical regions for homodimerization and their surrounding regions (Fig. 3a), we found that a few to several basic/acidic residues, as well as proline residues, are involved in the OR homodimerization (for particular amino acids, see the colocalization indexes shown in Fig. 3d), suggesting that the electrostatic interactions and the overall structure of the cytoplasmic C-terminal domain could be important for homodimerization. Indeed, the IUPred2 scores (energy-estimation-based predictions for ordered and disordered residues; http://iupred2a.elte.hu)^54, 55^ indicated that the C-terminal region could be weakly intrinsically disordered (Supplementary Fig. 4b), and the scores decreased (more ordered) with these point mutations (Supplementary Fig. 4c). Accordingly, in addition to the specific, relatively strong amino-acid interactions, multiple specific, but weak interactions, made possible by the flexible intrinsically disordered regions (IDRs) of the ORs’ C-terminal regions, might facilitate the OR homodimerization. Such a mechanism resembling liquid-liquid phase separation was previously found for the dimerization of the transcription factor PU.1.^56^ Furthermore, the IDRs in the C-terminal regions might be necessary to bring the specific binding sites closer so that the actual binding can occur.^57^

### Peptides mimicking the homodimerization sites block homodimer formation

We then examined whether peptides with the same amino-acid sequences as those of the deleted parts of the mutants could block homodimerization. The use of these peptides was critical for unequivocally demonstrating that the specific amino-acid sequences in the C-terminal regions of KOR, MOR, and DOR are responsible for homodimerizations, because the deletion and point mutations might have induced conformational changes of the true homodimer interaction sites, thus inhibiting homodimerization. Furthermore, if we could develop such homodimerization blockers, they could become extremely useful tools to dissect the functions of OR monomers and homodimers, and might inform future developments of more effective analgesic treatments with fewer side effects and less tolerance development.

We employed two approaches. The first one involved the expression of peptides with the same amino-acid sequences as those of the deleted parts of the mutants, conjugated to the C-terminus of mGFP (named mGFP-Kpep, Mpep, and Dpep, collectively called mGFP-Xpeps; numbers in parentheses following mGFP-Xpeps indicate the amino-acid residue ranges in the wild-type ORs) in CHO-K1 cells stably expressing SNAPf-KOR, MOR, or DOR, respectively (Fig. 4a, b). The concentration of the cytoplasmic mGFP-Xpep was measured by confocal fluorescence microscopy, based on calibration with various concentrations of purified mGFP protein dissolved in Ham’s F12 observation medium in the glass-base dish (Supplementary Fig. 5a; see Methods).

**Fig 4:**
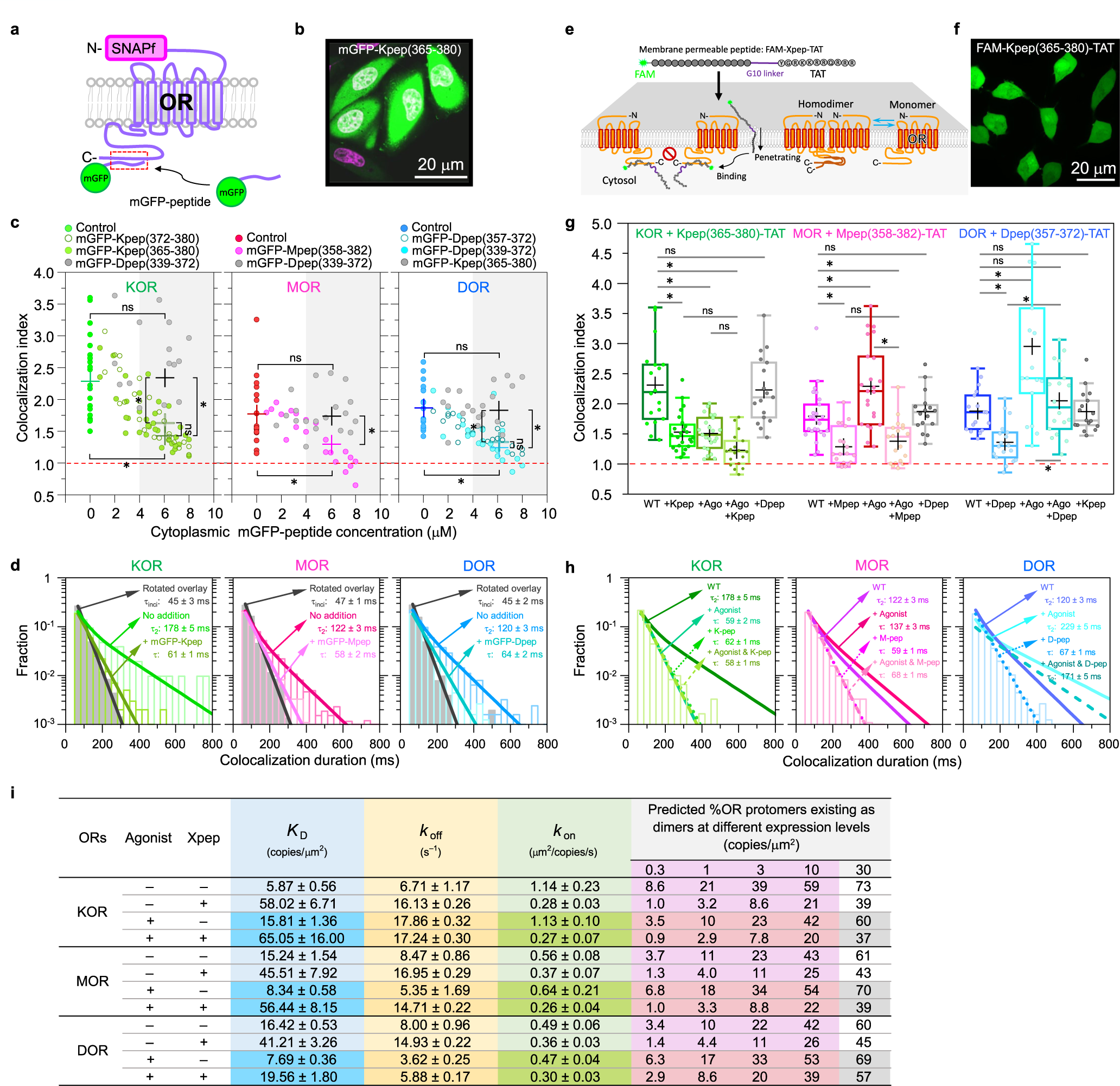
Specific C-terminal-domain peptides suppress OR homodimerization. **a** Schematic figure showing the experimental design to examine the effects of mGFP-Xpeps expressed in the cell on OR homodimerization. **b** Representative confocal image of CHO-K1 cells transfected with cDNAs encoding SNAPf-KOR (not visible here; only visible by single-molecule imaging conditions) and mGFP-Kpep (green), co-stained with the Live 650 Nuclear Stain (magenta; white nuclei indicate the entrance of mGFP-Kpep into the nuclei, whereas a magenta nucleus indicates the cell without mGFP-Kpep expression). **c** The OR homo-colocalization index in each cell tends to decrease with an increase of the mGFP-Xpep concentration in the cytoplasm. The + keys at x = 0 and 6 μM mGFP-Xpep indicate the mean value of the colocalization indexes without mGFP-Xpep expression and that averaged over all the data points in the range of 3.8-7.8 μM mGFP-Xpep, respectively. Significant reductions of the colocalization indexes in the presence of the mGFP-Xpeps, but not the control mGFP-peptides, were found for all three ORs (Table 1). **d** Histograms showing the *duration* distributions of transient homodimers of SNAPf-ORs in cells co-expressing 3.8-7.8 μM mGFP-Xpeps (only the results of correct overlays; the value for each bin is the mean of 18 cells), compared with those without co-expression (the same as those shown in Fig. 1d; for both correct and rotated overlays). For further details of these graphs, see the caption to Fig. 3c. **e** Schematic figure showing the experimental design to examine the effect of the FAM-Xpep-TATs incorporated in the cell on OR homodimerization. **f** Representative confocal image of CHO-K1 cells containing an average of 3.4 μM FAM-Kpep-TAT in the cytoplasm. **g, h** Agonist addition (0.2 μM) suppressed KOR homodimerization, but enhanced the homodimerization of MOR and DOR (colocalization indexes in **g** and durations of colocalization events in **h**). The presence of ≈3 μM FAM-Xpep-TAT in the cytoplasm suppressed homodimerization both before and 2-5 min after the agonist addition. (**i**) Summary of *K*_D_ (homodimer-monomer equilibrium constant), *K*_off_ (dimer dissociation rate constant), and *K*_on_ (dimer formation rate constant) for three ORs in the presence and absence of ≈3 μM FAM-Xpep-TAT in the cytoplasm, both before and 2-5 min after the addition of agonists. Predicted percentages of OR protomers existing as dimers at various expression levels calculated from the *K*_D_s are also shown. Physiological expression levels found in the literature are highlighted by a pink background (see the main text).

The OR homo-colocalization index in the cells expressing various concentrations of mGFP-Xpep in the cytoplasm (homogeneously distributed throughout the cytoplasm; Fig. 4b) exhibited a clear tendency to decrease with an increase of the specific mGFP-Xpep concentration in the cytoplasm from 0 to 7.8 µM for all three ORs (Fig. 4c), whereas the control mGFP-peptides (mGFP-Dpep for KOR and MOR and mGFP-Kpep for DOR) had no effect (Fig. 4c). Consistently, the distributions of the colocalization durations in cells expressing 3.8-7.8 µM mGFP-Xpeps could be fitted by a single exponential function, rather than the sum of two exponential functions (based on both AIC and BIC; Fig. 4d), and the decay time constants became similar to those for the C-terminal deletion mutants (compare Fig. 4d with Fig. 3c; Table 1).

As the second approach, the K-, M-, and D-peps were conjugated with the fluorescent dye 5-FAM at their N-termini for visualization, and with the TAT sequence (YGRKKRRQRRR) (using a G10 linker sequence) at their C-termini for membrane permeabilization (FAM-Xpep-TAT; see Methods), and added to cells preincubated with 150 µM pyrenebutyrate for 5 min (Fig. 4e, f).^58^ The FAM-peptide-TATs exhibited diffuse spatial distributions throughout the cytosol (Fig. 4f). The cells containing 2.9-3.4 µM FAM-Xpep-TAT (see Supplementary Fig. 5b for the concentration calibration; in the following, we describe this concentration range as ≈3 µM for conciseness) were selected and the effects of these cytoplasmic peptides on the OR homodimerization were examined. The results indicated that ≈3 µM FAM-peptide-TAT significantly blocked the homo-dimerization (Fig. 4g; Table 1), consistent with the data obtained using mGFP-Xpeps (Fig. 4c). The control FAM-Xpep-TATs (FAM-Dpep-TAT for KOR and MOR and FAM-Kpep-TAT for DOR) did not affect the colocalization indexes. The effects on colocalization durations were consistent with these results and with those obtained by expressing mGFP-Xpeps (Fig. 4h; compare with Fig. 4d; Table 1).

The Xpep concentrations required for blocking homodimer formation might appear quite high (≈3 µM), compared with the ≈30-nM-level dissociation constants of various agonists and antagonists for ORs. However, note that the latter is the value for simple binding of the ligand to the receptor, whereas the former addresses the extremely efficient dimerization propensities of membrane molecules in the two-dimensional (2D) PM. The efficiency of dimerization in 2D space was previously found to be higher by a factor of 10^6^ than that in 3D space.^59^

The effects of the presence of ≈3 µM FAM-Xpep-TATs in the cytoplasm on the homodimer *K*_D_, *K*_off_, and *K*_on_ values are summarized in Fig. 4i. In addition to *K*_D_ and *K*_on_, FAM-Xpep-TATs increased *K*_off_ (enhanced dimer dissociation), indicating that the OR homodimerization cannot be described by simple first-order kinetics (simple binding reaction at the C-terminal domains), perhaps due to the conformational changes of the C-terminal domains induced by the peptide binding and/or to the presence of weaker secondary binding sites such as TM domains (Fig. 5, which will be discussed later). Nevertheless, we think the present simplified approach is useful as the starting point for quantitative understanding of the dynamic equilibrium between OR homodimers and monomers.

**Fig. 5:**
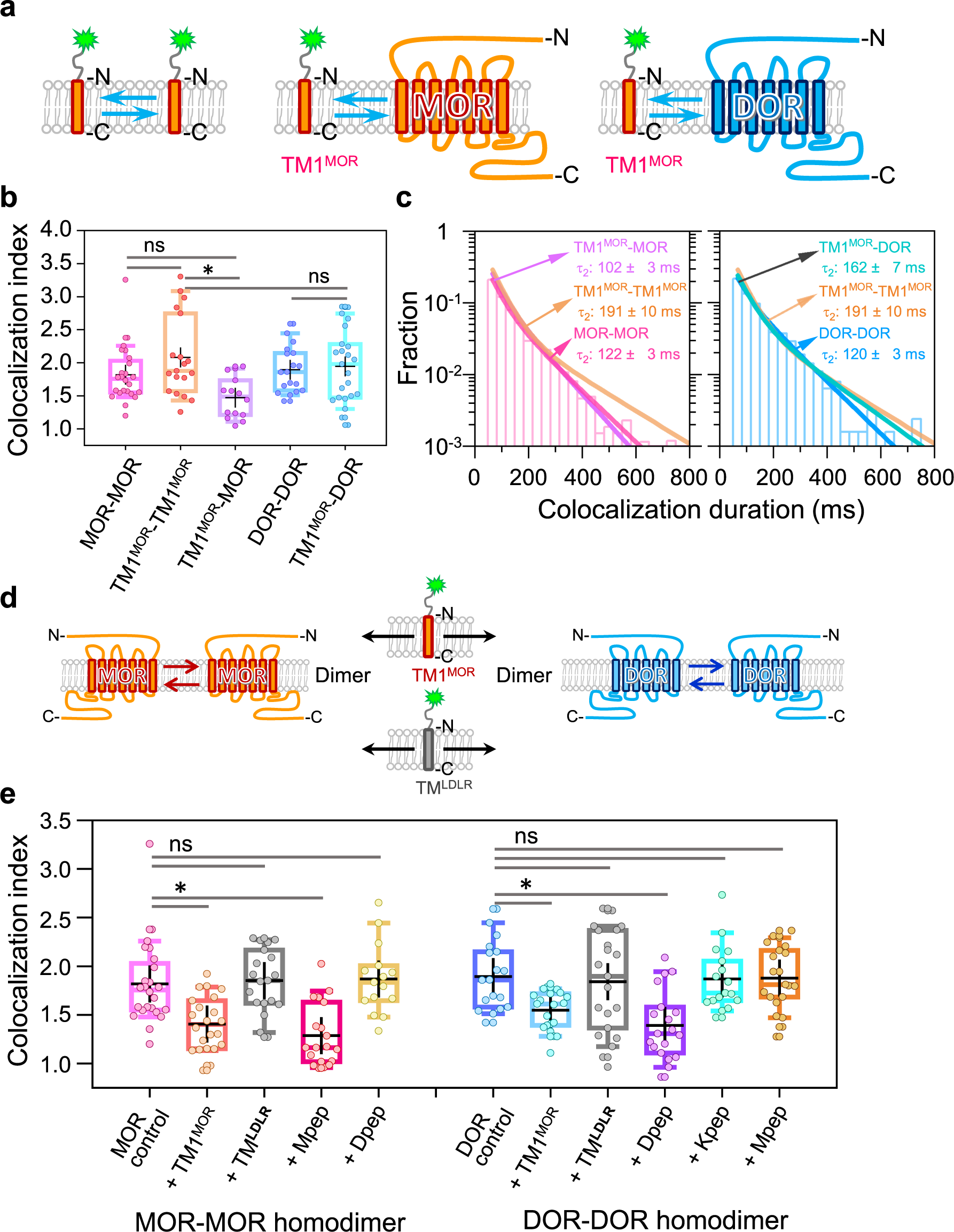
TM1^MOR^ suppresses both MOR and DOR homodimerization. **a** Schematic figure showing the experimental design for examining the interaction (binding) of TM1^MOR^ with itself, MOR, and DOR. **b, c** Colocalization indexes (**b**) and distributions of colocalization durations (**c**) showing that TM1^MOR^ forms metastable dimers with both MOR and DOR, suggesting that TM1^MOR^ might be involved in MOR-MOR homodimerization as well as MOR-DOR heterodimerization (see Figure 5 in the companion paper.^38^ **d** Schematic figure showing the experimental design for examining whether TM1^MOR^ can suppress homodimerizations of MOR and DOR. **e** TM1^MOR^ suppresses both MOR and DOR homodimerization, whereas Mpep and Dpep specifically suppress MOR or DOR homodimerization, respectively.

### Agonists modulate monomer-dimer interconversions, and FAM-Xpep-TAT peptides reduce their effects

The addition of representative agonists, U-50488 for KOR, [D-Ala^2^, N-Me-Phe^4^, Gly^5^-ol]-enkephalin acetate salt (DAMGO) for MOR, and SNC-80 for DOR, at 0.2 µM (a concentration sufficient to ligate virtually all OR molecules) affected the homodimerizations of the three ORs differently (observed during 2-5 min after agonist addition rather than after longer incubation because these agonists induce OR internalizations, which become evident after 5 min; discussed later in this report). The agonist-bound KOR exhibited less homodimerization (lower colocalization index), whereas the agonist-bound MOR and DOR exhibited more homodimerization (Fig. 4g and Table 1), although these results might vary depending on the particular agonist.^9, 14^ For example, the DOR agonists, DADLE, DSLET, and DPDPE, were previously found to induce fewer homodimers,^9^ but in the present research, another DOR agonist SNC-80 was found to increase the homodimer fraction, suggesting that the dimerization propensity depends on each agonist. We only examined one agonist for each OR subtype, because an extensive examination of the agonist effects on OR homodimerization is beyond the scope of the present work.

The colocalization indexes of the agonist-bound ORs were reduced by the addition of FAM-Xpep-TAT for all ORs, but in different ways depending on the OR type (Fig. 4g). The colocalization index of the agonist-bound KOR, which exhibited a smaller index as compared with that of non-ligated KOR, was further reduced by the presence of FAM-Kpep-TAT (although with no statistical significance). The colocalization index of the agonist-bound MOR was reduced in the presence of FAM-Mpep-TAT to a level near that for MOR with the FAM-Mpep-TAT without the agonist. The colocalization index of the agonist-bound DOR was reduced in the presence of FAM-Dpep-TAT, but was still greater than that of wild-type DOR in the absence of the agonist. The reason for this higher tendency to form homodimers even in the presence of FAM-Dpep-TAT is unknown, but we propose that the agonist-induced conformational changes enhanced the interaction in the TM domains, because DOR homodimerization also involves TM-domain interactions (see Fig. 5, which will be discussed in the next subsection). The effects of the agonists and homodimer blocker peptides on the colocalization durations were consistent with the colocalization index data (Fig. 4h), and the agonists’ effects on the homodimer *K*_D_, *K*_off_, and *K*_on_, as well as the percentages of OR protomers existing as homodimers are summarized in Fig. 4i. The actual copy numbers of OR protomers existing as monomers and homodimers at various total numbers of OR molecules in the entire PM in a cell are summarized in Supplementary Table 2.

### MOR’s transmembrane domain 1 (TM1^MOR^) is involved in MOR homodimerization with less specificity

Specific C-terminal cytoplasmic domain deletions/mutations and the addition of peptides with the same amino acid sequences as the deleted sequences greatly reduced the OR homodimers’ colocalization indexes, but never to be compatible with 1 found for rotated overlays or 1.1 found for the monomer control molecule TM^LDLR^ (Fig. 2a). Therefore, we considered the existence of other OR sites for homodimer formation. Since the involvement of transmembrane domains in OR homo- and hetero-dimerization has been proposed,^20, 21, 26, 49, 50, 52, 53^ and the first OR transmembrane domain (TM1) is often mentioned in the literature,^26, 49, 50^ we examined the involvement of MOR’s TM1 domain (TM1^MOR^) in MOR homodimerization (Fig. 5a).

Colocalization indexes indicated that TM1^MOR^ interacts with another TM1^MOR^ and MOR, as well as, importantly, with DOR (Fig. 5b; the fluorescent spot number densities of the two molecules in the PM were always adjusted to ≈1 spot/µm^2^, and thus the number ratio was approximately 1:1). These results suggest that TM1^MOR^ is involved in MOR homodimerization by the TM1^MOR^-TM1^MOR^ interaction, as well as, surprisingly, in MOR-DOR heterodimerization. These results are supported by the observation of the colocalization durations (Fig. 5c).

In contrast, in the homodimer blocking experiments where TM1^MOR^ is expressed ≈10x more than MOR or DOR (Fig. 5d), TM1^MOR^ moderately suppressed both MOR and DOR homodimers (positive controls, Mpep and Dpep, respectively: negative controls, TM^LDLR^ = the TM domain of LDL receptor and Xpeps specific for different OR homodimers; Fig. 5e).

Particularly, TM1^MOR^ reduced both MOR and DOR homodimers, whereas Mpep reduced MOR homodimers and not DOR homodimers and Dpep reduced DOR homodimers and not MOR homodimers. Taken together, we conclude that the interaction specificity of TM1^MOR^ is limited as shown from its ability to interact with both MOR and DOR (Fig. 5b, e). The results showing that TM1^MOR^ suppresses both MOR-MOR and DOR-DOR homodimerization indicate that TM1^MOR^ is involved in both MOR homodimerization and MOR-DOR heterodimerization (also refer to Figure 5C in the companion paper for the involvement of TM1^MOR^ for MOR-DOR heterodimerization). These results further indicate that TM1^MOR^ might not be useful as a specific blocker for MOR homodimerization or MOR-DOR heterodimerization.

### No major effects of Xpeps on the agonist-induced OR internalization

We specifically blocked OR homodimers by the addition of Xpeps (FAM-Xpep-TATs), and examined the effects on the agonist-induced OR internalization. Since all experiments were performed at expression levels of ≈1 OR spot/µm^2^, the percentages of the OR protomers existing as homodimers are 10-21% without any addition, 3.2-4.4% after the addition of FAM-Xpep-TATs, and 10-18% after the agonist addition, which is decreased to 2.9-8.6% after the further addition of FAM-Xpep-TATs (see the column of 1 copy/µm^2^ in Fig. 4i).

OR internalization might be involved in opioid side effects and tolerance development. Here, it was monitored by using the membrane-impermeable fluorescence quencher, Mn(III) meso-tetra(4-sulfonatophenyl)porphine (Mn^3+^-TSP). This quencher only suppresses fluorescence emission from the SNAP-Surface 549 dye on the SNAPf-OR on the cell surface, but not that in the cytoplasm.^60^

Accordingly, by subtracting the signal intensity after the quencher addition from that before the addition, we evaluated the percentages of OR molecules remaining in the PM (Fig. 6a and Supplementary Fig. 6).

**Fig. 6:**
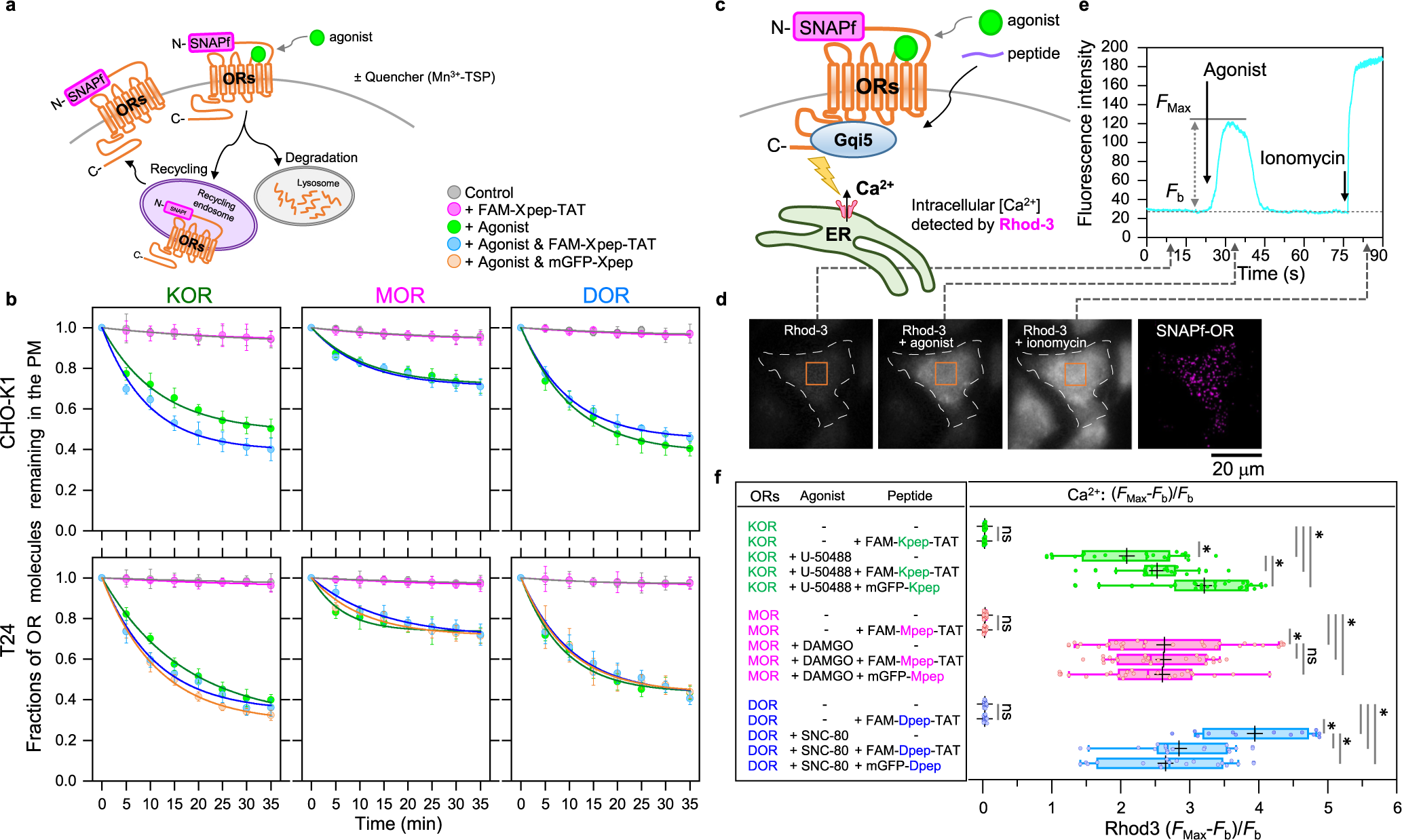
Xpeps hardly affect OR internalization in both the presence and absence of the agonist, whereas they enhanced, did not affect, or reduced the Ca^2+^ mobilization induced by the KOR, MOR, and DOR agonists, respectively. **a** Schematic figure showing the experimental design for observing the internalization of OR (with possible recycling). Also see Supplementary Fig. 6. **b** Time-dependent decreases of SNAPf-ORs on the PM due to their internalizations in CHO-K1 cells (top) and T24 cells (bottom) (mean ± SEM; 10 cells for each condition), before and after the addition of 0.2 μM agonist in the presence and absence of ≈3 μM FAM-Xpep-TATs and ≈6 μM mGFP-Xpeps in the cytoplasm. Photobleaching of the fluorescent probe is negligible because only one frame was recorded every 5 min. The time courses could be operationally fitted by single exponential functions, y = C_*_exp(-t/*τ*_0_) + (1-C), providing the fractions of OR molecules with detectable (C) and non-detectable (1 - C) internalizations for the observations up to 35 min, as well as the residency times (*τ*_0_) for the internalized component. **c** Schematic figure showing the experimental design for observing the Ca^2+^ mobilization after the agonist stimulation. The Ca^2+^ mobilization was monitored by the Rhod-3 fluorescence intensity (see the caption to Supplementary Fig. 1a and Methods). **d, e** Typical fluorescence images of Rhod-3 in cells (**d**; orange rectangles = ROI) and the time-dependent changes of the Rhod-3 signal intensity in the ROI (**e**), showing the changes in the cytosolic Ca^2+^ concentration after the agonist stimulation. Cells with similar expression levels of ORs (≈1 fluorescent spot/μm^2^) were selected. **f** Ca^2+^ mobilization at 75 s after the agonist addition, parametrized by using [*F*_Max_-*F*_b_]/*F*_b_ (see **e**).

The time courses of the decrease in the numbers of ORs remaining in the PM during 35 min (due to their internalization) were examined in both the presence and absence of 0.2 µM agonists and ≈3 µM Xpeps in the cytoplasm (Fig. 6b; Table 1 and Supplementary Table 1). All the time-course data obtained under various conditions could be fitted with a single exponential function plus a constant. The constant provides the OR fraction whose internalization is undetectable by the 35-min observations, and the exponential decay constant provides the residency lifetime in the PM for the OR component whose internalization was detectable by this observation scheme (Table 1 and Supplementary Table 1). Without agonist, the three ORs all exhibited non-internalized fractions in the range of 91-97% (for an observation period of 35 min). The internalized molecules found during the 35-min observations showed characteristic dwell lifetimes (inverse of internalization rates) between 24 and 45 min. The homodimer-blocking peptides, FAM-Xpep-TATs, did not significantly affect the OR internalization (Fig. 6b and Table 1).

The addition of agonists employed in this investigation greatly increased the fractions of detectable internalizations and the internalization rates for all three ORs (Fig. 6b and Table 1), although it decreases the percentages of KOR protomers existing as homodimers from 21 to 10% but increases the percentages from 11 to 18% and 10 to 17% for MOR and DOR, respectively (Fig. 4i). The FAM-Xpep-TATs reduced the homodimer percentages to 2.9, 3.3, and 8.6% for the engaged KOR, MOR, and DOR, respectively (Fig. 4i). The internalization observations indicated that for the engaged KOR, the addition of FAM-Kpep-TAT induced only slightly faster internalization, whereas for the engaged MOR and DOR, no differences in internalization were detectable after the addition of FAM-Mpep-TAT and FAM-Dpep-TAT (Fig. 6b and Table 1).

The lack of effect (or only the slight effect) of FAM-Xpep-TATs on the agonist-induced OR internalization is probably because monomers and homodimers are internalized at about the same rate. This might be induced if the internalization rate of homodimers as an internalization unit (two protomers together as a homodimer) might become about half of that of monomers or if the complex processes involving GRK and β-arrestin 2^29, 61, 62, 63^ might remove the ORs bound by these proteins from the monomer-dimer equilibrium in the bulk PM.^87–90^ However, due caution in the interpretation of these data is required because the Xpep-bound C-terminal domain might inhibit or enhance the interaction with GRK and β-arrestin 2 (for the Gqi5 binding, see the next subsection).

In addition to CHO-K1 cells, T24 cells were employed for these experiments because the expression level of β2-arrestin in T24 cells is higher than that in CHO-K1 cells,^64^ and thus T24 cells might undergo more active internalization. However, T24 cells exhibited internalization behaviors quite similar to those of CHO-K1 cells (Fig. 6b; Supplementary Tables 1 and 3).

### Xpeps modulate KOR and DOR signals

The agonist-induced signal downstream from the OR was examined by a widely used method for Gi-coupled GPCRs, employing the artificial G protein Gqi5 (Fig. 6c-6e; refer to the caption to Supplementary Fig. 1a and Methods).^65, 66, 67^ The addition of the homodimer-blocking FAM-Xpep-TATs alone (thus the monomerization of ORs alone) did not induce any detectable Ca^2+^ mobilization (Fig. 6f and Table 1). Meanwhile, the respective agonists triggered Ca^2+^ mobilization. The effects of the homodimer-blocking peptides on the agonist-induced Ca^2+^ mobilization were complex (both FAM-Xpep-TATs and mGFP-Xpeps produced similar effects for all three ORs). In cells expressing KOR, MOR, or DOR, the homodimer-blocking peptides (OR monomerization) enhanced, did not affect, and reduced the agonist-induced Ca^2+^ mobilization, respectively (Fig. 6f and Table 1). These results indicate that KOR and DOR monomers trigger higher and lower signals than their respective homodimers, whereas MOR monomers and homodimers induce the downstream signals at similar levels (without influencing the agonist-induced internalization of all ORs; Fig. 6b). In short, homodimerization suppresses the ligand-induced signaling of KOR and enhances the ligand-induced signaling of DOR.

It is possible that these results might be induced by the interfering effect of the bound peptide on the binding of Gqi5 to ORs (Gqi5’s binding site to ORs is the same as the original Gαi) because the Gi binding to class A GPCRs involves various intracellular domains.^62, 63, 68, 69, 70, 71^ However, since the Halo-tag protein bound to the C-termini of ORs do not affect OR- induced Ca^2+^ responses (Supplementary Fig. 1b of the companion paper) and since the peptide binding induces distinct effects on the three ORs, we think it likely that the effect of homodimer blocking peptides on the agonist-induced Ca^2+^ signal can be interpreted based on its effect on OR monomerization.

Furthermore, these results suggest that the homodimer-blocking peptide-TATs could be used as reagents (drugs) to enhance or suppress the agonist-induced G-protein-biased cellular responses for KOR and DOR, respectively, without affecting their internalization (Fig. 6b). However, note that these results are correct only for the employed agonists, and the results would be agonist dependent.

## Discussion

For the first time, the three critical parameters describing the homodimer-monomer dynamic equilibrium (*K*_D_, *K*_on_, and *K*_off_) have been determined for the OR. This was done for the three classical ORs before and after the addition of a representative agonist for each OR (Fig. 2b and 4i). The results unequivocally demonstrated that the three classical ORs all form transient metastable homodimers at physiological expression levels (0.3-10 copies/µm^2^) at 37°C, and predicting that 3.4-59% of OR protomers would exist as homodimers in various tissues both before and after the agonist binding (Fig. 2d). Accurate evaluation of the dimer dissociation rate constant (*K*_off_) in the live-cell PM using single-molecule colocalization data became possible by developing a theory (Supplementary Note 1), and the dimer-monomer dissociation equilibrium constant (*K*_D_) could be obtained much more readily compared with the method we developed previously.^37^ These theories/methods enabled accurate quantitative evaluations of *K*_off_ and *K*_D_, and would be applicable to evaluate the dimerization kinetics and equilibrium of any membrane molecules, as exemplified by their application to OR heterodimerization described in the companion paper.^38^ Accordingly, they represent major developments for studies of molecular interactions in the PM in general.

A previous single-molecule imaging study of neurotensin receptor 1, a GPCR, in reconstituted membranes containing only this receptor and a phospholipid revealed that it forms homodimers. This result demonstrated that neurotensin receptor 1 homodimerization occurs without the involvement of any other proteins^21^ (however, see^14^), and suggested that ORs might also form homodimers in a similar manner.

Importantly, all of these homodimers are forming and dispersing continually, with lifetimes shorter than 0.2 s at 37°C (Fig. 1d). When the homodimers dissociate into monomers, they will again form homodimers with the same and other partner molecules. Longer homodimer lifetimes on the order of 0.5-2 s or even 15 s have been reported, but they were observed at lower temperatures such as RT and 20°C.^33, 34, 35^ How the limited dimer fractions and lifetimes modulate the downstream signaling and internalization is extensively discussed in the companion paper (refer to Figure 7C and related main text in the companion paper).

Specific 9-26 residue amino-acid sequences in the near-C-terminal cytoplasmic domains, which lack sequence similarities among the three ORs, are involved in the distinct homodimerizations of all three classical ORs (Fig. 3b, c, and 4), and do not participate in heterodimerization (Fig. 2a and 3a in the companion paper). In addition to localized electrostatic interactions between charged amino acid residues, the overall structures of the cytoplasmic C-terminal domains, and particularly the intrinsically disordered structures of the C-terminal regions, might be important for homodimerization.^56^ The enhancement of homodimerization by the intrinsically disordered region might not be due to liquid-liquid phase separation, but rather to its ability to flexibly adopt various structures, bringing specific binding sites into positions and orientations for optimal interactions.

In the literature on homo- and hetero-dimerization of GPCRs, including ORs, the TM domains have been proposed or found to be responsible for homodimerization.^21, 26, 49, 50, 52, 53^ In the present study, we experimentally and directly confirmed this using TM1^MOR^, but we also found that TM1^MOR^ binding to ORs is likely to be quite non-specific, as found by its binding to DOR as well as MOR, and its suppressive effect on both DOR and MOR homodimerization. Blocking of DOR homodimerization of TM1^MOR^ suggests that it is also involved in DOR-MOR heterodimerization. Furthermore, we found that Xpeps failed to reduce the OR homo-colocalization indexes to the level of a monomer control molecule TM^LDLR^. Therefore, we propose that TM-TM interactions enhance both the homo- and hetero-dimerizations of ORs in less specific manners, whereas the cytoplasmic C-terminal domain interactions provide the specificities for OR homodimerization.

To the best of our knowledge, the Xpep-TATs developed here are the first peptide- based reagents that specifically reduce GPCR homodimerization. These homodimer-blocking peptides can become important tools to dissect the functions of OR monomers and homodimers. The use of the Xpeps revealed that DOR and KOR homodimers, but not MOR homodimers, induce G-protein-dependent signals differently from their respective monomers (Fig. 6f), although both monomers and homodimers are internalized at the same rate before and after agonist addition (Fig. 6b). These results suggest that the C-terminal domain conformations of the DOR and KOR homodimers might undergo allosteric changes upon homodimerization, affecting their G-protein-based signaling. The possibility of the involvement of allosteric conformational changes of ORs upon dimerization will be further discussed in the companion paper (allosteric effects might be more important for DOR-MOR heterodimers).

These homodimer blocking peptides could become the basis for designing modulators and potentiators in opioid therapy to enhance the efficacy and suppress side effects such as tolerance development, particularly for the opioids targeting KOR and DOR. The development of agonists for favorable biased signaling is important,^72, 73, 74, 75^ but the modulation of the OR homodimerization would provide another means to produce biased signaling. Since the Xpep- TATs are soluble and yet membrane-permeable (Fig. 4e-h) and thus they are likely to pass the blood-brain barrier, they could be readily delivered via intravenous injections or nasal sprays. In addition, Xpep-TATs would not compete with agonist binding because they bind to the cytoplasmic C-terminal domains. Taken together, our findings reported here have greatly extended our knowledge of ORs’ metastable homodimers and their formation mechanisms and functions, which can inform novel GPCR drug development strategies of modulating homodimerization.

## Methods

### cDNA construction

All of the newly generated cDNA constructs and other constructs obtained from outside sources, including gifts and constructs from commercial sources, were sequenced to examine their exact DNA sequences. The cDNA encoding rat MOR tagged with GFP was a gift from Dr. R. Schülz of the University of Münich, Germany.^76^ The cDNA encoding rat KOR and DOR was a gift from Dr. Hiroshi Takeshima of the Kyoto University.^77^ The mCherry was a gift from Prof. R. Y. Tsien of the University of California San Diego.^78^ The cDNA encoding CD47 was a gift from Eric C Brown of Genentech.^79^ The cDNA encoding CD28 was a gift from Simon J Davis of University of Oxford.^80^ The cDNAs encoding SNAPf and mGFP (A206K) were obtained from Promega and Clontech, respectively. To generate plasmids for expressing SNAPf-ORs in CHO- K1 cells, the tag protein SNAPf was attached to the N-terminus of the ORs, an additional signal sequence of interleukin 6 was attached to the N-terminus of the tag protein, and a 21 amino-acid linker (SGGGSGG x3) was inserted between the ORs and the tag protein. A Site- Directed Mutagenesis Kit (New England Biolabs) was used to generate the cDNAs encoding the OR point mutants.

### Cell culture, transfection, and microscope observations

CHO-K1 cells (Dainippon Pharma) and T24 cells (gift from Prof. M. Sokabe of Nagoya University),^81^ confirmed free of mycoplasma contamination by MycoAlert (Lonza), were routinely cultured in Ham’s Nutrient Mixture F12 (Sigma-Aldrich) supplemented with 10% (v/v) fetal bovine serum (FBS, Life Technologies), 100 units/ml penicillin, and 0.1 mg/ml streptomycin, at 37°C under a 5% CO_2_ atmosphere in the incubator. Transfection of CHO-K1 cells with the cDNAs of interest was performed by electroporation, according to the manufacturer’s instructions (4D-Nucleofector, Lonza; SF Cell Line solution and program CHO- K1 for CHO-K1 cells). The transfected cells were seeded in glass-base dishes (35-mm in diameter with a 12-mm diameter glass window, 0.15-mm-thick glass; Iwaki, Tokyo; 2×10^5^ cells/dish) and cultured for 24-48 h before fluorescence microscopy observations. All microscope observations were performed at 37°C by placing the entire microscope, except for the far ends of the excitation arms and the detection arms, in a home-built microscope environment chamber made with thermo- and electric-field-insulating plastic sheets and equipped with four heating circulators (SKH0-112-OT, Kokensya, Tokyo, Japan). The Ham’s F12 medium used for microscope observations was free of sodium bicarbonate and phenol red, and buffered with 2 mM N-[tris(hydroxymethyl)methyl]-2-aminoethanesulfonic acid (TES, Sigma-Aldrich) at pH 7.4 (called Ham’s F12 observation medium).

### Cell treatments with agonists and FAM-Xpep-TATs

U-50488, DAMGO, and SNC-80 (Sigma-Aldrich), agonists for KOR, MOR and DOR, respectively, were applied to the cells in the same way. The agonists were dissolved in DMSO (2 mM), and then diluted with Hanks’ balanced salt solution (HBSS, Nissui) buffered with 2 mM TES at pH 7.4 (T-HBSS), at a final concentration of 200 µM. The agonist solution (1 µl) was added to the cells in 1 ml Ham’s F12 observation medium (a final concentration of 200 nM) at 37°C.

For the cellular incorporation of FAM-Xpep-TATs (custom-synthesized by Cosmo-Bio), the cells were first incubated with 150 µM pyrenebutyrate (Sigma-Aldrich) in T-HBSS at 37°C for 5 min, and then 2 mM FAM-Xpep-TAT in T-HBSS was added at a final peptide concentration of 20 µM. After an incubation at 37°C for 10 min, the cells were washed three times with T-HBSS, and then fresh Ham’s F12 observation medium was added to the cells.

The presence of FAM-Xpep-TAT in the cytoplasm was confirmed by the addition of the membrane-impermeable quencher, trypan blue.^82^

### Fluorescence labeling of ORs

The SNAPf-tagged wild-type and mutant ORs expressed in the PM (SNAPf tag located at the extracellular N-terminus) were covalently conjugated by simultaneously incubating the cells with two fluorescent SNAP ligands, SNAP-Surface 549 (New England Biolabs) and SNAP- CF660R (Shinsei Kagaku), both at 300 nM, in the growth medium at 37°C in the CO_2_ incubator for 30 min. The cells were washed three times with fresh medium (5-min incubation each time), and then the Ham’s F12 observation medium was added. The labeling efficiency was determined by using the SNAPf-CD47-mGFP protein expressed in the PM. CD47 is a monomeric 5-pass transmembrane protein expressed in the PM.^83^ We could not use ORs for this purpose because they form transient dimers, and thus employed a monomeric CD47 (Supplementary Fig. 1c, d).

The labeling efficiency (the percentage of the SNAPf-tag protein labeled with the fluorescent SNAP ligand) was determined in the following way. The percentage of fluorescent mGFP spots of SNAPf-CD47-mGFP that was colocalized by the SNAP-dye spots were measured as a function the concentration of the SNAP-dye-conjugated ligand in the medium, after a 30 min incubation at 30°C. Note that not every mGFP molecule was fluorescent, but since we examined whether the fluorescent mGFP molecule identified in the image was colocalized by a SNAP-dye spot, the labeling efficiencies of the SNAPf-protein tag can be evaluated without the influence of the fluorescence efficiency of mGFP. Then the efficiency at each fluorescent ligand concentration was plotted against the dye concentration (Supplementary Fig. 1e). We found that the plot could be fitted by a function of a single exponential function plus a constant, providing the saturating labeling efficiency (constant) and the characteristic labeling efficiency (exponential constant).

To obtain the labeling efficiency at a dye concentration of 300 nM for an incubation period of 30 min, the conditions used throughout this study, the determined exponential function was used for a concentration of 300 nM. The fluorescence labeling efficiencies under our normal observation conditions were found to be 60% and 61% for SNAP-Surface 549 and SNAP-CF660, respectively.

### Single fluorescent-molecule imaging

Fluorescently labeled ORs expressed in the bottom PM (the PM facing the coverslip) at fluorescent-spot number densities of 0.5 ± 0.25 spots/µm^2^ for each color (SNAP-Surface 549 and SNAP-CF660R; total spot number densities of 1.0 ± 0.5 spots/µm^2^; for conciseness, we describe the number densities as ≈1 spot/µm^2^ throughout this report) were observed at the level of single molecules at 37°C, using a home-built objective lens-type TIRF microscope constructed on an inverted microscope (Olympus IX-83), with a 100x, 1.49 numerical aperture (NA) objective lens, optimized for the present research based on the instrument used previously.^44, 84^ ORs tagged with fluorescent probes were excited with TIR illumination using the following power densities: SNAP-Surface 549 at 561 nm (Coherent OBIS 561-100 LS) at 0.35 µW/µm^2^; and SNAP-CF660R at 642 nm (Omicron LuxXPlus 640-140) at 0.52 µW/µm^2^. The dual-color images were separated by a dichroic mirror ZT405/488/561/640rpc- UF3 (Chroma) and projected into two detection arms with band-pass filters of 572.5-647.5 nm (ET600/50m; Chroma) for the SNAP-Surface 549 dye and 662.5-737.5 nm (ET700/75m; Chroma) for the SNAP-CF660R dye. In some experiments using mGFP-Xpeps and FAM-Xpep- TATs, an additional detection arm equipped with a dichroic mirror ZT405/488/561/640rpc- UF3 (Chroma) and a band-pass filter of 500.0-550.0 nm (ET525/50m; Chroma) was employed. The fluorescence signal in each channel was first detected and amplified by a two- stage microchannel-plate image intensifier (C9016-02MLP24; Hamamatsu Photonics), and the intensified image was projected onto the scientific CMOS camera (C11440-22CU; Hamamatsu Photonics), operated at 30 Hz, which was synchronized with the same intensifier-camera set(s) placed on another detection arm. The image sequences in each channel were superimposed after correction for spatial distortions, as described previously.^44^ The positions (x and y coordinates) of all of the observed single fluorescent-molecules were determined by an in-house computer program, as described previously.^85^

### Evaluating colocalization durations

The colocalization of two fluorescent molecules was defined as the event where two fluorescent spots, representing these molecules, become localized within 200 nm of each other, as described previously.^37, 44^ Briefly, in single-color experiments using SNAP-Surface 549 (Supplementary Fig. 2), a spatial cross-correlation analysis was employed.^37^ Using this method, we found that when two fluorescent spots, each representing a single molecule of SNAP-Surface 549, are located close together, the threshold distance for discriminating one or two spots occurred at 200 nm in the present experimental set up. Using this definition, colocalized trajectories were obtained and colocalization durations were estimated. In simultaneous two-color single fluorescent-molecule tracking experiments, using the dye pairs of SNAP-Surface 549 and SNAP-CF660R, the distance between the two molecules was directly measured from the locations (x, y-positions) of each molecule (with different colors). Even when examining pairs of different-colored molecules that are known to be truly associated, the probability of scoring the two molecules as associated is limited by the localization accuracies of each molecule and the accuracies of superimposing the two images. Based on the method developed previously^44^ and the accuracies determined here, we found that, for truly associated molecules, the probability of scoring the two molecules as associated increases to 99% when using the criterion that the molecules lie within 200 nm of each other. Therefore, we used this criterion as the definition of colocalization in simultaneous two-color single-molecule observations. This distance of 200 nm coincided with the definition of colocalization in single-color experiments. Due to this coincidence, in the present research, we defined the colocalization of two fluorescent molecules as the event where the two fluorescent spots representing these molecules become localized within 200 nm from each other.

Each time we found a green-magenta pair located within 200 nm (colocalization), we measured the duration in which their distances remained within 200 nm (colocalized duration) (for the results obtained by using two dye molecular species with different excitation/emission wavelengths, we call them green and magenta probes/movies for convenience in this report). After obtaining the colocalization durations for all of the colocalization events, we generated a histogram (distribution) of colocalization durations. The distribution of incidental colocalization durations was obtained by superimposing the magenta image sequences with the 180-degree rotated (doubly flipped) green image sequences (Supplementary Fig. 2d). For the precise analysis of these histograms, we first produced cumulative histograms and fitted them with a single exponential function (C_0_ – C_1_exp[-t/*τ*_1_]) or the sum of two exponential functions (C_0_ – C_1_exp[-t/*τ*_1_] - C_2_exp[-t/*τ*_2_]). The choice of the number of exponentials (including the cases of the sum of three exponential functions) was made based on Akaike’s and Bayesian information criteria (they always agreed). Based on these functions, the functions describing the original histograms were derived and overlaid on the histograms.

First, we found that the distribution of incidental colocalization durations could be fitted by a single exponential function, with a time constant representing the incidental colocalization lifetime *τ*_inci_. Second, the distribution of colocalization durations was obtained for correctly superimposed magenta and green image sequences. This distribution could be fitted by the sum of two exponential decay functions (*τ*_1_ and *τ*_2_; *τ*_1_ < *τ*_2_,). These results are consistent with the theory developed here that predicts the distribution of colocalization durations based on the diffusion equation (Supplementary Note 1). Since *τ*_1_ was almost the same as *τ*_inci_ (Fig. 1e), following the theory developed in Supplementary Note 1, *τ*_2_ provided the true homodimer lifetime (inverse of *K*_off_; after correction for the trackable duration lifetimes of the two fluorescent probes). The trackable duration lifetime is obtained from the distribution of trajectory lengths, which would represent how long single molecules can be continuously tracked under the influences of photobleaching and loss of signals due to blinking and the entrance of the molecules into the PM areas located farther from the PM. The distributions could be fitted as single exponential functions providing trackable duration lifetimes (*τ*_track_’s) of 16.3 ± 1.2 s (n = 500) for SNAP-Surface 549 and 7.8 ± 0.6 s (n = 400) for SNAP-CF660R bound to SNAPf-MOR expressed in CHO-K1 cells observed at 30 Hz (Supplementary Fig. 1f). The correction was made by using the equation:

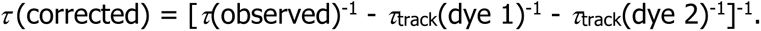

### Evaluating the colocalization index

For the quantitative evaluation of the extent of colocalization (representing both the frequency and lifetime of colocalization events) in simultaneous two-color single-molecule imaging movies, we defined a parameter called the colocalization index. This analysis method is essentially based on a pair cross-correlation analysis,^46^ and the detailed method used in this research is explained in Supplementary Fig. 2c-e and 3.

### Monte-Carlo simulations

For the detailed method for Monte-Carlo simulations to examine the theory to obtain *K*_D_ from the pair cross-correlation function (PCCF) (Supplementary Note 2), see the caption to Supplementary Fig. 3.

The colocalization index depends on the number density of fluorescent spots in the PM. In the present experimental observations, we selected the cells exhibiting fluorescent spot number densities in the PM of 0.5 ± 0.25 spots/µm^2^ for each color (SNAP-Surface 549 and SNAP-CF660R; total spot number densities of 1.0 ± 0.5 spots/µm^2^; for conciseness, we describe the number densities as ≈1 spot/µm^2^ throughout this report). These densities were found to be reasonable by the Monte-Carlo simulation results, which indicated that, for the *K*_D_ greater than 6.5 copies/µm^2^, the colocalization index does not vary sharply in the number density range of 0.5 - 1.5 spots/µm^2^ (refer to Supplementary Fig. 3c). At the same time, we developed the theory to evaluate the dimer-monomer dissociation equilibrium constant *K*_D_ from the PCCF and the total number of spots in the image (refer to Supplementary Note 2).

### Confocal imaging to study the effects of mGFP-Xpeps and FAM-Xpep-TATs on cells stably expressing ORs

CHO-K1 cells stably expressing ORs were transfected with mGFP-Xpeps. The cells were identified by the incubation with NucSpot Live 650 Nuclear Stain (Biotium), according to the protocol recommended by the manufacturer. After three washes with the complete medium, it was replaced with the Ham’s F12 observation medium. The incorporation of the FAM-Xpep- TATs in the cells was performed as described in a previous subsection “Cell treatments with agonists and FAM-Xpep-TATs”. Confocal fluorescence images were acquired on the same microscope station used for single-molecule imaging at 37°C, equipped with an Olympus SR10 spinning-disc confocal super-resolution unit (with a Plan-Apochromat 100x oil immersion objective; NA = 1.49).

GFP and FAM were excited at 488 nm and detected through a 505-530 nm band pass filter. The NucSpot Live 650 was excited at 642 nm and detected through a 662.5-737.5 nm long pass filter. The concentrations of cytoplasmic mGFP-Xpep and FAM-Xpep-TAT were evaluated using calibration curves, obtained by observing the purified EGFP protein (BioVision) and FAM-Xpep-TATs dissolved in Ham’s F12 observation medium at various concentrations, with a focus at 5 µm above the cover-glass surface (Supplementary Fig. 5b).

### Ca^2+^ mobilization assay

Since the OR expression level varies from cell to cell, the signaling process must be observed at the level of individual cells. The ORs are coupled to the inhibitory trimeric G protein Gi, which induces the decrease of the cytoplasmic cAMP concentration by inhibiting adenylyl cyclase. However, using the cells with low OR expression levels employed for single-molecule observations (≈1 fluorescent spot/µm^2^), the decrease of the cytoplasmic cAMP levels in individual cells from those in non-stimulated state was impossibly difficult to measure. Meanwhile, the increase of the cytoplasmic Ca^2+^ concentrations by the PLCβ activated by the Gq signaling pathway could be measured in individual cells.^65, 66, 67^ Therefore, in this assay, Gαq was modified so that OR could be coupled to Gαq. Namely, the C-terminal five amino-acid sequence of Gαq, ECGLY, was replaced with that of Gαi2, DCGLF (this chimeric protein is called Gqi5), because the Gα protein binds to the specific GPCR by way of its short C-terminal sequence.^86^ Therefore, we generated the CHO-K1 cell line stably expressing Gqi5, had it express the OR, and then observed the cytoplasmic Ca^2+^ mobilization upon agonist addition, using the Ca^2+^-sensitive dyes (Fig. 6c-f and Supplementary Fig. 1b). This method of using Gqi5 has been widely employed in the research of ORs^65^ and other Gi-coupled GPCRs.^66^ Briefly, CHO-K1 cells stably expressing Gqi5 were generated and transfected with the cDNAs encoding the wild-type and SNAPf-linked ORs. The Ca^2+^-sensitive dyes Fluo-4 AM (Dojindo) and Rhod-3 AM (Thermo Fisher Scientific) were employed. Rhod-3 was used for the experiments with mGFP-Xpeps and FAM-Xpep-TATs, and Fluo-4 was used for the experiments that did not employ these homodimer-blocking reagents. These AM dyes were incorporated in the cell, according to the manufacturers’ recommendations, using the following solutions: for Fluo-4, 4.6 µM Fluo-4 AM in T-HBSS containing 1.25 mM Probenecid (Dojindo) and 0.04 % (w/v) Pluronic F127 (Dojindo), and for Rhod-3, 10 µM Rhod-3 AM in T-HBSS containing 2.5 mM Probenecid and 1× PowerLoad^TM^ (Thermo Fisher Scientific). These loading solutions (2 ml) were added to the cells and incubated in the dark at 37°C for 30 min, and then the cells were washed three times with T-HBSS.

For the observations of the Ca^2+^ mobilization downstream of the SNAPf-tagged ORs, we selected cells expressing SNAPf-tagged ORs bound by SNAP-Surface 549 (or SNAP-CF660R for experiments using Rhod-3) at number densities of ≈1 fluorescent spot/µm^2^ in the basal PM, using single-molecule detection (TIRF illumination) in the 561-nm channel (642-nm channel when we employed Rhod-3). These cells were then observed by epifluorescence illumination using the 488-nm channel to monitor the Fluo-4 signal (561-nm channel to observe the Rhod-3 signal and 488-nm channel for mGFP-Xpeps and FAM-Xpep-TATs).

Agonist stimulation was performed by adding the DMSO solutions of agonists at a final concentration of 200 nM. To determine the saturation levels of the fluorescence signal intensity at higher concentrations of Ca^2+^, 1 µM (final concentration) ionomycin (Wako) was added (which would increase the intracellular Ca^2+^ concentration to that outside the cells [1.3 mM]). The Fluo-4 and Rhod-3 image sequences were analyzed using the ImageJ software.

For the comparison of the functions of the SNAPf-ORs with those of the wild-type ORs, we hoped to compare the cells expressing the SNAPf-ORs at the levels of ≈1 spot/µm^2^ in the basal PM with the cells expressing the wild-type ORs at levels comparable to or higher than those of SNAPf-ORs, because the cells expressing the wild-type ORs should serve as the positive controls. For this purpose, the expression levels of wild-type ORs were monitored by using cells transfected with the cDNA linking the wild-type OR sequence to the mCherry sequence via the self-cleavable 2A linker sequence (mCherry-2A-OR). This way, mCherry is released from the OR into the cytoplasm at the ER, and the wild-type OR is then transported to the PM. The expression of wild-type OR was detected by the presence of mCherry in the cytoplasm using the epi-illumination at 561 nm (sensitivities much lower than single-molecule imaging, showing the presence of rather high concentrations of mCherry). Therefore, the expression of the wild-type OR used in this study is considered higher than that of SNAPf- ORs.

### Software and statistical analysis

The microscope station that combined a single-molecule imaging system and a super- resolution confocal microscope was controlled by in-house LabVIEW2018-based software, and the single-molecule movie acquisitions were performed using the MCR software (Hamamatsu Photonics) for Windows. All statistical analyses for in vitro experiments were performed with Tukey’s multiple comparison test except for the analysis of the colocalization lifetime data, which was performed with the Brunner-Munzel test, using OriginPro 2019b for Windows and RStudio 1.2.1335 for Windows. P values less than 0.05 were considered statistically significant. The confocal images were processed and analyzed using Image J for Windows.

Curve fitting was performed by OriginPro 2019b for Windows. The simulation study was performed using the in-house software based on MATLAB 2019a for Windows.

### Data availability

All data generated or analyzed for this study are available within the paper and its associated Supplementary Information. Any additional information required to reanalyze the data reported in this paper is available from the lead contact upon reasonable request.

Superimpositions of image sequences obtained in two colors and tracking single-molecules for in-vitro observation data were performed using C + +-based computer programs, produced in-house^87^ and based on the well-established approaches. Further information regarding the experimental design may be found in the Nature Research Reporting Summary.

### Code availability

Source codes for this have been integrated into a large, complex software package. While the software package cannot be extracted in a useful way, the entire software is available from the lead contact upon reasonable request, who will provide guidance on how to use it, as its manual and comments are written in Japanese.

## Acknowledgements

We thank Profs. H. Takeshima of Kyoto University, R. Schülz of the University of Münich, R. Y. Tsien of the University of California San Diego, Simon J. Savis of the University of Oxford, Michiyuki Matsuda of Kyoto University, and M. Sokabe of Nagoya University, School of Medicine, for their kind gifts of cDNAs encoding rat KOR and DOR, that encoding rat MOR, that encoding mCherry, that encoding CD28, plasmids of pPBpuro and pCMV-mPBase, and the human T24 epithelial cell line, respectively. We also thank Ms. Irina Meshcheryakova for technical help in preparing the cDNAs. This work was supported in part by Japan Society for the Promotion of Science (JSPS) Grants-in-Aid for Scientific Research (Kiban A to A.K. [21H04772], Kiban S to A.K. [16H06386], Kiban B to T.K.F. [16H04775, 20H02585], Kiban C to R.S.K. [17K07333], Wakate B to R.S.K. [26870292], Wakate to T.A.T. [21K15058], and Challenging Exploratory Research to T.K.F. [18K19001] and A.K. [22K19334]), a Grant-in-Aid from the Ministry of Education, Culture, Sports, Science and Technology of Japan (MEXT) for Transformative Research Areas (A) to T.K.F. (21H05252), and a JST grant ACT-X to T.A.T. (JPMJAX211B). WPI-iCeMS of Kyoto University is supported by the World Premiere Research Center Initiative (WPI) of MEXT.

## Contributions

P.Z., R.S.K., and A.K. conceived and formulated the project. A.K., T.A.T., T.K.F., and P.Z. developed the simultaneous dual-color single-molecule tracking station. P.Z., T.A.T., R.S.K., and A.K. designed biological experiments, and P.Z., with help from T.A.T., performed virtually all of the biological and microscopic experiments. T.K.F. developed and improved the in-house image analysis software. S.P. developed the theory for evaluating the *K*_off_ from the single- molecule experimental data (Note S1). K.M.H., T.A.T., and A.K. developed the colocalization index method, and T.A.T., Z.K., and A.K. developed the theory for linking the PCCF to the dimer dissociation equilibrium constant and performed Monte-Carlo simulations to prove the theory (Note S2). A.A. performed the OR sequence analysis to detect intrinsically disordered regions. A.K., P.Z., and T.A.T. wrote the manuscript, and all authors discussed the results and participated in the manuscript revisions.

