## Supplementary Information for "Single-molecule detection of transient dimerization of opioid receptors 1: Homodimers’ effect on signaling and internalization"

### Supplementary Materials

Captions to Supplementary Videos 1 and 2

Supplementary Figures 1-6

Supplementary Tables 1-3 (Supplementary Table 1 is an Excel table)

Supplementary Note 1

Supplementary Note 2

Supplementary References

### Captions to Supplementary Videos 1 and 2

#### Supplementary Video 1

Simultaneous dual color imaging of single SNAPf-KOR molecules (≈0.5 copies/$\mu m^2$) in the PM at 37°C, recorded at video rate for 100 frames (3.3 s) and replayed at a 3x-slowed rate from real time. The movie on the right shows the results of the automatic detection of colocalized spots (yellow squares) in the movie on the left.

#### Supplementary Video 2

A typical colocalization event of two single SNAPf-KOR molecules in the PM (yellow circles) shown by expanded image frames, replayed at a 30x-slowed rate from real time.

### Supplementary Figures

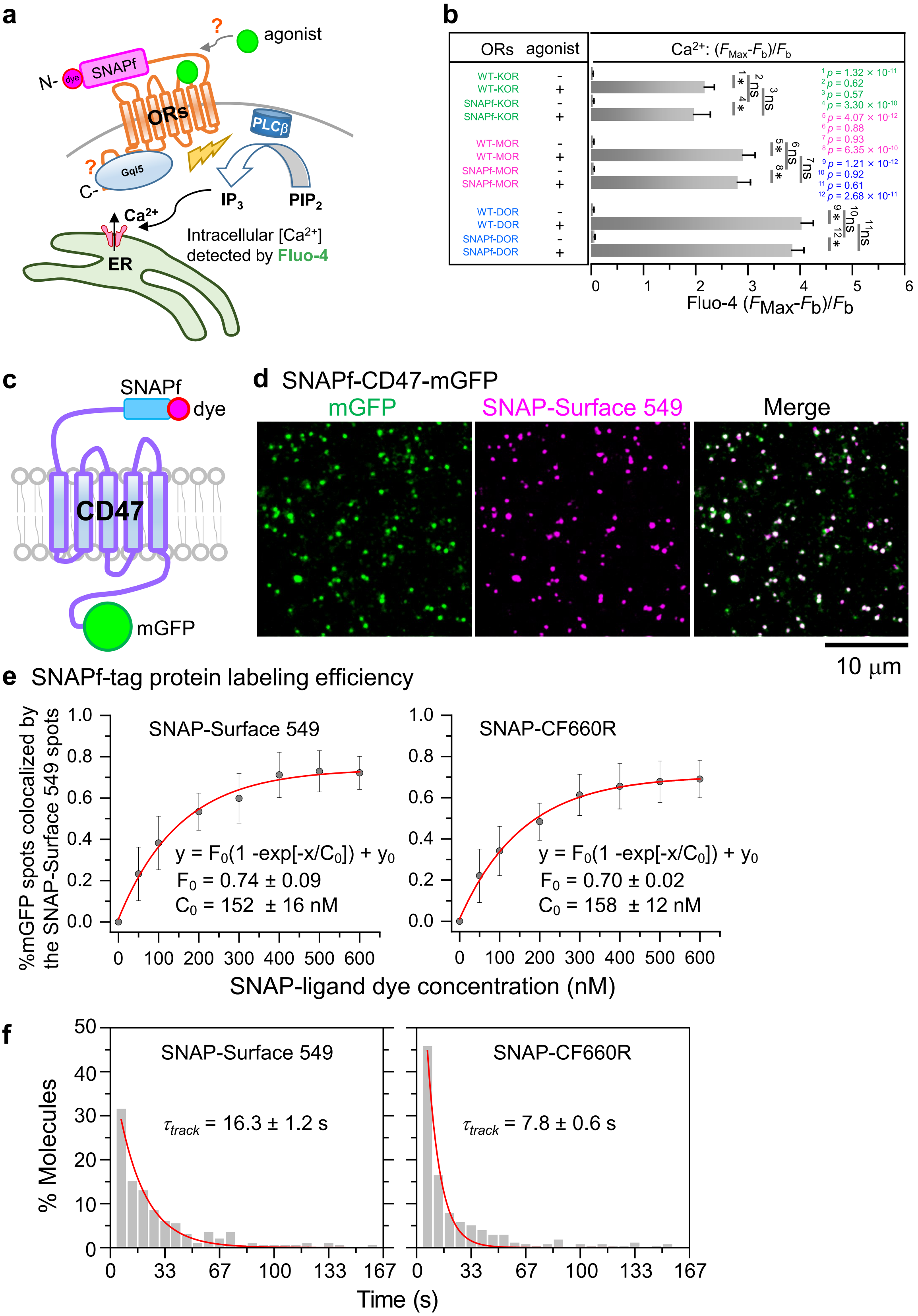


**Supplementary Fig. 1. Function test of the SNAPf-tagged ORs, labeling efficiencies of the SNAPf-tag with SNAP-Surface 549 and SNAP-CF660R, and photobleaching lifetimes of these dye molecules.**

**a.** Schematic figure showing the experimental design for testing whether the ORs conjugated with the tag protein SNAPf at the N-terminal ectodomain can function as well as the non-tagged ORs, by observing $Ca^{2+}$ mobilization (see Methods for further details).

CHO-K1 cells stably expressing Gqi5, the G$\alpha$q with the final five amino-acid sequence (ECGLY) replaced by that of Gαi2 (DCGLF),[1-4] were generated and then transfected with the cDNA encoding the SNAPf-linked OR or non-tagged OR conjugated to mCherry by way of the 2A-peptide linker (mCherry-2A-[non-tagged]OR). Cells expressing the SNAPf-OR in the range of 0.5 – 1.5 spots/$\mu m^2$ and mCherry (to identify cells expressing non-tagged OR, using epi-illumination) were employed.

The $Ca^{2+}$ mobilization was monitored by the fluorescence intensity of the $Ca^{2+}$-sensitive fluorescent dye Fluo-4. For the agonist stimulation, 0.2 μM U-50488, DAMGO, and SNC-80 were used for KOR, MOR, and DOR, respectively.

**b.** Definition of the measure of intracellular $Ca^{2+}$ mobilization employed in the work reported in this and companion papers. The $Ca^{2+}$ mobilization was parametrized by using $[F_{Max}-F_b]/F_b$, where $F_{Max}$ is the maximal Fluo-4 signal intensity within 75 s after the addition of the stimulants and $F_b$ is the baseline intensity (Fig. 6e).[5]

**c**. Evaluation method for determining the labeling efficiency of SNAPf-tag attached to the extracellular N-terminus of a TM protein. Since ORs form homodimers, their labeling efficiencies were difficult to evaluate, and therefore, a monomeric TM protein CD47 was employed,[6] after linking the SNAPf-tag and mGFP-tag to its N- and C-termini, respectively (SNAPf-CD47-mGFP expressed in the PM).

**d.** Typical simultaneous two-color, single-molecule images of SNAPf-CD47-mGFP in the PM after the incubation with 0.6 μM SNAP-Surface 549-conjugated ligand at 37°C for 30 min (among 20 cells).

**e.** Determination of the labeling efficiency, i.e., the percentage of the SNAPf-tag protein labeled with the fluorescent SNAP ligand: SNAP-Surface 549 (left) and SNAP-CF660R (right) (see Methods). Each datapoint shows the percentage of fluorescent mGFP spots of SNAPf-CD47-mGFP that were colocalized by the SNAP-dye spots (mean ± SEM for n=20 cells), and these datapoints were plotted as a function of the dye concentration in the incubation medium (30-min reaction time). The plot could be fitted by a function of a single exponential function plus a constant (values in the figure panels). Note that not every mGFP molecule was fluorescent, but since we examined whether the identified fluorescent mGFP molecule in the image was colocalized by a dye spot, the labeling efficiency of the SNAPf-protein tag can be evaluated without the influence of the fluorescence efficiency of mGFP. To obtain the labeling efficiency at a dye concentration of 300 nM, the conditions used throughout this study, the determined exponential function was used for a concentration of 300 nM. The fluorescence labeling efficiencies under our normal observation conditions were found to be 60% and 61% for SNAP-Surface 549 and SNAP-CF660R, respectively.

**f.** Distributions of the trackable duration times (trajectory lengths) of SNAP-Surface 549 and SNAP-CF660R bound to SNAPf-MOR expressed at densities of ≈0.1 spots/μm$^2$ in CHO-K1 cells observed at 30 Hz. The decay time constants obtained by single exponential fitting provided the trackable duration lifetimes of the two fluorescent probes of 16.3 ± 1.2 for SNAP-Surface 549 and SNAP-CF660R for 7.8 ± 0.6 s, respectively, under our experimental conditions. The photobleaching lifetimes for these fluorescent dye molecules adsorbed on the cover glass of the glass-base dish were found to be 36.6 ± 1.0 s (n = 567) for SNAP-Surface 549 and 15.1 ± 0.15 s (n = 468) for SNAP-CF660R.

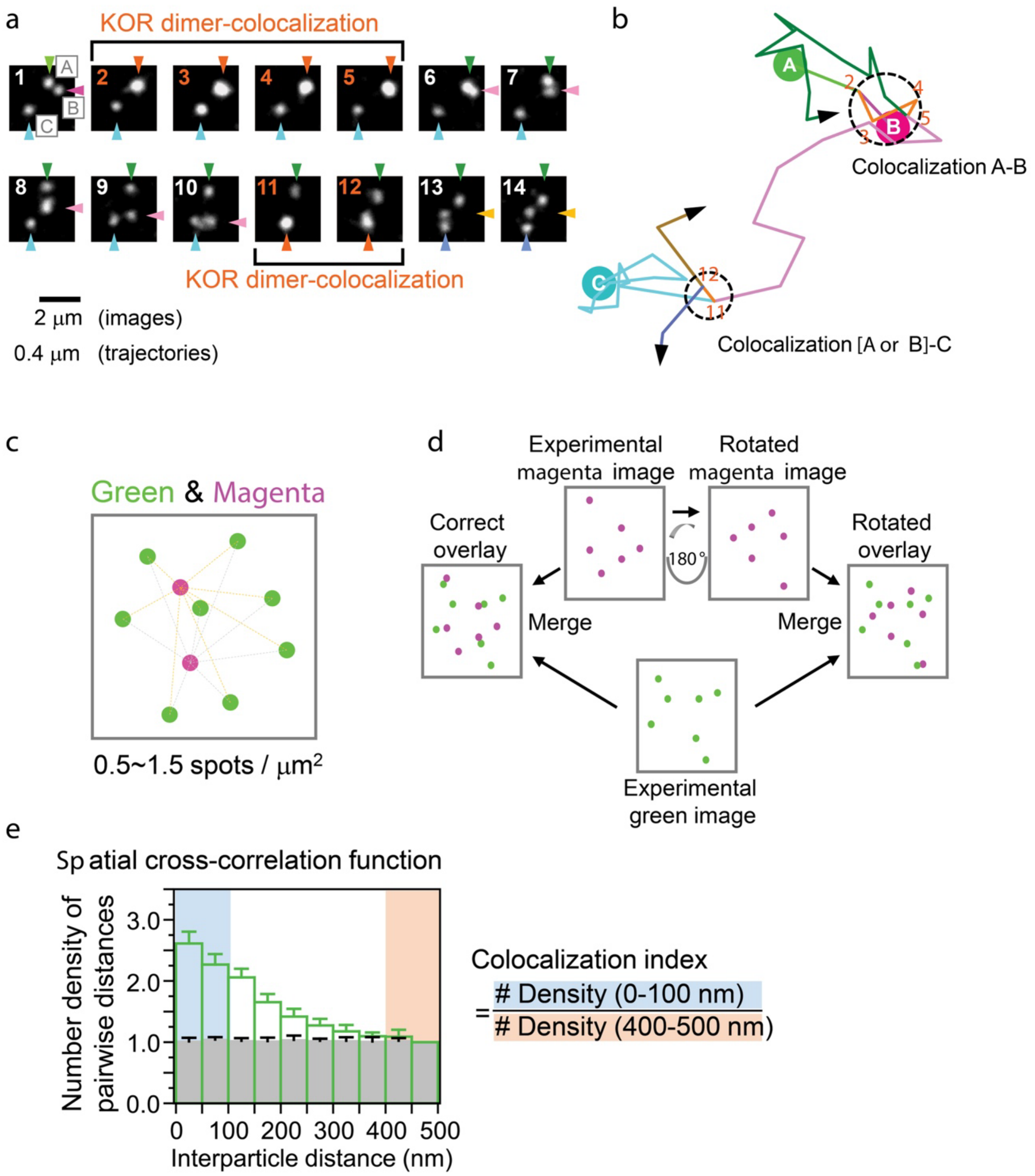


**Supplementary Fig. 2. Transient homodimerization events of KOR in the PM detected by single-color, single-molecule imaging, as done previously,[7-8] and definition of the colocalization index for the results of simultaneous two-color single-molecule imaging of ORs based on the spatial pair cross-correlation function (PCCF).**

**a** and **b.** Single-color, single-molecule imaging experiments showed that the homodimerization lifetime is ≈0.1 s, consistent with the simultaneous two-color results: The figure here shows typical KOR homodimer formation, dissociation, homodimer reformation with a different partner molecule, and then dissociation. Typical image sequence (every 33 ms) (**a**) and trajectories (**b**) of three diffusing SNAPf-KOR molecules (tagged with SNAP-Surface 549) undergoing two transient homodimerization events. In **b**, circles labeled A, B, and C indicate the locations of the three molecules in the first video frame (see the fluorescent spots in **a**). After the first colocalization of molecules A and B lasting for 133 ms (frames 2-5; see the numbers indicated on the trajectory), the second colocalization of the [A or B] molecule with the C molecule started after 333 ms (11th frame), and lasted for 66 ms (frames 11 and 12). Due to the single-color labeling, after the first colocalization event, we cannot tell which molecules are A and B, and thus we state molecule [A or B].

**c, d, e**. Using simultaneous two-color single-molecule imaging of ORs (green and magenta), we obtain the colocalization index based on the spatial pair cross-correlation function (PCCF).

**c.** First, a region of interest (ROI) with a rectangular (generally a square of 200x200 pixels) shape greater than 10 μm on each side is selected (to avoid the effect of edges, employing an ROI of this size or greater is useful). The ROIs with total spot number densities in the range of 0.5-1.5 spots/$\mu m^2$ are selected (see Supplementary Fig. 3 and Supplementary Note 2).

**d.** As a control for incidental colocalization, the green image is superimposed on the 180 degree-rotated magenta image (green and magenta are interchangeable), which is called the "180°-rotated overlay" (vs. the correct overlay).

**e**. Definitions of the PCCF in the histogram format and colocalization index. The distances between all pairs of green and magenta spots in the ROI in the simultaneously obtained video frames are measured, and this is repeated for all of the video frames of a single movie. Using this set of interparticle distances, the distribution of the number of pair distances is obtained. Throughout this research, we employed a bin size of 50 nm and a maximal distance of 500 nm. This distribution is then normalized for the unit area (i.e., the number densities at a given distance) and by the value obtained for the bin of 450-500 nm. This is the PCCF in the histogram format employed in this study. This process is repeated for all the movies to be analyzed, providing the mean ± SEM for each point (the height of each bar in the histogram) (green histogram). Because the histogram for each movie (ROI) is normalized by the value for the 450-500 nm bin, the value there is always 1 without SEM. This is repeated for the 180°-rotated control overlays (grey histogram). The histograms shown here are the same as those shown in Fig. 1b.

The colocalization index is defined as [the mean of the PCCF value for 0-50 and that for 50-100 nm; shown by blue highlighting] divided by [the mean of the PCCF value for 400-450 and that for 450-500 nm; shown by orange highlighting]. When no significant colocalization exists, the colocalization index is 1, and with an increase of colocalizations, the index increases. For the actual functional form describing the PCCF based on the number densities of spots in the images and the dimer-monomer dissociation equilibrium constant $K_D$, see Supplementary Note 2.

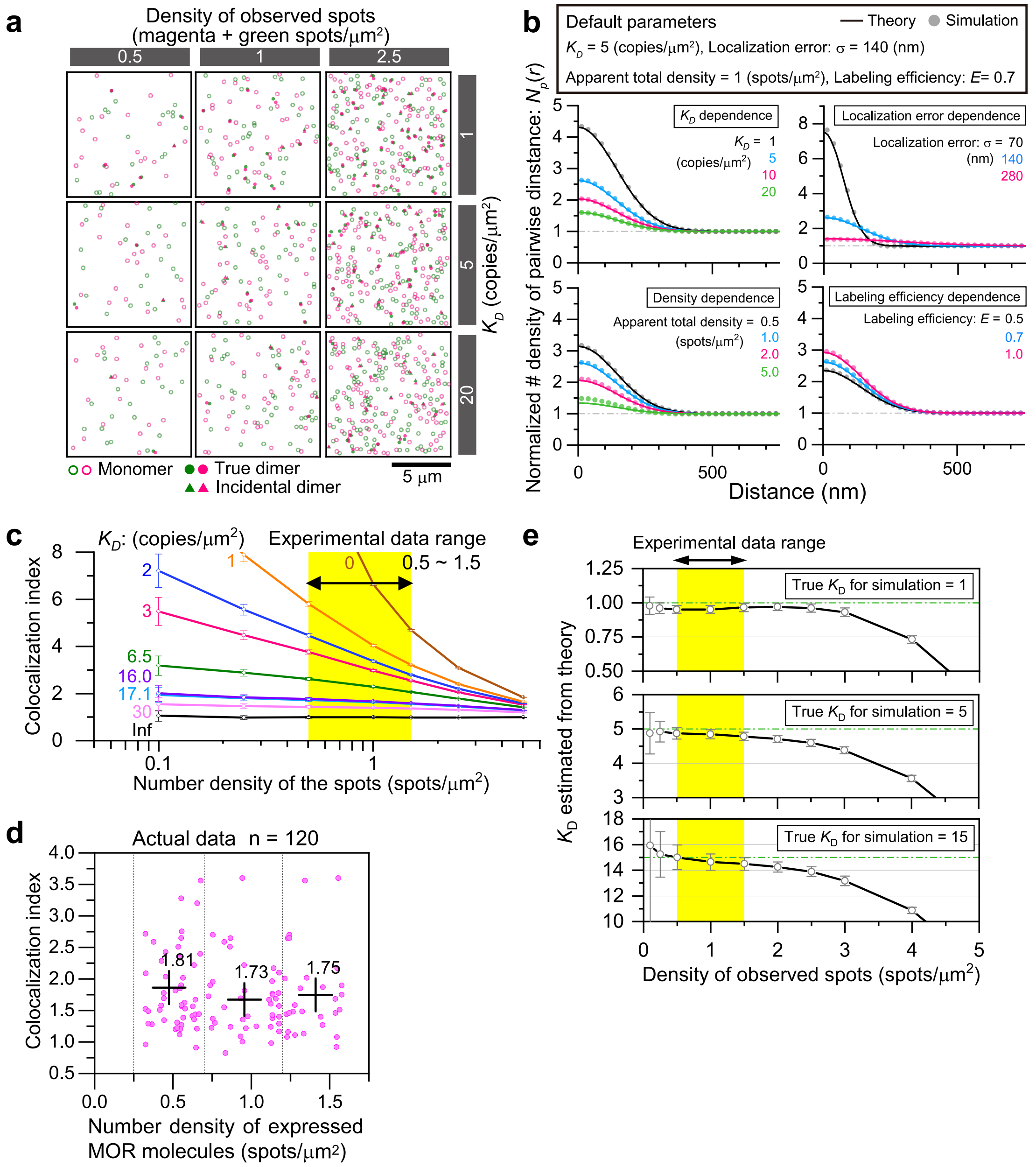


**Supplementary Fig. 3. The influences of the number density of the fluorescent spots in the image (expression levels of molecules) on PCCF, colocalization index, and $K_D$ obtained by using the theory developed in Supplementary Note 2, examined by Monte-Carlo simulations.**

**a.** Typical distributions of the fluorescent spots in the square ROI (10 μm x 10 μm) in the two-color images, generated by Monte-Carlo simulations. The numbers of magenta and green spots ($sN_{TA}$ and $sN_{TB}$ in Supplementary Note 2) are approximately the same, and the number densities of the spots shown are the total numbers of magenta and green spots. For the detailed method of the simulation, see the end of this caption. This figure represents a snapshot showing the spatial distributions of the fluorescent spots at the end of step 6 in the simulation steps, in the method part at the end of this caption.

**b.** Dependence of PCCF on four primary parameters, $K_D$, $\sigma$ (error for single-molecule localization and overlay of two-color images), total spot density (of green and magenta molecules), and labeling efficiency. The simulation-based PCCFs obtained from the simulated spot distributions (+; see step 8-1 in the method at the end of this caption; representing experimentally obtained PCCFs) are compared with theoretically obtained PCCFs (solid curves) as described in Supplementary Note 2, to find the applicability limitations of the theoretically obtained PCCF. This theory does not include the incidental colocalization of three or more molecules or true oligomers greater than dimers, but does include the incidental colocalization of two molecules as well as that of any two (but not three or more) combinations of true homodimers and true heterodimers. The theoretical PCCFs (solid curves) and $N_{AB}$ were calculated with the given $sN_{TA}$, $sN_{TB}$, and $K_D$ using Eqs. 1, 28-32, and 42-43 in Supplementary Note S2.

The theoretical PCCFs (solid curves) agree well with the simulation-based experimental PCCFs in broad ranges of the parameters (+), except for the case in which the spot density is 5 spots/μm$^2$ (**bottom-left**). Under these conditions, the incidental colocalizations of three or more molecules, which we neglected in our theory, would become important. However, spot densities ⩾5 spots/μm$^2$ are unrealistic for single-molecule imaging anyway (individual single molecules are not distinctly detectable).

**c.** Dependence of the colocalization index on the number density of spots in the image (top) and $K_D$ (bottom), found by Monte-Carlo simulations. The range of the total spot densities employed in the present research (0.5 - 1.5 spots/μm$^2$) is highlighted in yellow (top). In the $K_D$ ranges found for GPCRs; i.e., 3.6 copies/μm$^2$ for formyl-peptide receptor[7] and 1.6 copies/μm$^2$ for $\alpha_2$-adrenergic receptor,[9] as well as 5.9, 15.2, and 16.4 copies/μm$^2$ for KOR, MOR, and DOR, respectively (present research), the colocalization index weakly depends on the spot number density in the experimental range employed here (0.5 - 1.5 spots/μm$^2$). Also see **f**.

**d.** Experimental data showing the dependence of the colocalization index of MOR on the spot number density in the image (spots/μm$^2$). This panel is the only one in Figure S4 showing experimental results (other panels show Monte Carlo simulation results). Each magenta key represents the colocalization index and the spot number density for each examined cell (n = 120 cells). The plus marks placed at spot number densities of 0.475,

0.95, and 1.425 spots/μm$^2$ indicate the mean colocalization index values for the cells with spot number densities in the ranges of 0.25 - 0.7, 0.7 - 1.2, and 1.2 - 1.65 spots/μm$^2$, respectively. This result shows that the variations of the spot number density in the range of 0.5-1.5 spots/μm$^2$ would not be the limiting factor for the accuracy in determining the colocalization index of MOR.

**e.** Dependence of the estimated $K_D$ on the total spot number density, evaluated by simulation, using true $K_D$s of 1, 5, and 15 copies/μm$^2$ (with localization + overlay error of 140 nm and labeling efficiency of 0.7). The range of the total spot densities employed in the present research, 0.5 - 1.5 spots/μm$^2$, is highlighted in yellow. In this range, the estimated $K_D$ using the method described here is underestimated (higher affinities). The underestimation is worse with an increase in the spot number density, suggesting that this is caused by neglecting the incidental colocalization of more than two molecules in our theory.

**The Monte-Carlo simulation method for examining the dependence of the colocalization index, PCCF, and $K_D$ on the number density of the fluorescent spots in the images of molecules A and B.**

We describe the steps for simulating homodimerization, as detected by simultaneous two-color single-molecule imaging using two different fluorescent probes. Therefore, molecules A and B (the same protein but conjugated with two different dyes) form homodimers (AA + BB) and heterodimers (AB + BA) at equal densities. All equation numbers refer to those in Supplementary Note 2.

For the simulation, we give the total numbers of fluorescent spots (in the square 10 μm x 10 μm ROI). In **a**, $sN_{TA}$ = $sN_{TB}$ = [0.25, 0.5, and 1.25 copies/μm$^2$] x 100 μm$^2$ or the total numbers, $sN_{TA}$ + $sN_{TB}$, of [0.5, 1, and 2.5 copies/μm$^2$] x 100 μm$^2$. We also give the $K_D$ value for the simulation. In **a**, we used $K_D$ = 1, 5, and 20 copies/μm$^2$.

1. Using Eqs. 45-49 and 59-60, the true numbers of molecules $N_{0A}$ and $N_{0B}$ ($N_{0A}$ = $N_{0B}$) are determined, and using the given $K_D$ values, $N_{0AB}$ is calculated using Eq. 1. Note that, in the homodimerization assay, $N_{0AB} = N_{0AA} = N_{0BB}$.
2. The calculated numbers of molecules A and B ($N_{0A}$ and $N_{0B}$; equal numbers) and dimers ($N_{0AB}$, $N_{0AA}$, $N_{0BB}$; equal numbers) are randomly placed in a square 10 μm × 10 μm ROI, with additional shifts of positions to include the localization accuracy (Gaussian localization error $\sigma^2 = \sigma_A{}^2 + \sigma_B{}^2 + \sigma_{OL}{}^2$ ) (Eq. 14), which is (140 nm)$^2$ in **a**.

3. To include the effect of incomplete labeling (labeling efficiency, $E$), from all spots of A and B, $(1-E)$ x 100% of the spots are eliminated randomly.

4. To change the homodimers AA and BB to the apparent monomers A and B, respectively, in the image, one of the two spots in the remaining homodimer (after step 4) is eliminated.

5. To simulate incidental colocalization, a cluster analysis is performed. The clusters of the spots are identified by the hierarchical clustering algorithm (single-linkage), which links the nearest-neighbor spots within a threshold distance of 200 nm ($R_{\mathrm{Th}}$; determined from the diffraction limit). The linkage is performed by using Matlab's built-in function, "linkage" with 'single' method, and the clusters are identified by Matlab's built-in function, "cluster". Each identified cluster of spots is then replaced by a spot placed at the mass center of the cluster. The distributions of the spots shown in **a** are those after this step (step 6).

6. The process of steps 3 to 6 is repeated 200 times. We call the set of these 200 independent simulated images a movie, and produce 20 movies. For each movie, the normalized pairwise distance histogram (PCCF simulating the experimental PCCF) is obtained for each frame and then averaged over 200 frames. These averaged PCCFs are obtained for 20 movies and further averaged (the PCCF in the form of a histogram in which the value for each bin is the mean value obtained 4,000 times; i.e., 200 frames/movie x 20 movies). Using the averaged PCCF, $K_D$ is calculated using the theory described in Supplementary Note 2.

7. The process of steps 3 to 7 is repeated 100 times.

8-1) The averaged PCCF obtained at step 7 is averaged 100 times (the value for each bin is the mean of the 100 mean values), to obtain the average of the averaged PCCF, which we call the experimental PCCF obtained by simulation (histogram shown with + keys in **b**).

8-2) The graph in **c**, showing the colocalization index determined from the simulation, is plotted against the $K_D$ value (**bottom**), and the spot number density, [$sN_{TA}$ + $sN_{TB}$]/[ROI area size of 10 μm x 10 μm)] (**top**), given at step 1, is obtained using all of the simulated spot distributions.

8-3) Each spot in **e** shows the mean of the 100 $K_D$ values estimated by using the theory developed in Supplementary Note 2 for simulated distributions of the spots, starting from the given $K_D$ value (true $K_D$ value) and the total number of observed spots ($sN_{TA}$ + $sN_{TB}$), given at step 1. The theoretically estimated $K_D$ value for a given true $K_D$ is plotted as a function of the density of the total number of observed spots ($sN_{TA}$ + $sN_{TB}$)/[ROI area].

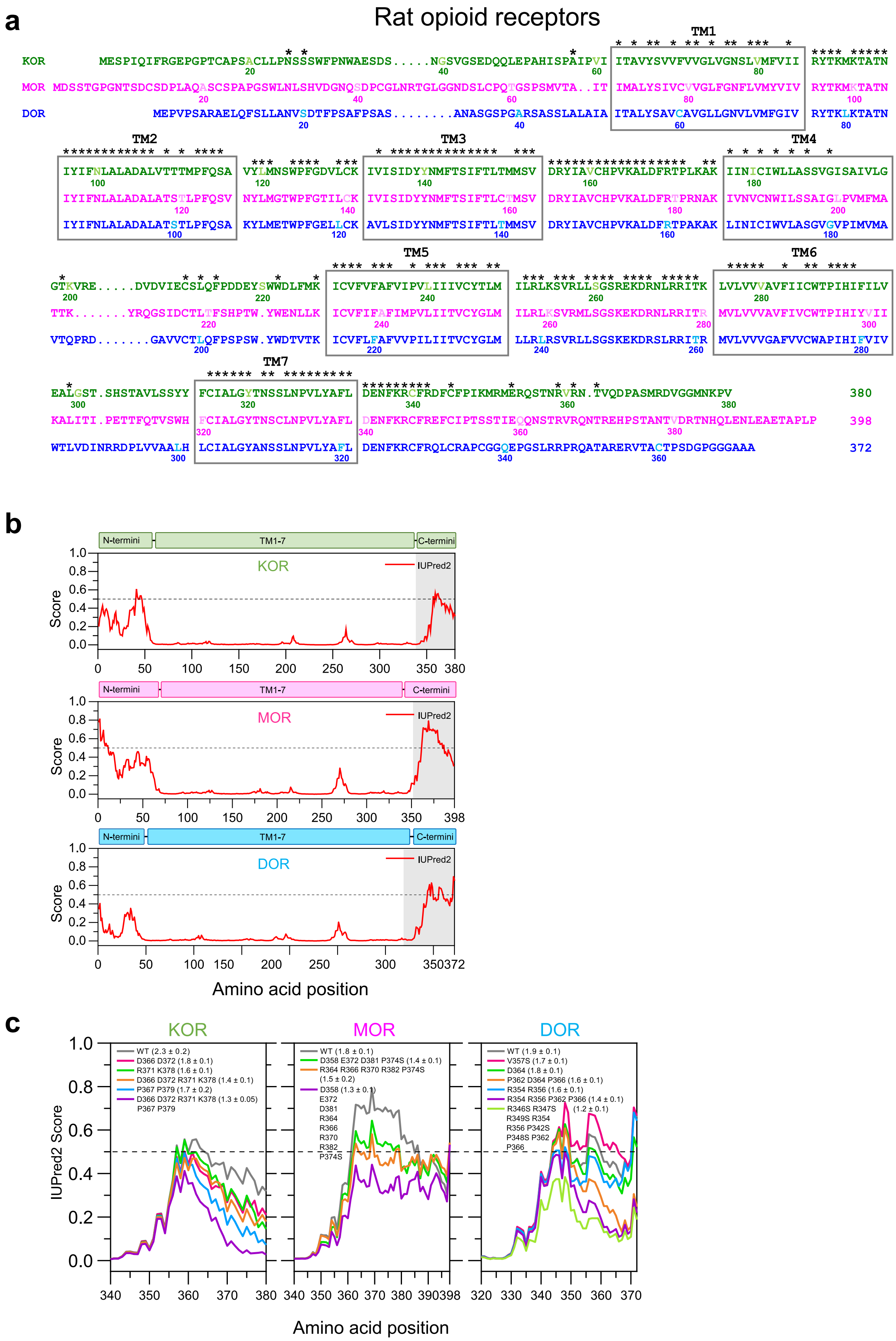


**Supplementary Fig. 4. Sequence comparison among the three classical ORs (rat), and IUPred2A disorder score plots suggesting the intrinsically disordered conformations of the C-terminal cytoplasmic domains of ORs.**

**a.** Sequence comparison among the three classical subtypes of rat ORs. The seven putative membrane-spanning domains are enclosed in boxes. Asterisks mark the identical amino acids among the three ORs.

**b.** The IUPred2A disorder score, a parameter indicating the level of intrinsic conformational disorder,[10,11] plotted against the full amino-acid sequence of each OR. The amino-acid sequences with IUPred2A disorder scores greater than 0.5 for some lengths are often considered intrinsically disordered. The C-terminal cytoplasmic domains exhibit propensities of being intrinsically disordered.

**c.** The IUPred2A disorder scores plotted against the sequences in the C-terminal cytoplasmic region of the WT- and mutant-ORs. For each curve, the amino acid mutations and colocalization index (in parentheses) are indicated. With an IUPred2A score reduction, the colocalization index generally becomes smaller, suggesting the involvement of disordered conformations in homodimerization.

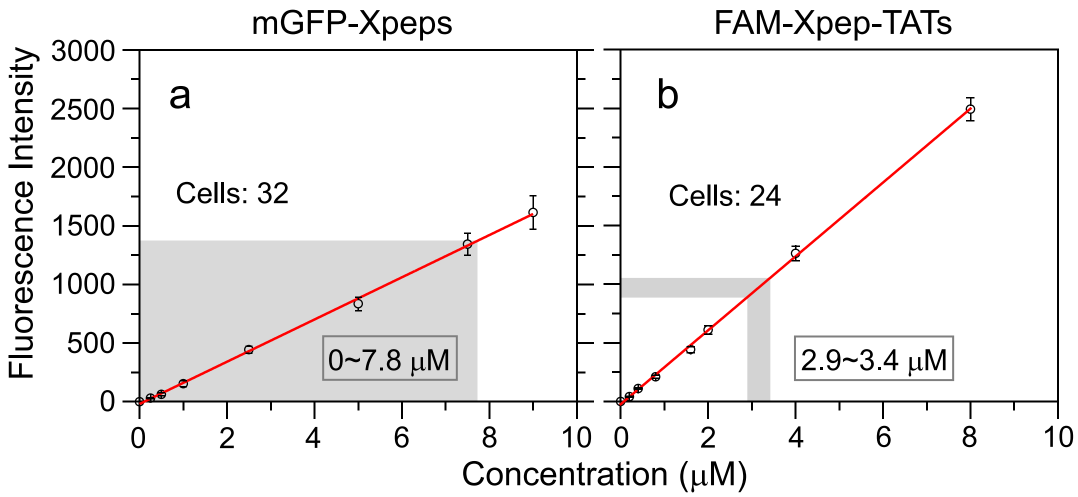


**Supplementary Fig. 5. Calibration curves (red straight lines) for evaluating the concentrations of mGFP-Xpeps and FAM-Xpep-TATs in the cytosol.**

Purified mGFP and FAM-Kpep-TAT were dissolved in Ham's F12 observation medium at various concentrations. The solutions were placed in glass-base dishes, and then confocal images were captured with a focus at 5 μm above the cover-glass surface. The confocal microscopy was performed on the same microscope station used for single-molecule imaging at 37°C, equipped with an Olympus SR10 spinning-disc confocal super-resolution unit (see Methods). The fluorescence intensities in a 10 μm x 10 μm square ROI were plotted as a function of the concentration of mGFP (**a**) or FAM-Kpep-TAT (**b**) (n = three dishes). The ranges of the fluorescence intensities and the peptide concentrations in the cells used for the actual observations are shown by grey areas.

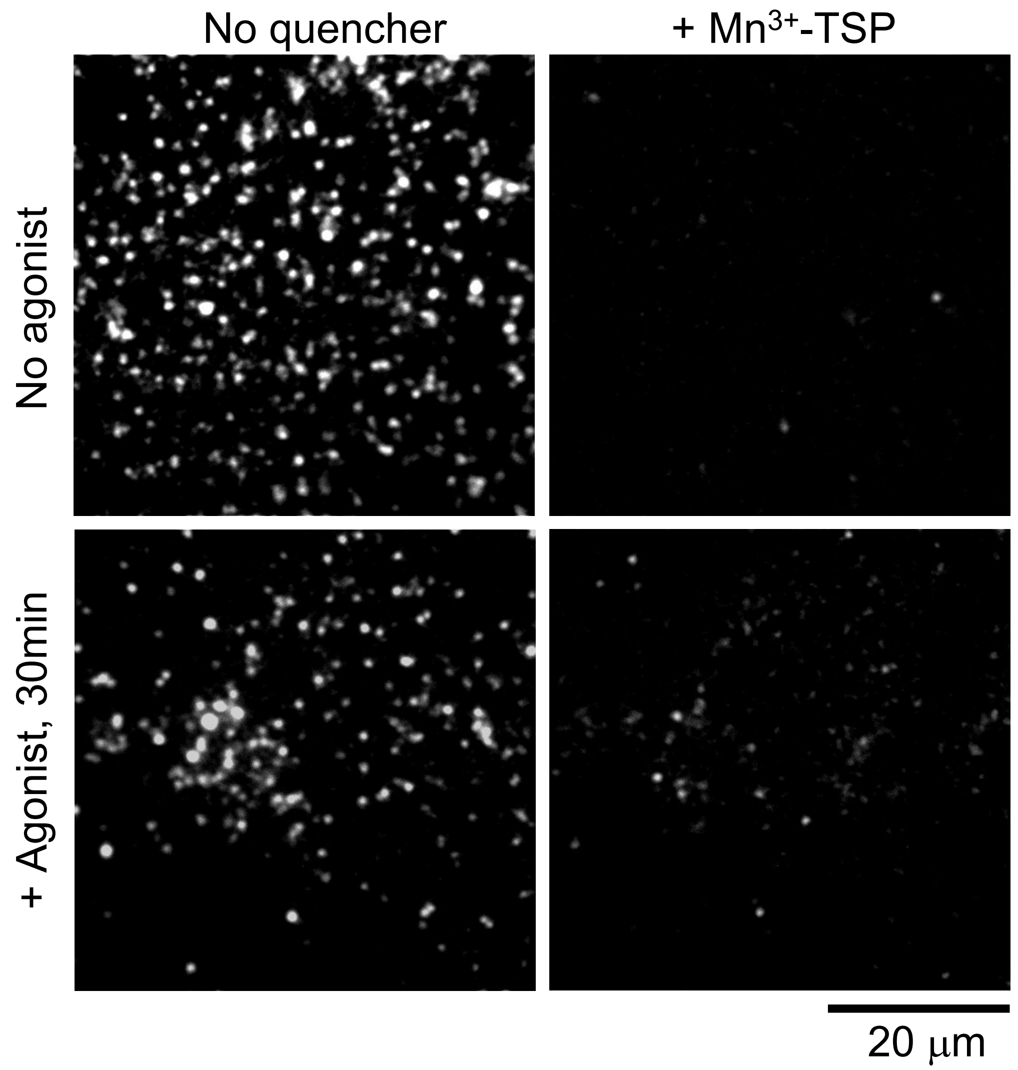


**Supplementary Fig. 6. Images of SNAPf-DOR labeled with SNAP-Surface 549 before and after the addition of the membrane-impermeable fluorescence quencher $Mn^{3+}$-TSP, in the absence of the agonist SNC-80 and 30 min after the agonist addition.**

The fluorescent spots after the addition of the quencher represent the internalized SNAPf-DOR molecules (labeled with SNAP-Surface 549).

**Supplementary Table 2. Predicted numbers of protomers expected to exist as monomers and dimers in the PM at different expression levels (total numbers of molecules in the PM).**

For the number densities of ORs as 0.3, 1, 3, and 10 copies/μm$^2$ (the same as in Fig. 2b), if the cell's surface area is 1500 μm$^2$ (for example, a disk-shaped cell with a 15.5 μm radius and no thickness, although this is unrealistic for neurons), the total numbers of expressed molecules of 450, 1,500, 4,500, and 15,000 copies per cell, respectively. This table shows how these numbers of protomers distribute between monomers and homodimers. Agonists used are U-50488 for KOR, DAMGO for MOR, and SNC-80 for DOR.

| ORs | Agonist | Predicted number of protomers existing as monomers and dimers in the PM (copies) at different expression levels | | | | | | | |
|---|---|---|---|---|---|---|---|---|---|
| | | 450 copies | | 1,500 copies | | 4,500 copies | | 15,000 copies | |
| | | Monomers | Dimers | Monomers | Dimers | Monomers | Dimers | Monomers | Dimers |
| KOR | − | 410 | 40 | 1,200 | 300 | 2,800 | 1,700 | 6,500 | 8,500 |
| | + | 430 | 20 | 1,400 | 100 | 3,500 | 1,000 | 8,800 | 6,200 |
| MOR | − | 430 | 20 | 1,300 | 200 | 3,500 | 1,000 | 8,700 | 6,300 |
| | + | 420 | 30 | 1,300 | 200 | 3,100 | 1,400 | 7,300 | 7,700 |
| DOR | − | 440 | 10 | 1,400 | 100 | 3,500 | 1000 | 8,900 | 6,100 |
| | + | 420 | 30 | 1,300 | 200 | 3,000 | 1,500 | 7,100 | 7,900 |

**Supplementary Table 3. Summary of the fractions of internalized ORs, detectable by the observation durations employed here, and their residency lifetimes in the PM. CHO-K1 cells vs. T24 cells. The agonists were 0.2 μM U-50488 for KOR, DAMGO for MOR, and SNC-80 for DOR (see Fig. 6b).**

| | | CHO-K1 | | T24 | |
|---|---|---|---|---|---|
| ORs | Additions | Fraction of internalized molecules (%) | Residency time of internalized molecules (min) | Fraction of internalized molecules (%) | Residency time of internalized molecules (min) |
| KOR | Control | 8.4 ± 4.1 | 32.1 ± 23.8 | 3.9 ± 2.9 | 44.6 ± 46.1 |
| | + FAM-pep-TAT | 6.9 ± 2.4$^{\dagger}$ | 26.2 ± 15.4$^{\dagger}$ | 5.3 ± 4.2$^{\dagger}$ | 37.4 ± 42.8$^{\dagger}$ |
| | + Agonist | 51.4 ± 2.7$^{*}$ | 11.9 ± 1.7$^{*}$ | 70.3 ± 2.0$^{*}$ | 16.9 ± 1.0$^{*}$ |
| | + Agonist<br>+ FAM-pep-TAT | 60.5 ± 2.5$^{*\dagger}$ | 9.6 ± 1.2$^{*}$ | 65.6 ± 2.5$^{*}$ | 11.1 ± 1.2$^{*}$ |
| | + Agonist<br>+ mGFP-pep | - | - | 70.9 ± 1.0$^{*}$ | 11.5 ± 0.4$^{*}$ |
| MOR | Control | 6.4 ± 1.9 | 25.8 ± 12.6 | 4.2 ± 2.4 | 40.3 ± 31.8 |
| | + FAM-pep-TAT | 6.3 ± 1.0$^{\dagger}$ | 22.1 ± 4.1$^{\dagger}$ | 5.2 ± 2.6$^{\dagger}$ | 36.8 ± 26.3$^{\dagger}$ |
| | + Agonist | 28.1 ± 1.7$^{*}$ | 11.3 ± 1.9$^{*}$ | 26.3 ± 1.1$^{*}$ | 6.5 ± 1.2$^{*}$ |
| | + Agonist<br>+ FAM-pep-TAT | 28.9 ± 2.0$^{*}$ | 10.7 ± 2.1$^{*}$ | 29.2 ± 1.9$^{*}$ | 14.4 ± 2.2$^{*}$ |
| | + Agonist<br>+ mGFP-pep | - | - | 28.6 ± 2.0$^{*}$ | 10.6 ± 2.2$^{*}$ |
| DOR | Control | 4.5 ± 3.5 | 29.8 ± 37.4 | 2.9 ± 1.0 | 24.2 ± 5.6 |
| | + FAM-pep-TAT | 4.9 ± 2.5$^{\dagger}$ | 26.2 ± 22.6$^{\dagger}$ | 3.9 ± 1.0$^{\dagger}$ | 21.4 ± 9.6$^{\dagger}$ |
| | + Agonist | 61.6 ± 1.7$^{*}$ | 10.8 ± 0.9$^{*}$ | 56.5 ± 1.9$^{*}$ | 8.9 ± 1.0$^{*}$ |
| | + Agonist<br>+ FAM-pep-TAT | 55.0 ± 1.0$^{*\dagger}$ | 10.1 ± 0.5$^{*}$ | 57.2 ± 2.9$^{*}$ | 10.5 ± 1.6$^{*}$ |
| | + Agonist<br>+ mGFP-pep | - | - | 56.5 ± 1.6$^{*}$ | 9.9 ± 0.9$^{*}$ |

* Significantly different from the values for the control samples. The significance was examined for all applicable cases, and when the difference was non-significant, no indication is shown. For the *p* values, see Supplementary Table 1.

† Significantly different from the values for the samples of "+ Agonist". The significance was examined for all applicable cases, and when the difference was non-significant, no indication is shown. For the *p* values, see Supplementary Table 1.

### Supplementary Note 1: Theory for evaluating the dissociation rate constant $k_{\text{off}}$ (and thus the homodimer lifetime $\tau_2$) from the single-molecule colocalization duration distribution

For the outline of the theory, see the main text.

#### (1) Dwell time in the absence of molecular binding

The scope of this **Note S1** is to provide the exact results for the time spent by a diffusing molecule in a circular domain. Our first step is to compute the distribution of dwell times in the absence of molecular binding. To this aim, we consider the two-dimensional diffusion equation

$$\partial_t p(x,y;t) = D\nabla^2 p(x,y;t) \quad . \tag{1}$$

We rewrite the equation in polar coordinates:

$$\partial_t p(r,\theta;t) = D\left(\partial_r{}^2 + \frac{1}{r}\partial_r + \frac{1}{r^2}\partial_\theta{}^2\right) p(r,\theta;t) \quad . \tag{2}$$

We consider isotropic solutions, so that $p(r,\theta;t)$ does not depend on the angle $\theta$ and the derivative with respect to the angle vanishes. We wish to solve the equation in the domain $r \in [0,R]$ with an absorbing boundary condition in $R$. The isotropic probability $p(r,t)$ is initially normalized

$$\int_0^R dr\, 2\pi r\, p(r;t=0) = 1 \quad . \tag{3}$$

We choose a localized initial condition, $p(r;t=0) = \delta(R-\epsilon-r)/[2\pi(R-\epsilon)]$. The parameter $\epsilon \ll R$ is a regularization: we cannot place the initial condition on the circle but only close to it, since this is an absorbing state. The solution to Eq. 2 satisfying this initial condition and the boundary condition is

$$p(r;t) = \sum_{n=1}^{\infty} \frac{1}{\pi R^2 J_1{}^2(\mu_n)} J_0\left(\mu_n \frac{r}{R}\right) J_0\left(\mu_n \frac{R-\epsilon}{R}\right) e^{-\frac{D\mu_n{}^2 t}{R^2}} \quad , \tag{4}$$

see e.g..[12] In this expression, $\mu_n$ is the $n$-th real positive zero of the Bessel function $J_0(x)$. The survival probability is then

$$S(t) = \int_0^R dr\, 2\pi r\, p(r;t)$$

$$= \sum_{n=1}^{\infty} \frac{2}{\mu_n{}^2 J_1{}^2(\mu_n)} [\mu_n J_1(\mu_n)] J_0\left(\mu_n \frac{R-\epsilon}{R}\right) e^{-\frac{D\mu_n{}^2 t}{R^2}} \tag{5}$$

$$(6)$$

$$= \sum_{n=1}^{\infty} \frac{2}{\mu_n J_1(\mu_n)} J_0\left(\mu_n \frac{R-\epsilon}{R}\right) e^{-\frac{D\mu_n{}^2 t}{R^2}} \quad ,$$

where we used $\int dx'x'J_0(x') = xJ_1(x) + C$. Note that $S(0) = 1$ due to the relation $1 = 2\sum_{m=1}^{\infty} J_0(y\mu_n)/[\mu_n J_1(\mu_n)]$, valid for any $y$ in the range $0 \le y < 1$; see Eq. 10.23.21 in NIST Digital Library of Mathematical Functions. https://dlmf.nist.gov.

The dwell time distribution $p_d(t)$ is given by the derivative of the survival probability with a changed sign:

$$p_d(t) = -\frac{d}{dt}S(t) = \sum_{n=1}^{\infty} \frac{2D\mu_n}{R^2 J_1(\mu_n)} J_0\left(\mu_n \frac{R-\epsilon}{R}\right) e^{-\frac{D{\mu_n}^2 t}{R^2}} \quad . \tag{7}$$

Finally, we use $\epsilon \ll R$ and expand:

$$J_0\left(\mu_n - \mu_n \frac{\epsilon}{R}\right) \approx J_0(\mu_n) - J_0'(\mu_n)\mu_n \frac{\epsilon}{R} = J_1(\mu_n)\mu_n \frac{\epsilon}{R} \quad . \tag{8}$$

This approximation is only valid for small $\mu_n \epsilon/R$. Substituting this result into Eq. 7, we obtain

$$p_d(t) \approx \sum_{n=1}^{\infty} \frac{2D\epsilon{\mu_n}^2}{R^3} e^{-\frac{D{\mu_n}^2 t}{R^2}} \quad . \tag{9}$$

This latter expression is handy as it does not involve Bessel functions, but only their zeros $\mu_n$. In particular, the first three zeros are $\mu_1 \approx 2.40$, $\mu_2 \approx 5.52$, $\mu_3 \approx 8.65$. As the argument of the exponential is proportional to $-{\mu_n}^2$, we truncate the series to only the first two terms, thus explaining the double-exponential behavior observed in the experiments.

#### (2) Dwell time with binding/unbinding events

We now consider the case in which a molecule, once inside the circle, can bind with other molecules, possibly multiple times. We denote the binding rate with $k_+$ and the unbinding rate with $k_-$. Calling $t$ the dwell time (excluding the time when the chosen molecule is bound), the number $n$ of binding events is a Poissonian variable with average $\lambda = tk_+$. The duration $\tau_i$ of each binding event $i$ is exponentially distributed with mean $k_-$. The total time $T$ spent inside the circle is equal to the sum of the time of unbound dwelling plus the sum of the individual times when the molecule is bound:

$$T = t + \sum_{i=1}^{n} \tau_i \quad . \tag{10}$$

We now compute the distribution $P(T)$ of the total time. Our starting point is the expression

$$P(T) = \int_0^{\infty} dt\, p_d(t) \left[\sum_{n=0}^{\infty} \frac{(k_+ t)^n e^{-k_+ t}}{n!}\right] \left(\prod_{i=1}^{n} \int_0^{\infty} d\tau_i\, k_- e^{-k_- \tau_i}\right) \cdot \delta\left(t + \sum_{i=1}^{n} \tau_i - T\right) \quad . \tag{11}$$

We substitute the integral representation of the delta function:

$$P(T) = \frac{1}{2\pi} \int_{-\infty}^{\infty} ds \int_0^{\infty} dt\, p_d(t) \left[\sum_{n=0}^{\infty} \frac{(k_+ t)^n e^{-k_+ t}}{n!}\right] \left(\prod_{i=1}^{n} \int_0^{\infty} d\tau_i\, k_- e^{-k_- \tau_i}\right) \cdot e^{is\left(t + \sum_{i=1}^{n} \tau_i - T\right)} . \tag{12}$$

We are now in the position to evaluate the integrals over $\tau_i s$:

$$P(T)=\frac{1}{2\pi}\int_{-\infty}^{\infty}ds\int_{0}^{\infty}dt\,p_d(t)\left[\sum_{n=0}^{\infty}\frac{(k_+t)^n e^{-k_+t}}{n!}\right][g_\tau(s)]^n e^{is(t-T)}\quad , \tag{13}$$

where we introduced the generating function $g_\tau(s)=\int_0^\infty d\tau\, k_- e^{-k_-\tau}e^{is\tau}=(1-is/k_-)^{-1}$. We compute the sum over $i$:

$$P(T)=\frac{1}{2\pi}\int_{-\infty}^{\infty}ds\int_{0}^{\infty}dt\,p_d(t)\,e^{k_+t[g_\tau(s)-1]}e^{is(t-T)}\quad . \tag{14}$$

To perform the integral over $t$, we approximate the distribution $p_d(t)$ using only the first mode: $p_d(t)\approx\alpha e^{-\alpha t}$ with $\alpha=D{\mu_1}^2/R^2$. This approximation is well justified by our experimental results. Computing the integral within this approximation, we obtain

$$P(T)=\frac{1}{2\pi}\int_{-\infty}^{\infty}ds\frac{\alpha}{\alpha+k_+[1-g_\tau(s)]-is}e^{-isT} \tag{15}$$

$$=\frac{1}{2\pi}\int_{-\infty}^{\infty}ds\frac{(1-is/k_-)}{(1-is/\alpha)(1-is/k_-)-isk_+/(k_-\alpha)}e^{-isT}\quad . \tag{16}$$

We rewrite Eq. 15 as

$$P(T)=\frac{1}{2\pi}\int_{-\infty}^{\infty}ds\frac{\alpha k_-(is/k_--1)}{(s-s_+)(s-s_-)}e^{-isT}\quad , \tag{17}$$

where the two roots $s_+, s_-$ are given by

$$s_\pm=\frac{i}{2}(-\Delta\pm\Gamma)\quad , \tag{18}$$

where $\Delta=(\alpha+k_-+k_+)$ and $\Gamma=\sqrt{\Delta^2-4\alpha k_-}$. Both roots are purely imaginary and with negative imaginary parts. We now compute the integral in Eq. 15 by closing the path in the complex plane with a semicircle over the negative imaginary half-plane. Thanks to Jordan's lemma, the integral over this semicircle vanishes. Our integral is therefore equal to the sum of the residues:

$$P(T)=(\alpha k_-)i\left[\frac{1-is_+/k_-}{s_+-s_-}e^{-is_+T}+\frac{1-is_-/k_-}{s_--s_+}e^{-is_-T}\right]$$

$$=\frac{\alpha k_-}{\Gamma}\left[(1-is_+/k_-)e^{-is_+T}-(1-is_-/k_-)e^{-is_-T}\right] \tag{19}$$

(20)

$$=\frac{\alpha k_-}{\Gamma}\left\{\left[1-\frac{(\Delta-\Gamma)}{2k_-}\right]e^{-\frac{T}{2}(\Delta-\Gamma)}\right. \tag{21}$$

$$\left.-\left[1-\frac{(\Delta+\Gamma)}{2k_-}\right]e^{-\frac{T}{2}(\Delta+\Gamma)}\right\}\quad .$$

In this case, the distribution is the difference of two exponentials. Proper normalization of the solution can be checked by direct integration and substituting the expressions of $\Delta$ and $\Gamma$.

#### (3) Limiting cases

**(3-1) Limit $k_+ \to 0$.**

In this limit, we should recover the result of the first section, since the binding probability is negligible. Taking the limit, we obtain $\Gamma \approx (\alpha - k_-)$ and therefore $(\Delta - \Gamma)/2 \approx \alpha$, $(\Delta + \Gamma)/2 \approx k_-$. Substituting into Eq. 19, we directly obtain $P(T) \approx \alpha e^{-\alpha T}$ as expected.

**(3-2) Limit $k_- \to 0$.**

In this limit, the rate of the first exponential in Eq. 19 diverges. The reason is that, if particles cannot unbind, then the permanence time inside the circle becomes infinite, as we are assuming that particles do not diffuse while bound.

#### (4) Dwell time with binding/unbinding events: a simpler derivation

In this section, we derive our main result following an alternative approach. We consider the probability $P_u(t)$, $P_b(t)$ that a molecule is still in the circle after a time $t$, while being unbound/bound (respectively) with other particles. These probabilities evolve according to the coupled rate equations:

$$\frac{d}{dt}P_u(t) = k_- P_b(t) - (k_+ + \alpha)P_u(t) \quad ,$$
$$\frac{d}{dt}P_b(t) = k_+ P_u(t) - k_- P_b(t) \quad . \qquad (22)$$

The eigenvalues associated with this linear system are

$$\lambda_\pm = \frac{1}{2}(-\Delta \pm \Gamma) \quad . \qquad (23)$$

Where $\Delta = (\alpha + k_- + k_+)$ and $\Gamma = \sqrt{\Delta^2 - 4\alpha k_-}$, as in the previous section. We need to solve the coupled rate equations 22 with the initial conditions $P_u(0) = 1$, $P_b(0) = 0$. The solution is then of the form

$$P_u(t) = \beta e^{\lambda_- t} + (1 - \beta) e^{\lambda_+ t} \quad ,$$
$$P_b(t) = \gamma \left(e^{\lambda_+ t} - e^{\lambda_- t}\right) \quad . \qquad (24)(25)$$

Substituting these expressions back into Eq. 22, we find

$$\beta = \frac{(\lambda_+ + k_+ + \alpha)}{(\lambda_+ - \lambda_-)} \quad , \qquad (26)$$

$$\gamma = -\frac{(\lambda_+ + k_+ + \alpha)}{(\lambda_+ - \lambda_-)} \frac{(\lambda_- + k_+ + \alpha)}{k_-} \quad . \qquad (27)$$

The probability of leaving the circle at time $T$ is equal to the derivative of the total probability of being bound and unbound with a changed sign:

$$P(T) = -\frac{d}{dt}\left[P_u(t) + P_b(t)\right]\Big|_{t=T} \qquad (28)(29)$$

$$= (\beta - 1 - \gamma)\lambda_+ e^{\lambda_+ T} + (\gamma - \beta)\lambda_- e^{\lambda_- T} \quad .$$

This distribution has the same double-exponential behavior of Eq. 19 with the same two decay rates. We numerically verified that Eqs. 19 and 28 are equivalent.

#### (5) Experimental evaluation of the unbinding rate constant $k_-$

In Eq. 19, we define the rate constants that can be determined by experiments:

$$k_1 = (\Delta + \Gamma)/2 \quad ,$$
$$k_2 = \frac{(\Delta - \Gamma)}{2} \tag{30)(31}$$

where $k_1$ and $k_2$ represent the faster and slower decay rates (inverse of the decay time constants) determined from experiments, respectively. From Eqs. 30 and 31, we obtain:

$$4k_1k_2 = \Delta^2 - \Gamma^2 \quad . \tag{32}$$

From the definition of $\Gamma$, we obtain

$$\Gamma^2 = \Delta^2 - 4\alpha k_- \quad . \tag{33}$$

Namely,

$$\Delta^2 - \Gamma^2 = 4\alpha k_- \quad . \tag{34}$$

By combining Eqs. 32 and 34, we obtain:

$$4k_1k_2 = 4\alpha k_- \quad . \tag{35}$$

Namely,

$$k_- = \frac{k_1k_2}{\alpha} \quad . \tag{36}$$

$k_1$ and $k_2$ can be directly determined from the experiment, and $\alpha$ (colocalization lifetime without actual binding) can be evaluated by overlaying an image obtained in one color (say cyan) with the image simultaneously obtained in the other color (say magenta) after a rotation of 180° (rotated overlay). Therefore, $k_-$ can be obtained from experiments. At the time resolution employed in the present study (33 ms), we found

$$k_1 \approx \alpha \quad . \tag{37}$$

Therefore, in the present study, with the limitation of the employed time resolution, we use the relationship

$$k_- \approx k_2 \quad . \tag{38}$$

### Supplementary Note 2

#### (1) Basic experimental and theoretical strategies

The purpose of this **Note 2** is to develop the theoretical framework to evaluate the dimer dissociation constant (dimer-monomer equilibrium constant) $K_{\mathrm{D}}$ for molecules in the monomer-dimer equilibrium state, using the simultaneous two-color single-molecule imaging method. In these experiments, the protein(s) of interest is (are) labeled with two fluorescent dyes with separable characteristic wavelengths, A and B, and the fluorescent spots are detected in the A- and B-channels. We will show that we can obtain $K_{\mathrm{D}}$ by experimentally determining the pair cross-correlation function (PCCF) of the fluorescent spots A and B; i.e., the distribution of the distances between all pairs of fluorescent spots A and B in the images, and counting the numbers of visible spots representing molecules A and B in the A- and B-channel images, respectively.

The subject matter of this study is the homodimerization of membrane molecules. However, even when homodimerization is examined, labeling the target molecule (one molecular species) with two different dye probes with distinct excitation-emission wavelengths (but maintaining the labeling condition of 1 dye molecule/1 protein copy) is often the preferred way of detecting homodimers, as done in this study. This is because the image analysis for detecting homodimers using simultaneously recorded two-color movies is easier than that using single-color movies (see the main text). In principle, the homodimer detection frequency using two-color movies is lower by a factor of two as compared with that using single-color movies, because the homodimers of the molecules with the same color are not detected in two-color experiments (since we neglect homodimers of molecules with the same color). However, we found that the ease of two-color colocalization detection outweighs the lower frequency of finding dimers.

When the dimerization of two molecular species A and B is examined, both homo- and hetero-dimerization must be considered (AA, BB, and AB, respectively). However, it is quite difficult to determine all three dimer dissociation equilibrium constants $K_{\mathrm{D}}$ at the same time. Thus, the two $\mathrm{homo}\text{–}K_{\mathrm{D}}$ must be determined in independent experiments before the $\mathrm{hetero}\text{–}K_{\mathrm{D}}$ determination. When the dimerization of only one molecular species (homodimerization) is examined using two probes (two-color movies), the molecules labeled with different dyes are regarded as different molecules. Namely, even for the investigation of homodimerization, the proteins labeled with different probes should be considered different molecules A and B, and the heterodimerization of proteins A and B (the same molecular species labeled with two different dyes) should be considered.

In two-color experiments, proteins A and B are imaged simultaneously in two separate optical channels A and B, respectively. The (x, y)-coordinates for individual fluorescent spots of A and B are determined in the separate images obtained in the optical channels A and B, and then are corrected for image distortions (with overlaying precision of $\sigma_{\mathrm{OL}}$, see Eq. 12). Using the corrected (x, y)-coordinates for all fluorescent spots representing molecules A and B, the distances of all pairs of detected spots A and B are calculated, providing the distribution, which is the PCCF. In the plasma membrane (PM), both the heterodimers and monomers are considered to be spread uniformly, and in this case the PCCF will be the sum of that for heterodimers and that for monomers, although in the experimental A- and B-channel images, we cannot directly determine which fluorescent spots represent heterodimers and monomers.

The dimer dissociation equilibrium constant (dimer-monomer equilibrium constant) $K_{\mathrm{D}}$ can be expressed by using the numbers of true monomers A and B ($N_{0\mathrm{A}}$ and $N_{0\mathrm{B}}$, respectively) and the number of true heterodimers of AB ($N_{0\mathrm{AB}}$), or the rate-constants of the forward and reverse reactions ($k_{\mathrm{on}}$ and $k_{\mathrm{off}}$, respectively),

$$K_{\mathrm{D}} = \frac{N_{0\mathrm{A}} N_{0\mathrm{B}}}{N_{0\mathrm{AB}} A} = \frac{k_{\mathrm{on}}}{k_{\mathrm{off}}} \quad , \tag{1}$$

where $A$ is the area of the ROI.

Accordingly, to obtain the $K_{\mathrm{D}}$, we will develop a method to evaluate $N_{0\mathrm{A}}$, $N_{0\mathrm{B}}$, and $N_{0\mathrm{AB}}$ from the numbers of "visible" spots A and B in the experimental images and the experimentally obtained PCCF. These spot numbers and the PCCF are obtained based on the visible fluorescent spots in the images detected in the A- and B-channels, and thus experimental limitations distort the number of fluorescent spots visible in the images and the experimentally obtained PCCF. Therefore, here, we will develop mathematical expressions for describing the true numbers $N_{0\mathrm{A}}$, $N_{0\mathrm{B}}$, and $N_{0\mathrm{AB}}$ using the numbers of visible fluorescent spots and the experimentally obtained PCCF, by including the effects of various experimental limitations.

#### (2) Mathematical expression of the pair cross-correlation function (PCCF) of molecules that form heterodimers in equilibrium with monomers in the plasma membrane (PM): an ideal case where all molecules are visible and separable in the image

Consider an ideal case in which all of the molecules in the ROI in the PM are visible (separable) in the image, and let us obtain the PCCF in this case. First, we will derive the PCCF for true AB heterodimers; i.e., the distribution of the distances between molecules A and B when they exist as true heterodimers. The true distance between molecules A and B in a true heterodimer is assumed to be 0. However, the experimentally measured distance will not be 0, due to errors in measurements of the particle positions and the overlaying accuracy of the two channels, and thus if we experimentally measure the distance between A and B in each pair, it will have certain distributions as a function of the localization error.

As described, the heterodimers and monomers coexist in the plasma membrane (PM), and they are both considered to be spread uniformly. The PCCF will be the sum of that for heterodimers and that for monomers, although there is no way to tell which fluorescent spots represent heterodimers and monomers in the experimentally obtained images in channels A and B. Therefore, secondly, we will obtain the PCCF for randomly distributed monomers A and B in the PM.

##### (2-1) PCCF for heterodimers: the distribution of the distances between molecules A and B when they exist as true heterodimers

The experimentally measured positions of molecules A and B could be described as probability variables $\mathrm{R_A} = (\mathrm{X_A^{err}}, \mathrm{Y_A^{err}})$ and $\mathrm{R_B} = (\mathrm{X_B^{err}}, \mathrm{Y_B^{err}})$, respectively, where $\mathrm{X_A^{err}}$, $\mathrm{Y_A^{err}}$, $\mathrm{X_B^{err}}$, and $\mathrm{Y_B^{err}}$ are independently distributed Gaussian random variables with zero means and variances ${\sigma_{\mathrm{Ax}}}^2$, ${\sigma_{\mathrm{Ay}}}^2$, ${\sigma_{\mathrm{Bx}}}^2$, and ${\sigma_{\mathrm{By}}}^2$, respectively. Then, the square of the distance between molecules A and B is given by

$$\mathrm{R}^2 = (\mathrm{X_A^{err}} - \mathrm{X_B^{err}})^2 + (\mathrm{Y_A^{err}} - \mathrm{Y_B^{err}})^2 \quad . \tag{2}$$

Here, we define $\mathrm{E_X} = \mathrm{X_A^{err}} - \mathrm{X_B^{err}}$ and $\mathrm{E_Y} = \mathrm{Y_A^{err}} - \mathrm{Y_B^{err}}$. $\mathrm{E_X}$ and $\mathrm{E_Y}$ are Gaussians with zero-means, and variances equal to ${\sigma_{\mathrm{Ax}}}^2 + {\sigma_{\mathrm{Bx}}}^2$ and ${\sigma_{\mathrm{Ay}}}^2 + {\sigma_{\mathrm{By}}}^2$, respectively, as the variance of the sum of two independent Gaussian random variables is the sum of the variances of each variable. Accordingly, writing the Gaussian function with the 0 mean and variance $s^2$ as $N(0,\ s^2)$,

$$\mathrm{E_X} = N(0,\ {\sigma_{\mathrm{Ax}}}^2 + {\sigma_{\mathrm{Bx}}}^2),\ \ \mathrm{E_Y} = N\left(0, {\sigma_{\mathrm{Ay}}}^2 + {\sigma_{\mathrm{By}}}^2\right)\ . \tag{3}$$

In many cases, the errors in the position measurements in the x and y directions are similar. Therefore, we assume

$$\sigma_{\mathrm{Ax}} = \sigma_{\mathrm{Ay}} = \sigma_{\mathrm{A}},\ \ \sigma_{\mathrm{Bx}} = \sigma_{\mathrm{By}} = \sigma_{\mathrm{B}}\ . \tag{4}$$

Re-writing **Eq. 2**, using **Eq. 3,**

$$\begin{aligned} \mathrm{R}^2 &= {\mathrm{E_X}}^2 + {\mathrm{E_Y}}^2\ , \\ &= ({\sigma_{\mathrm{A}}}^2 + {\sigma_{\mathrm{B}}}^2)\left[\left(\frac{\mathrm{E_X}}{\sqrt{{\sigma_{\mathrm{A}}}^2 + {\sigma_{\mathrm{B}}}^2}}\right)^2 + \left(\frac{\mathrm{E_Y}}{\sqrt{{\sigma_{\mathrm{A}}}^2 + {\sigma_{\mathrm{B}}}^2}}\right)^2\right]\ , \\ &= ({\sigma_{\mathrm{A}}}^2 + {\sigma_{\mathrm{B}}}^2)({\mathrm{E_X}'}^2 + {\mathrm{E_Y}'}^2)\ , \end{aligned} \tag{5}$$

where $\mathrm{E'_X} = N(0,1)$ and $\mathrm{E'_Y} = N(0,1)$ are Gaussians with zero-mean and unit variance. Then, ${\mathrm{E_X}'}^2 + {\mathrm{E_Y}'}^2$ has a chi-square distribution with 2-degrees of freedom.

$$\mathrm{R}^2 = ({\sigma_{\mathrm{A}}}^2 + {\sigma_{\mathrm{B}}}^2)\chi^2(2)\ . \tag{6}$$

The probability density of $\chi^2(2)$ is equal to $e^{-\frac{x}{2}}/2$, and thus the probability density function of $\mathrm{R}^2$ is given by

$$f_{\mathrm{R}^2}(x) = \frac{1}{2({\sigma_{\mathrm{A}}}^2 + {\sigma_{\mathrm{B}}}^2)} e^{-\frac{x}{2({\sigma_{\mathrm{A}}}^2 + {\sigma_{\mathrm{B}}}^2)}}\ , \tag{7}$$

since

$$f_{\mathrm{Y}}(y) = \frac{1}{a} f_{\mathrm{X}}(y/a),\ \ \text{for } \mathrm{Y} = a\mathrm{X}\ .$$

The cumulative density function of $\mathrm{R}^2$ is

$$F_{\mathrm{R}^2}(x) = 1 - e^{-\frac{x}{2({\sigma_{\mathrm{A}}}^2 + {\sigma_{\mathrm{B}}}^2)}}\ . \tag{9}$$

The cumulative density function of R can then be obtained from

$$F_{\mathrm{R}}(x) = \mathrm{Prob}(\mathrm{R} < x) = \mathrm{Prob}\left(\sqrt{\mathrm{R}^2} < x\right) = \mathrm{Prob}(\mathrm{R}^2 < x^2) = F_{\mathrm{R}^2}(x^2)\ . \tag{10}$$

Namely, the probability density function of R is calculated as,

$$f_{\mathrm{R}}(x) = \frac{d}{dx} F_{\mathrm{R}}(x) = \frac{d}{dx} F_{\mathrm{R}^2}(x^2) = \frac{x}{({\sigma_{\mathrm{A}}}^2 + {\sigma_{\mathrm{B}}}^2)} \exp\left(-\frac{x^2}{2({\sigma_{\mathrm{A}}}^2 + {\sigma_{\mathrm{B}}}^2)}\right)\ . \tag{11}$$

This represents the probability density function of the distances between molecules A and B in a true AB heterodimer. Meanwhile, its cumulative function would also be useful.

$$F_{\mathrm{R}}(x) = 1 - e^{-\frac{x}{2({\sigma_{\mathrm{A}}}^2 + {\sigma_{\mathrm{B}}}^2)}}\ . \tag{12}$$

We will need to include the overlaying precision of $\sigma_{\mathrm{OL}}$ for overlaying the two images separately (although simultaneously) obtained for molecules A and B (i.e., green and red signals).[13] By following the same argument made to determine **Eq. 11**, we obtain,

$$f_{\mathrm{R}}(x) = \frac{x}{\sigma^2} \exp\left(-\frac{x^2}{2\sigma^2}\right) \quad , \tag{13}$$

where

$$\sigma^2 = \sigma_{\mathrm{A}}{}^2 + \sigma_{\mathrm{B}}{}^2 + \sigma_{\mathrm{OL}}{}^2 \, . \tag{14}$$

For clarity, in the following, we write $f_{\mathrm{R}}(x)$ as $P_{\mathrm{HetD}}(r)$ (the probability density function describing the distribution of the distances between molecules A and B that exist as a true heterodimer AB); i.e.,

$$P_{\mathrm{HetD}}(r) = \frac{r}{\sigma^2} \exp\left(-\frac{r^2}{2\sigma^2}\right) \quad . \tag{15}$$

To convert the probability density function to the histogram of molecular number density (discrete function, $H_{\mathrm{HetD}}(r_i)$), we multiply the total number of heterodimers AB ($N_{\mathrm{AB}}$) existing in the ROI in the image and the bin width ($\Delta r$),

$$H_{\mathrm{HetD}}(r_i) = N_{\mathrm{AB}} \cdot \Delta r \cdot P_{\mathrm{HetD}}(r) = N_{\mathrm{AB}} \cdot \Delta r \cdot \frac{r_i}{\sigma^2} \exp\left(-\frac{r_i{}^2}{2\sigma^2}\right) \quad , \tag{16}$$

where $r_i$ is the central value of the $i$th bin.

**(2-2) PCCF for randomly distributed monomers A and B: the distribution of the distances between randomly distributed monomers A and B**

When monomers A and B are randomly distributed in the PM, the distribution of the distances between molecules A and B will be circularly symmetric. Therefore, by referring to the numbers of monomers A and B as $N_{\mathrm{A}}$ and $N_{\mathrm{B}}$, respectively, the distribution of the number of distances between randomly distributed monomers A and B (histogram, $H_{\mathrm{Random}}(r_i)$) can be expressed as

$$H_{\mathrm{Random}}(r_i) = N_{\mathrm{A}} N_{\mathrm{B}} \cdot \Delta r \cdot \frac{2\pi r_i}{A} \quad , \tag{17}$$

where $A$ is the ROI size.

**(2-3) PCCF for the case where randomly distributed heterodimers AB and monomers A and B coexist: the distribution of the number density of pairwise distances A and B**

We first define the total numbers of molecules A and B; i.e., the sum of the numbers of "monomers" A (B) and the numbers of molecule A (B) in the heterodimer AB ($N_{\mathrm{TA}}$, $N_{\mathrm{TB}}$, respectively). Namely,

$$N_{\mathrm{TA}} = N_{\mathrm{A}} + N_{\mathrm{AB}} \quad , \tag{18}$$

$$N_{\mathrm{TB}} = N_{\mathrm{B}} + N_{\mathrm{AB}} \quad . \tag{19}$$

Using $N_{\mathrm{TA}}$ and $N_{\mathrm{TB}}$, the total number of distances for randomly distributed molecules of A and B can be written as

$$N_{\mathrm{TA}} N_{\mathrm{TB}} - N_{\mathrm{AB}} \quad . \tag{20}$$

Accordingly, in the case where monomers A and B and heterodimers AB coexist, $N_{\mathrm{A}}N_{\mathrm{B}}$ in **Eq. 17** must be replaced by **Eq. 20**, and we obtain

$$H_{\mathrm{Random}}(r_i) = (N_{\mathrm{TA}}N_{\mathrm{TB}} - N_{\mathrm{AB}}) \cdot \Delta r \cdot \frac{2\pi r_i}{A} \quad . \tag{21}$$

When $N_{\mathrm{AB}}$ is 0, $N_{\mathrm{TA}} = N_{\mathrm{A}}$ and $N_{\mathrm{TB}} = N_{\mathrm{B}}$, and therefore, **Eq. 17** can be obtained from **Eq. 21**, indicating that **Eq. 17** is a special case of **Eq. 21**.

The histogram for the distribution of all pairwise distances between molecules A and B will be the sum of that in heterodimers and that for randomly distributed molecules A and B; i.e., the sum of **Eqs. 16 and 21**.

$$H_{\mathrm{T}}(r_i) = N_{\mathrm{AB}} \cdot \Delta r \cdot \frac{r_i}{\sigma^2} \exp\left(-\frac{{r_i}^2}{2\sigma^2}\right) + \frac{N_{\mathrm{TA}}N_{\mathrm{TB}} - N_{\mathrm{AB}}}{A} \cdot \Delta r \cdot 2\pi r_i \quad . \tag{22}$$

The histogram is more useful if it is normalized by the total number of distances at $r_i$, which is,

$$\frac{N_{\mathrm{TA}}N_{\mathrm{TB}}}{A} \cdot \Delta r \cdot 2\pi r_i \quad . \tag{23}$$

Therefore, the normalized number density of pairwise distances is expressed as

$$H_{\mathrm{Norm}}(r_i) = \frac{A \cdot N_{\mathrm{AB}}}{N_{\mathrm{TA}}N_{\mathrm{TB}} \cdot 2\pi\sigma^2} \exp\left(-\frac{{r_i}^2}{2\sigma^2}\right) + \left(1 - \frac{N_{\mathrm{AB}}}{N_{\mathrm{TA}}N_{\mathrm{TB}}}\right) \quad . \tag{24}$$

Under normal experimental conditions for single-molecule imaging, because $N_{\mathrm{TA}}N_{\mathrm{TB}}$ is far greater than $N_{\mathrm{AB}}$, the second term will become ~1.

This functional form is quite interesting. It shows that the width of the Gaussian function, and thus how rapidly the PCCF decreases from the 0 distance, is simply determined by the sum of the single-molecule localization accuracies for molecules A and B and the overlaying accuracy of the images A and B.

#### (3) Obtaining $N_{0\mathrm{A}}$, $N_{0\mathrm{B}}$, and $N_{0\mathrm{AB}}$ from the number of the fluorescent spots in the images in A- and B-channels and the experimentally determined PCCF

In the actual experiments, we only detect fluorescent spots representing A molecules in the A-channel image and those representing B molecules in the B-channel image, obtained simultaneously. We will call the numbers of these spots in the ROI $sN_{\mathrm{TA}}$ and $sN_{\mathrm{TB}}$, respectively. In this notation, "*s*" before "*N*" indicates that it is the number of "spots" in the image and not the number of "molecules"; meanwhile, T in the subscript is to clarify that we count all of the visible spots; i.e., "Total numbers".

Based on **Eq. 1**, to evaluate $K_{\mathrm{D}}$, we need to obtain $N_{0\mathrm{A}}$, $N_{0\mathrm{B}}$, and $N_{0\mathrm{AB}}$ from the experimentally determined PCCF and the numbers of fluorescent spots visible in the A- and B-channels, $sN_{\mathrm{TA}}$ and $sN_{\mathrm{TB}}$. These spot numbers are different from $N_{0\mathrm{A}}$ and $N_{0\mathrm{B}}$, respectively, due to the following three problems.

(Problem a) The possible presence of AA and BB homodimers,

(Problem b) Limited labeling efficiencies for molecules A and B, and

(Problem c) Incidental colocalizations of the fluorescent spots (due to the limited single-molecule localization precisions and overlaying precisions of images A and B).

Note that the limited single-molecule localization precisions and overlaying precisions of images A and B also make it impossible to directly determine, from images A and B, which A spots and B spots represent heterodimers (see the introductory part of Section 2, right before subsection 2-1). Namely, we are unable to directly count the number of heterodimers in the image. Therefore, we will develop a method to evaluate the number of heterodimers from the (simultaneously obtained) images A and B, which we will call $N_{\mathrm{AB}}$. Note that $N_{\mathrm{AB}}$ is different from $N_{0\mathrm{AB}}$ in that, while $N_{0\mathrm{AB}}$ is the correct number of heterodimers, $N_{\mathrm{AB}}$ is the number of heterodimers that can be determined from PCCF, meaning that it is the number of heterodimers in which both molecule A and B are fluorescently labeled. Due to the limited labeling efficiency, certain fractions of heterodimers consist of a labeled molecule A (B) and a non-labeled molecule B (A) and of non-labeled molecules A and B. The experimentally obtained PCCF only includes the heterodimers in which both molecules A and B are labeled. According to the definitions of $sN_{\mathrm{TA}}$ and $sN_{\mathrm{TB}}$, $sN_{\mathrm{TA}}$ and $sN_{\mathrm{TB}}$ include the number of spots representing $N_{\mathrm{AB}}$.

For the evaluation of an unknown $N_{\mathrm{AB}}$, we will take advantage of the experimentally determined PCCF, which is considered to be represented by **Eq. 24**, in which $N_{\mathrm{TA}}$ and $N_{\mathrm{TB}}$ are replaced by $sN_{\mathrm{TA}}$ and $sN_{\mathrm{TB}}$, respectively. This is because **Eq. 24** is the simple addition of the number of distances between A and B in heterodimers and those between randomly distributed “visible molecules” A and B. Therefore, **Eq. 24** is rewritten as

$$H_{\mathrm{Norm}}(r_i) = \frac{A \cdot N_{\mathrm{AB}}}{sN_{\mathrm{TA}} sN_{\mathrm{TB}} \cdot 2\pi\sigma^2} \exp\left(-\frac{{r_i}^2}{2\sigma^2}\right) + \left(1 - \frac{N_{\mathrm{AB}}}{sN_{\mathrm{TA}} sN_{\mathrm{TB}}}\right) \quad . \tag{25}$$

As described, the second term can be well approximated by 1. To determine the first term from the experiments, we fit the experimental histogram (PCCF) using the following Gaussian function

$$y = C \cdot \exp\left(-\frac{x^2}{2w^2}\right) + y_0 \quad , \tag{26}$$

where

$$C = \frac{A \cdot N_{\mathrm{AB}}}{sN_{\mathrm{TA}} sN_{\mathrm{TB}} \cdot 2\pi w^2} \quad , \tag{27}$$

from the comparison with **Eq. 25**. Namely,

$$N_{\mathrm{AB}} = \frac{C \cdot sN_{\mathrm{TA}} sN_{\mathrm{TB}} \cdot 2\pi w^2}{A} \quad . \tag{28}$$

$sN_{\mathrm{TA}}$ and $sN_{\mathrm{TB}}$ can be directly obtained by counting the fluorescent spots in the images in the A- and B-channels, and **Eq. 28** indicates that, by using these numbers, $sN_{\mathrm{TA}}$ and $sN_{\mathrm{TB}}$, as well as $C$ and $w^2$ determined from the experimentally obtained PCCF, $N_{\mathrm{AB}}$ can be determined. Therefore, our task here is simplified to determine the relationship of the set of true numbers $N_{0\mathrm{A}}$, $N_{0\mathrm{B}}$, and $N_{0\mathrm{AB}}$ with the experimentally obtainable numbers, $sN_{\mathrm{TA}}$, $sN_{\mathrm{TB}}$, and $N_{\mathrm{AB}}$. In this section 3, we will obtain this relationship.

In subsection (3-1), we will consider the effects of the presence of AA and BB homodimers (Problem a) and the limited labeling efficiencies for molecules A and B (Problem b) on the number of observable fluorescent spots in images A and B. For simplicity, we assume that we cannot differentiate the same two molecules when they are located within a threshold distance $R_{\mathrm{Th}}$, which is approximately the diffraction limit of the microscope resolution (~200 nm).

In subsection (3-2), we will derive the equations describing the effect of incidental colocalizations of the fluorescent spots (Problem c) on the number of observable fluorescent spots in the images A and B, $sN_{\mathrm{TA}}$ and $sN_{\mathrm{TB}}$, including the effects we consider in subsection (3-1). This will provide the relationship between the set of experimentally obtained numbers $sN_{\mathrm{TA}}$, $sN_{\mathrm{TB}}$, and $N_{\mathrm{AB}}$ and the set of three true numbers, $N_{0\mathrm{A}}$, $N_{0\mathrm{B}}$, and $N_{0\mathrm{AB}}$.

**(3-1) Effects of the presence of homodimers and limited labeling efficiencies**

Even when we are interested in only one molecule (one molecular species) and studying its homodimerization, we often examine heterodimerization by labeling the target molecular species with two different dyes, as described in subsection 1 (and done in this study). This means that in the experimentally obtained images, AA and BB homodimers will exist. Even in the case where different target molecules A and B are observed to examine AB heterodimers, molecules A and/or B could form homodimers. Therefore, to generalize the argument, we consider the case in which, in addition to AB heterodimers, AA and BB homodimers also form.

We consider the experiments in which we only detect the locations of fluorescent spots, but not their intensities. Even the intensity distribution of monomers is quite large, and so if possible, we would avoid depending on the spot signal intensities for detecting dimers (this is one of the reasons why we often prefer two-color experiments over single-color experiments). Due to this limitation, we do not distinguish monomers A (B) and homodimers AA (BB) in the images obtained in the A (B) channel.

Considering the case where both homo- and hetero-dimers co-exist as follows,

$$[\mathrm{A}] + [\mathrm{A}] \Leftrightarrow [\mathrm{AA}] \ , \tag{29}$$

$$[\mathrm{B}] + [\mathrm{B}] \Leftrightarrow [\mathrm{BB}] \ , \tag{30}$$

$$[\mathrm{A}] + [\mathrm{B}] \Leftrightarrow [\mathrm{AB}] \ . \tag{31}$$

In this system, simultaneous ordinary differential equations can be written as,

$$\frac{d[\mathrm{A}]}{dt} = -2k_{\mathrm{on1}}[\mathrm{A}]^2 + 2k_{\mathrm{off1}}[\mathrm{AA}] - k_{\mathrm{on3}}[\mathrm{A}][\mathrm{B}] + k_{\mathrm{off3}}[\mathrm{AB}] \ , \tag{32}$$

$$\frac{d[\mathrm{B}]}{dt} = -2k_{\mathrm{on2}}[\mathrm{B}]^2 + 2k_{\mathrm{off2}}[\mathrm{BB}] - k_{\mathrm{on3}}[\mathrm{A}][\mathrm{B}] + k_{\mathrm{off3}}[\mathrm{AB}] \ , \tag{33}$$

$$\frac{d[\mathrm{AA}]}{dt} = k_{\mathrm{on1}}[\mathrm{A}]^2 - k_{\mathrm{off1}}[\mathrm{AA}] \ , \tag{34}$$

$$\frac{d[\mathrm{BB}]}{dt} = k_{\mathrm{on2}}[\mathrm{B}]^2 - k_{\mathrm{off2}}[\mathrm{BB}] \ , \tag{35}$$

$$\frac{d[\mathrm{AB}]}{dt} = k_{\mathrm{on3}}[\mathrm{A}][\mathrm{B}] - k_{\mathrm{off3}}[\mathrm{AB}] \ , \tag{36}$$

where $k_{\mathrm{on1,2,3}}$ and $k_{\mathrm{off1,2,3}}$ represent the on-rate and off-rate constants for the dimerization of $[\mathrm{AA}]$, $[\mathrm{BB}]$, and $[\mathrm{AB}]$, respectively.

Assuming that the total densities of molecule A and B ($[\mathrm{TA}]$ and $[\mathrm{TB}]$) are not changed,

$$[\mathrm{TA}] = [\mathrm{A}] + 2[\mathrm{AA}] + [\mathrm{AB}] \tag{37}$$

$$[\mathrm{TB}] = [\mathrm{B}] + 2[\mathrm{BB}] + [\mathrm{AB}]. \tag{38}$$

Using these relationships, **Eqs. 34-36** can be re-written as,

$$\frac{d[\mathrm{AA}]}{dt} = k_{\mathrm{on1}}([\mathrm{TA}] - 2[\mathrm{AA}] - [\mathrm{AB}])^2 - k_{\mathrm{off1}}[\mathrm{AA}] \tag{39}$$

$$\frac{d[\mathrm{BB}]}{dt} = k_{\mathrm{on2}}([\mathrm{TB}] - 2[\mathrm{BB}] - [\mathrm{AB}])^2 - k_{\mathrm{off2}}[\mathrm{BB}] \quad (40)$$

$$\frac{d[\mathrm{AB}]}{dt} = k_{\mathrm{on3}}([\mathrm{TA}] - 2[\mathrm{AA}] - [\mathrm{AB}])([\mathrm{TB}] - 2[\mathrm{BB}] - [\mathrm{AB}]) - k_{\mathrm{off3}}[\mathrm{AB}]. \quad (41)$$

Under the equilibrated conditions, all the left sides are 0. In addition, converting the densities to the numbers,

$$0 = k_{\mathrm{on1}}(N_{\mathrm{0TA}} - 2N_{\mathrm{0AA}} - N_{\mathrm{0AB}})^2 - k_{\mathrm{off1}}N_{\mathrm{0AA}} \quad (42)$$

$$0 = k_{\mathrm{on2}}(N_{\mathrm{0TB}} - 2N_{\mathrm{0BB}} - N_{\mathrm{0AB}})^2 - k_{\mathrm{off2}}N_{\mathrm{0BB}} \quad (43)$$

$$0 = k_{\mathrm{on3}}(N_{\mathrm{0TA}} - 2N_{\mathrm{0AA}} - N_{\mathrm{0AB}})(N_{\mathrm{0TB}} - 2N_{\mathrm{0BB}} - N_{\mathrm{0AB}}) - k_{\mathrm{off3}}N_{\mathrm{0AB}}. \quad (44)$$

By solving these simultaneous equations, we can obtain the true numbers of dimers $N_{\mathrm{0AA}}$, $N_{\mathrm{0BB}}$, and $N_{\mathrm{0AB}}$.

Furthermore, in actual experiments, not all copies of the molecules can be fluorescently labeled, because the labeling efficiency tends to be limited to 60-90% under our typical experimental conditions (even when fluorescent proteins are used as probes). Accordingly, the observed fluorescent spots in the image represent only the molecules actually labeled with fluorescent probes.

Considering all the monomeric and dimeric states of the molecules, neglecting oligomers greater than dimers (certain molecules have special structures forming greater oligomers, but we do not consider them here), and accounting for the possible combinations of labeled and non-labeled molecules, all the molecules in the PM are classified into 14 cases. They are listed in **Supplementary Note 1-Table 1**.

Referring to the labeling efficiencies of protein molecules A and B as $E_{\mathrm{A}}$ and $E_{\mathrm{B}}$, respectively, and the true numbers of homodimers of AA and BB in the ROI as $N_{\mathrm{0AA}}$ and $N_{\mathrm{0BB}}$, respectively, the number of molecules that belong to each case (among 14 cases) can be expressed as shown in this Table. Note that AA (BB) looks like A (B) because we do not measure the signal intensities of the fluorescent spots. Considering this, the table includes how the molecules in each case appear (Appearance of the spot).

**Supplementary Note 1-Table 1**

| Case No. | Molecular composition | # of dye molecules | Number of molecules | Appearance of the spot |
|---|---|---|---|---|
| 1 | A | 1 | $E_{\mathrm{A}}N_{\mathrm{0A}}$ | A |
| 2 | A | 0 | $(1 - E_{\mathrm{A}})N_{\mathrm{0A}}$ | Invisible |
| 3 | B | 1 | $E_{\mathrm{B}}N_{\mathrm{0B}}$ | B |
| 4 | B | 0 | $(1 - E_{\mathrm{B}})N_{\mathrm{0B}}$ | Invisible |
| 5 | AA | 2 | ${E_{\mathrm{A}}}^2N_{\mathrm{0AA}}$ | A |
| 6 | AA | 1 | $2E_{\mathrm{A}}(1 - E_{\mathrm{A}})N_{\mathrm{0AA}}$ | A |
| 7 | AA | 0 | $(1 - E_{\mathrm{A}})^2N_{\mathrm{0AA}}$ | Invisible |
| 8 | BB | 2 | ${E_{\mathrm{B}}}^2N_{\mathrm{0BB}}$ | B |
| 9 | BB | 1 | $2E_{\mathrm{B}}(1 - E_{\mathrm{B}})N_{\mathrm{0BB}}$ | B |
| 10 | BB | 0 | $(1 - E_{\mathrm{B}})^2N_{\mathrm{0BB}}$ | Invisible |
| 11 | AB | 2 | $E_{\mathrm{A}}E_{\mathrm{B}}N_{\mathrm{0AB}}$ | AB |
| 12 | AB | 1 to A | $E_{\mathrm{A}}(1 - E_{\mathrm{B}})N_{\mathrm{0AB}}$ | A |
| 13 | AB | 1 to B | $E_{\mathrm{B}}(1 - E_{\mathrm{A}})N_{\mathrm{0AB}}$ | B |

| | | | | |
|---|---|---|---|---|
| 14 | AB | 0 | $(1-E_{\mathrm{A}})(1-E_{\mathrm{B}})N_{0\mathrm{AB}}$ | Invisible |

For example, in case 11, both molecules A and B in the heterodimer AB are labeled and contribute to the counts for $sN_{\mathrm{TA}}$, $sN_{\mathrm{TB}}$, and PCCF, whereas in the cases of 12 and 13, only one of the molecules in the heterodimer AB is labeled and thus it appears like monomers A and B, respectively. In case 14, neither of the molecules A and B in the heterodimer AB is labeled and thus this heterodimer makes no direct contributions to $sN_{\mathrm{TA}}$, $sN_{\mathrm{TB}}$, and PCCF. Another interesting problem occurs in cases 5 and 8. Here, both molecules A and B are labeled, and thus homodimers AA and BB are visible, but since we do not measure the intensities of the spots, they are indistinguishable from monomers A and B, respectively (i.e., they appear like monomers).

In the experiments, we count the numbers of fluorescent spots in the images in the A- and B-channels (again, note that we do not measure the intensities of individual spots here). Therefore, in the images of the A- and B-channels, we detect the following molecular spots.

[A-channel image]

Group 1 (called *apparent-monomer* group A): monomer A (case 1), homodimer AA (both labeled, case 5), homodimer AA (only one of the AA is labeled, case 6), and heterodimer AB with non-labeled B (case 12). The number of *apparent monomers* A counted in the ROI is written as $N_{\mathrm{A}}^{\mathrm{App}}$. Here, the word "apparent" is used with two meanings: (1) Apparent monomers A include AA spots and AB spots without labeled B in addition to the real monomer A, and (2) Due to incidental colocalizations, $N_{\mathrm{A}}^{\mathrm{App}}$ does not represent the number of molecules A found in the image in the A-channel, which is obtained after subtracting the number of incidentally overlapped spots. The same applies to all other apparent groups.

Group 2 (called *apparent-heterodimer* group): heterodimer AB with labeled molecules A and B (case 11). The number of *apparent heterodimers* AB counted in the ROI is written as $N_{\mathrm{AB}}^{\mathrm{App}}$.

[B-channel image]

Group 1 (called *apparent-monomer* group B): monomer B (case 3), homodimer BB (both labeled, case 8), homodimer BB (only one of the BB is labeled; case 9), and heterodimer AB with non-labeled A (case 13). The number of *apparent monomers* B counted in the ROI is written as $N_{\mathrm{B}}^{\mathrm{App}}$.

Group 2 (called *apparent-heterodimer* group): heterodimer AB with labeled molecules A and B (case 11). This is the same as Group 2 of the A-channel image.

Since we will determine the number of hypothetically existing heterodimers $N_{\mathrm{AB}}$, from the experiments, using **Eq. 28** ("Hypothetical" because although these dimers should exist, we cannot directly count the numbers of heterodimers in the images. This number is determined from the PCCF, according to **Eq. 28**.), in the following part of the theory, we will need to treat $N_{\mathrm{AB}}$ separately from the numbers of other A and B spots. Therefore, we have the grouping of Groups 1 and 2 as described in the previous paragraphs.

Based on the groups described above, $N_{\mathrm{A}}^{\mathrm{App}}$, $N_{\mathrm{B}}^{\mathrm{App}}$, and $N_{\mathrm{AB}}^{\mathrm{App}}$ can be expressed in the following way ($N$ with the subscripts of 1, 2, … indicate the number of molecules in cases 1, 2, …., respectively).

$$\begin{aligned} N_{\mathrm{A}}^{\mathrm{App}} &= N_1 + N_5 + N_6 + N_{12} \\ &= E_{\mathrm{A}}(N_{0\mathrm{A}} + (2-E_{\mathrm{A}})N_{0\mathrm{AA}} + (1-E_{\mathrm{B}})N_{0\mathrm{AB}}) \ , \end{aligned} \tag{45}$$

$$N_{\mathrm{B}}^{\mathrm{App}} = N_3 + N_8 + N_9 + N_{13}$$
$$= E_{\mathrm{B}}(N_{0\mathrm{B}} + (2 - E_{\mathrm{B}})N_{0\mathrm{BB}} + (1 - E_{\mathrm{A}})N_{0\mathrm{AB}}) \quad , \tag{46}$$

$$N_{\mathrm{AB}}^{\mathrm{App}} = N_{11}$$
$$= E_{\mathrm{A}}E_{\mathrm{B}}N_{0\mathrm{AB}} \quad . \tag{47}$$

**(3-2) Effect of incidental colocalizations**

$N_{\mathrm{A}}^{\mathrm{App}}$, $N_{\mathrm{B}}^{\mathrm{App}}$, and $N_{\mathrm{AB}}^{\mathrm{App}}$ cannot be directly linked to the number of spots A and B observed in the image or the $N_{\mathrm{AB}}$ determined from PCCF. This is because some of these *apparent monomers* A (B) and *apparent heterodimers* AB could not be separable as fluorescent spots in the image due to the diffraction-limited spatial resolutions (~200 nm), which is called "incidental colocalization". For simplicity, we assume that we cannot differentiate the same two molecules when they are located within a threshold distance $R_{\mathrm{Th}}$, where $R_{\mathrm{Th}}$ is much smaller than $\sqrt{A}$ ($A$: ROI area size). We also assume that $R_{\mathrm{Th}}$ is the same for monomers A (B) and homodimers AA (BB) (in principle, the localization accuracy of AA is better than that of A when both A's in AA are labeled, due to higher signal intensities).

To apply the correction for incidental colocalizations of apparent monomers A and B, and apparent heterodimers AB, we define the total numbers of apparent molecules A and B in the ROI, as **in Eqs. 18 and 19**, as $N_{\mathrm{TA}}^{\mathrm{App}}$ and $N_{\mathrm{TB}}^{\mathrm{App}}$, respectively. Namely,

$$N_{\mathrm{TA}}^{\mathrm{App}} = N_{\mathrm{A}}^{\mathrm{App}} + N_{\mathrm{AB}}^{\mathrm{App}} \quad , \tag{48}$$

$$N_{\mathrm{TB}}^{\mathrm{App}} = N_{\mathrm{B}}^{\mathrm{App}} + N_{\mathrm{AB}}^{\mathrm{App}} \quad . \tag{49}$$

$N_{\mathrm{TA}}^{\mathrm{App}}$ and $N_{\mathrm{TB}}^{\mathrm{App}}$ equal the numbers of fluorescent spots in the A- and B-channel images, $sN_{\mathrm{TA}}$ and $sN_{\mathrm{TB}}$, respectively, only when incidental colocalizations do not occur; i.e., when the experiments are performed with very low number densities of molecules A and B. For the general cases, $N_{\mathrm{TA}}^{\mathrm{App}}$ and $N_{\mathrm{TB}}^{\mathrm{App}}$ are greater than $sN_{\mathrm{TA}}$ and $sN_{\mathrm{TB}}$.

The following five cases of two apparent monomers and heterodimers located within the threshold distance $R_{\mathrm{Th}}$ should be considered.

[Incidental TA-TA]: incidental colocalizations of (apparent monomers A + apparent heterodimers AB) vs. (apparent monomers A + apparent heterodimers AB), affecting the counts in the A-channel image: represented as $N_{\mathrm{App(TA-TA)}}^{\mathrm{Incid}}$,

[Incidental TB-TB]: incidental colocalizations of (apparent monomers B + apparent heterodimers AB) vs. (apparent monomers B + apparent heterodimers AB), affecting the counts in the B-channel image: represented as $N_{\mathrm{App(TB-TB)}}^{\mathrm{Incid}}$,

[Incidental AB-AB]: incidental colocalizations of two apparent heterodimers AB, affecting the counts in both the A- and B-channel images: represented as $N_{\mathrm{App(AB-AB)}}^{\mathrm{Incid}}$,

[Incidental A-AB]: incidental colocalizations of an apparent monomer A and an apparent heterodimer AB, affecting the counts in the A-channel image: represented as $N_{\mathrm{App(A-AB)}}^{\mathrm{Incid}}$, and

[Incidental B-AB]: incidental colocalizations of an apparent monomer B and an apparent heterodimer AB, affecting the counts in the B-channel image: represented as $N_{\mathrm{App(B-AB)}}^{\mathrm{Incid}}$.

Since apparent monomers A and their counts $N_{\mathrm{A}}^{\mathrm{App}}$ (B and $N_{\mathrm{B}}^{\mathrm{App}}$) include both monomers A (B) and homodimers AA (BB) (**Eqs. 45 and 46**), we do not need to account for the incidental colocalizations of A and AA (B and BB).

Note that we only consider the colocalizations of two apparent molecules in each image A and B, and neglect the incidental colocalizations of more than two apparent molecules. Therefore, this theory is only applicable to the results obtained with lower number densities of molecules (see Supplementary Fig. 5).

The numbers of two incidentally colocalized apparent molecules are expressed in the following way.

$$N_{\mathrm{App(TA-TA)}}^{\mathrm{Incid}} = \frac{\pi {R_{\mathrm{Th}}}^2 N_{\mathrm{TA}}^{\mathrm{App}} N_{\mathrm{TA}}^{\mathrm{App}}}{2A} = \frac{\pi {R_{\mathrm{Th}}}^2 {N_{\mathrm{TA}}^{\mathrm{App}}}^2}{2A} \quad , \tag{50}$$

$$N_{\mathrm{App(TB-TB)}}^{\mathrm{Incid}} = \frac{\pi {R_{\mathrm{Th}}}^2 N_{\mathrm{TB}}^{\mathrm{App}} N_{\mathrm{TB}}^{\mathrm{App}}}{2A} = \frac{\pi {R_{\mathrm{Th}}}^2 {N_{\mathrm{TB}}^{\mathrm{App}}}^2}{2A} \quad , \tag{51}$$

$$N_{\mathrm{App(AB-AB)}}^{\mathrm{Incid}} = \frac{\pi {R_{\mathrm{Th}}}^2 N_{\mathrm{AB}}^{\mathrm{App}} N_{\mathrm{AB}}^{\mathrm{App}}}{2A} = \frac{\pi {R_{\mathrm{Th}}}^2 {N_{\mathrm{AB}}^{\mathrm{App}}}^2}{2A} \quad . \tag{52}$$

The factor of 2 in the denominator is to avoid doubly counting the same pairs.

$$N_{\mathrm{App(A-AB)}}^{\mathrm{Incid}} = \frac{\pi {R_{\mathrm{Th}}}^2 N_{\mathrm{A}}^{\mathrm{App}} N_{\mathrm{AB}}^{\mathrm{App}}}{A} \quad , \tag{53}$$

$$N_{\mathrm{App(B-AB)}}^{\mathrm{Incid}} = \frac{\pi {R_{\mathrm{Th}}}^2 N_{\mathrm{B}}^{\mathrm{App}} N_{\mathrm{AB}}^{\mathrm{App}}}{A} \quad . \tag{54}$$

The ROI should have the following number of AB pairs consisting of labeled A and labeled B molecules, $N_{\mathrm{AB}}$, although from the A- and B-channel images, we would be unable to determine which pairs represent true heterodimers and which are incidentally colocalized pairs.

$$\begin{aligned} N_{\mathrm{AB}} &= N_{\mathrm{AB}}^{\mathrm{App}} - N_{\mathrm{App(A-AB)}}^{\mathrm{Incid}} - N_{\mathrm{App(B-AB)}}^{\mathrm{Incid}} - N_{\mathrm{App(AB-AB)}}^{\mathrm{Incid}} \\ &= N_{\mathrm{AB}}^{\mathrm{App}} \cdot \left[1 - \frac{\pi {R_{\mathrm{Th}}}^2}{2A}\left(2N_{\mathrm{A}}^{\mathrm{App}} + 2N_{\mathrm{B}}^{\mathrm{App}} + N_{\mathrm{AB}}^{\mathrm{App}}\right)\right] \quad . \end{aligned} \tag{55}$$

Using **Eqs. 47-49**, **Eq. 55** can be rewritten as

$$N_{\mathrm{AB}} = E_{\mathrm{A}} E_{\mathrm{B}} N_{\mathrm{0AB}} \cdot \left[1 - \frac{\pi {R_{\mathrm{Th}}}^2}{2A}\left(2N_{\mathrm{TA}}^{\mathrm{App}} + 2N_{\mathrm{TB}}^{\mathrm{App}} - 3E_{\mathrm{A}} E_{\mathrm{B}} N_{\mathrm{0AB}}\right)\right] \quad . \tag{56}$$

Namely, based on this equation, the number of true heterodimers $N_{\mathrm{0AB}}$ could be evaluated by experimentally determining $N_{\mathrm{AB}}$ from the PCCF and $sN_{\mathrm{TA}}$ and $sN_{\mathrm{TB}}$ from the images.

$sN_{\mathrm{TA}}$ and $sN_{\mathrm{TB}}$ can be expressed as

$$sN_{\mathrm{TA}} = N_{\mathrm{TA}}^{\mathrm{App}} - N_{\mathrm{App(TA-TA)}}^{\mathrm{Incid}}$$
$$= N_{\mathrm{TA}}^{\mathrm{App}} - \frac{\pi {R_{\mathrm{Th}}}^2 {N_{\mathrm{TA}}^{\mathrm{App}}}^2}{2A} \quad , \tag{57}$$

$$sN_{\mathrm{TB}} = N_{\mathrm{TB}}^{\mathrm{App}} - N_{\mathrm{App(TB-TB)}}^{\mathrm{Incid}}$$
$$= N_{\mathrm{TB}}^{\mathrm{App}} - \frac{\pi {R_{\mathrm{Th}}}^2 {N_{\mathrm{TB}}^{\mathrm{App}}}^2}{2A} \quad . \tag{58}$$

These equations are solved to provide $N_{\mathrm{TA}}^{\mathrm{App}}$ and $N_{\mathrm{TB}}^{\mathrm{App}}$ using $sN_{\mathrm{TA}}$ and $sN_{\mathrm{TB}}$, which can be directly counted in the A- and B-channel images, respectively.

$$N_{\mathrm{TA}}^{\mathrm{App}} = \frac{A - A\sqrt{1 - 2\pi {R_{\mathrm{Th}}}^2 sN_{\mathrm{TA}}/A}}{\pi {R_{\mathrm{Th}}}^2} \quad . \tag{59}$$

$$N_{\mathrm{TB}}^{\mathrm{App}} = \frac{A - A\sqrt{1 - 2\pi {R_{\mathrm{Th}}}^2 sN_{\mathrm{TB}}/A}}{\pi {R_{\mathrm{Th}}}^2} \quad . \tag{60}$$

$N_{0\mathrm{A}}$ can be calculated using $N_{0\mathrm{AB}}$ by rewriting **Eq. 57** using **Eqs. 45, 47, and 48**,

$$sN_{\mathrm{TA}} = E_{\mathrm{A}} N_{0\mathrm{A}} + E_{\mathrm{A}}(2 - E_{\mathrm{A}}) N_{0\mathrm{AA}} + E_{\mathrm{A}} N_{0\mathrm{AB}} - N_{\mathrm{App(TA-TA)}}^{\mathrm{Incid}} \quad . \tag{61}$$

Likewise,

$$sN_{\mathrm{TB}} = E_{\mathrm{B}} N_{0\mathrm{B}} + E_{\mathrm{B}}(2 - E_{\mathrm{B}}) N_{0\mathrm{BB}} + E_{\mathrm{B}} N_{0\mathrm{AB}} - N_{\mathrm{App(TB-TB)}}^{\mathrm{Incid}} \quad . \tag{62}$$

The true total numbers of molecules ($N_{0\mathrm{TA}}$ and $N_{0\mathrm{TB}}$) are not changed (as in **Eqs. 37 and 38**).

$$N_{0\mathrm{TA}} = N_{0\mathrm{A}} + 2N_{0\mathrm{AA}} + N_{0\mathrm{AB}} \quad , \tag{63}$$

$$N_{0\mathrm{TB}} = N_{0\mathrm{B}} + 2N_{0\mathrm{BB}} + N_{0\mathrm{AB}} \quad . \tag{64}$$

Using these, **Eqs. 61 and 62** can be re-written as,

$$sN_{\mathrm{TA}} = E_{\mathrm{A}}(N_{0\mathrm{TA}} - 2N_{0\mathrm{AA}} - N_{0\mathrm{AB}}) + E_{\mathrm{A}}(2 - E_{\mathrm{A}}) N_{0\mathrm{AA}} + E_{\mathrm{A}} N_{0\mathrm{AB}} - N_{\mathrm{App(TA-TA)}}^{\mathrm{Incid}} \quad , \tag{65}$$

$$sN_{\mathrm{TB}} = E_{\mathrm{B}}(N_{0\mathrm{TB}} - 2N_{0\mathrm{BB}} - N_{0\mathrm{AB}}) + E_{\mathrm{B}}(2 - E_{\mathrm{B}}) N_{0\mathrm{BB}} + E_{\mathrm{B}} N_{0\mathrm{AB}} - N_{\mathrm{App(TB-TB)}}^{\mathrm{Incid}} \quad . \tag{66}$$

#### (4) Application to the analysis of hetero- and homo-dimerization using two-color movies

From here, we first consider the general case in which both hetero- and homo-dimerizations occur. Due to the presence of three equilibrium constants; i.e., those for heterodimers AB (${K_{\mathrm{D}}}^{\mathrm{AB}}$) and homodimers AA and BB (${K_{\mathrm{D}}}^{\mathrm{AA}}$ and ${K_{\mathrm{D}}}^{\mathrm{BB}}$, respectively), they could not be simultaneously evaluated. First, ${K_{\mathrm{D}}}^{\mathrm{AA}}$ and ${K_{\mathrm{D}}}^{\mathrm{BB}}$ must be determined, by separately performing experiments using one molecular species based on the theory described in this and the following sections. Then, by using these two homodimerization equilibrium constants, ${K_D}^{AB}$ could be determined by performing experiments for the mixed molecules A and B.

(i) By separately performing experiments, the rate constants for homodimerization ($k_{on1,2}$ and $k_{off1,2}$) and the off-rate of heterodimerization ($k_{off3}$) are determined.

(ii) From the observed numbers of A and B spots ($sN_{TA}$ and $sN_{TB}$), the apparent numbers of A and B spots ($N_{TA}^{App}$ and $N_{TB}^{App}$) can be calculated from **Eqs. 59 and 60**.

(iii) Using these results and the observed numbers of AB spots ($N_{AB}$), the true number of heterodimer AB ($N_{0AB}$) can be calculated from **Eq. 56.**

(iv) Using **Eqs. 50 and 51**, and the results obtained in step (ii), $N_{App(TA-TA)}^{Incid}$ and $N_{App(TB-TB)}^{Incid}$ can be calculated. Then, by combining $N_{App(TA-TA)}^{Incid}$, $N_{App(TB-TB)}^{Incid}$, $N_{0AB}$ (calculated in step (iii)), and the relationship between **Eqs. 42-44, 65 and 66**, these simultaneous equations are solved against $k_{on3}$.

(v) Finally, using the obtained $k_{on3}$ and $k_{off3}$, hetero$-K_D$ can be calculated as,

$$\text{hetero}-K_D = \frac{k_{on3}}{k_{off3}} \quad . \tag{67}$$

Next, we consider the case in which the homodimerization of one molecular species is investigated using two different probes with different colors (1 probe/1 protein copy), as done in the present research. Consider the two-color experiments for determining the homodimerization equilibrium constant. As explained, even in the case of studying homodimerization, including the two different probes used to label the protein, we can consider that two different molecular species A and B exist (although our target protein is just one molecular species). Meanwhile, we assume that the probes do not influence the homodimerization of the target protein molecules.

Under these conditions, all three off-rates $k_{off1,2,3}$ are the same as the true off-rate of homodimerization $k_{off}$ in **Eqs. 42-44**. However, since the formation probability of the AB dimers is twice that of the AA and BB dimers, $k_{on3}$ is twice as great as $k_{on1,2}$. Namely, the true on-rate of homodimerization $k_{on}$ can be expressed as,

$$k_{off} = k_{off1} = k_{off2} = k_{off3} \quad , \tag{68}$$

$$k_{on} = k_{on1} = k_{on2} = 2k_{on3} \quad . \tag{69}$$

Thus, the ratio of the heterodimer and the homodimers under the equilibrated conditions can be calculated with **Eqs. 42-44**.

$$N_{0AA} = N_{0BB} = 2N_{0AB} \quad . \tag{70}$$

This argument has been previously discussed.[14] Briefly, if *N* molecules of two chemical species exist in the system, and they are distinguishable, then the probability of collision between them is proportional to the number of possible combinations $N^2$. However, if they are un-distinguishable, the number of possible combinations becomes $N(N-1)/2$. If *N* is large enough, this can be approximated to $N^2/2$. Since the probability of collision is simply proportional to the on-rate, the on-rate of the distinguishable pair AB is twice those of the un-distinguishable pairs AA and BB.

In other words, **Eq. 1** should be re-determined as,

$$\text{homo-}K_\mathrm{D} = \frac{[\text{Monomer}]^2}{[\text{Dimer}]} = \frac{(N_{0\mathrm{A}}+N_{0\mathrm{B}})^2}{(N_{0\mathrm{AB}}+N_{0\mathrm{AA}}+N_{0\mathrm{BB}})A} = \frac{(N_{0\mathrm{A}}+N_{0\mathrm{B}})^2}{(2N_{0\mathrm{AB}})A} = \frac{k_\mathrm{on}}{k_\mathrm{off}} \quad . \tag{71}$$

Consider the practical method to calculate the $\text{homo-}K_\mathrm{D}$ in the case which the homodimerization of one molecular species is investigated using two different probes with different colors. Step (i) is simplified to obtain the single $k_\mathrm{off}$ ($= k_\mathrm{off1} = k_\mathrm{off2} = k_\mathrm{off3}$). The additional relationship written in **Eqs. 68 and 69** is then added to the simultaneous equations in step (iv). Finally, $\text{homo-}K_\mathrm{D}$ can be calculated with the obtained $k_\mathrm{on}$ and $k_\mathrm{off}$.

$$\text{homo-}K_\mathrm{D} = \frac{k_\mathrm{on}}{k_\mathrm{off}} \quad . \tag{72}$$

#### (5) Estimation of the $K_\mathrm{D}$ value from PCCF (the distribution of pair-wise distances)

Here, we develop the equations to determine the dimer dissociation equilibrium constant from the experimentally obtained PCCF. Steps (i) ~ (ii) in section 4 are performed as described previously.

Step (iii) is done as follows. As described in **Eqs. 26-28**, the experimentally obtained PCCF is fit by a Gaussian function, and we obtain the amplitude of the function from **Eq. 26**,

$$C = \frac{2A \cdot N_\mathrm{AB}}{sN_\mathrm{T}^{\ 2} \cdot \pi w^2} \quad . \tag{73}$$

Using **Eq. 56**,

$$N_{0\mathrm{AB}} = \frac{-F_2 + \sqrt{F_2^{\ 2} - 4F_1F_3}}{2F_1} \quad , \tag{74}$$

where

$$F_1 = \frac{3\pi R_\mathrm{Th}^{\ 2} E_\mathrm{A}^{\ 2} E_\mathrm{B}^{\ 2}}{2A}, \quad F_2 = E_\mathrm{A}E_\mathrm{B}\left[1 - \frac{\pi R_\mathrm{Th}^{\ 2}}{A}\left(N_\mathrm{TA}^\mathrm{App} + N_\mathrm{TB}^\mathrm{App}\right)\right], \\ F_3 = -\frac{C\pi w^2 (sN_\mathrm{TA} + sN_\mathrm{TB})^2}{2A} \quad . \tag{75}$$

$N_{0\mathrm{AB}}$ can be experimentally evaluated by using these equations. $C$ and $w$ are obtained from the fitting result. $E$ (labeling efficiency) should be predetermined, and $R_\mathrm{Th}$ is the diffraction limit distance (~ 200 nm). $sN_\mathrm{TA}$ and $sN_\mathrm{TB}$ are the total numbers of observed molecules A and B. $N_\mathrm{TA}^\mathrm{App}$ and $N_\mathrm{TB}^\mathrm{App}$ can be calculated from **Eqs. 59 and 60**, using $sN_\mathrm{TA}$ and $sN_\mathrm{TB}$.

After performing the calculations written in steps (iv) and (v), we can obtain $\text{hetero-}K_\mathrm{D}$ or $\text{homo-}K_\mathrm{D}$.

#### (6) Comparison of the developed theory with the simulation results

To validate these mathematical expressions, we performed Monte-Carlo simulations. The results are shown in Supplementary Fig. 5.

The theoretical PCCFs were very well fitted to the simulated PCCFs (Supplementary Fig. 5a, b), indicating that our calculations were virtually correct. However, we found that the theoretical PCCF was slightly different from the simulated data in the higher density range (Supplementary Fig. 5b, bottom right). This might be caused by the miscalculation of incidental colocalizations. As we stated in section 3-2, incidental colocalizations of more than two apparent molecules were neglected, and this effect is more critical in the higher density range. Since the theoretical number of incidentally colocalized spots was divided by 2 to avoid doubly counting the same pairs (**Eqs. 50-52**), the number of incidentally colocalized spots was over-counted (*n*-mer should be divided by the factor of *n*). This over-counting leads to the over-estimation of $N_{0\mathrm{AB}}$ (**Eqs. 55 and 74**), the under-estimation of $N_{0\mathrm{A}}$ and $N_{0\mathrm{B}}$ (**Eqs. 63 and 64**), and the under-estimation of the $K_{\mathrm{D}}$ (**Eq. 1**). The Monte Carlo simulation showed that incidental colocalizations of more than two molecules rapidly increased in the range of greater ~2 copies/µm$^2$ (Supplementary Fig. 5e). Consistent with this result, the under-estimation of $K_{\mathrm{D}}$ was enhanced in the range of greater than ~2 spots/µm$^2$ (Supplementary Fig. 5f). However, in our experimental data range (0.5 ~ 1.5 spots/µm$^2$), the differences between the true $K_{\mathrm{D}}$ and the estimated values were less than 5%. Thus, we concluded that our method to estimate the $K_{\mathrm{D}}$ is sufficiently reliable in the density range of our experimental conditions.
